# Identification of 2-amino benzothiazoles with bactericidal activity against *Mycobacterium tuberculosis*

**DOI:** 10.1101/2022.09.23.509298

**Authors:** Shilah Bonnett, Jo-Ann Jee, Somsundaram Chettiar, Yulia Ovechkina, Aaron Korkegian, Eric Greve, Joshua Odingo, Tanya Parish

## Abstract

We identified an amino-benzothiazole scaffold from a whole cell screen against recombinant *Mycobacterium tuberculosis* under expressing the essential signal peptidase LepB. The seed molecule had two-fold higher activity against the LepB hypomorph. Through a combination of purchase and chemical synthesis we explored the structure activity relationship for this series; 34 analogs were tested for anti-tubercular activity and for cytotoxicity against eukaryotic cells. We identified molecules with improved potency and reduced cytotoxicity. However, molecules did not appear to target LepB directly and did not inhibit protein secretion. Key compounds showed good permeability, low protein binding, and lack of CYP inhibition, but metabolic stability was poor with short half-lives. The seed molecule showed good bactericidal activity against both replicating and non-replicating bacteria, as well as potency against intracellular *M. tuberculosis* in murine macrophages. Overall, the microbiological properties of the series are attractive if metabolic stability can be improved, and identification of the target could assist in development of this series.

## INTRODUCTION

Tuberculosis remains a major public health threat with more than 1 million people dying annually and approximately millions of new cases per year ^1^. The COVID-19 global pandemic has considerably worsened the situation and may lead to increased cases of tuberculosis due to lack of diagnosis and treatment. Although there has been a large increase in the number of studies focused on identifying new tuberculosis drugs, the drug pipeline remains weak due to high attrition rates and the lack of new classes of drugs.

We are interested in identifying and progressing new molecular scaffolds with potential as anti-tubercular agents. Phenotypic screening using high throughput assays to identify agents from large molecular libraries has proved to be a good starting point with multiple new classes at the earliest stages of discovery ^2–7^. A number of screening approaches have been adopted including aerobic replicating growth in standard medium, varying carbon sources, nutrient and/or oxygen starvation, acidic conditions, and combinations of all these factors in multi stress models, as well as intracellular growth in macrophages and in multicellular granuloma models ^5,7–15^.

Biochemical screening to identify compounds with activity against *M. tuberculosis* has been largely unsuccessful due to their lack of whole cell activity ^16^. Alternative approaches to use hypomorph strains have been developed which enable the identification of additional chemical scaffolds (which may have weak activity against a wild-type strain) ^17,18^. The advantage of such an approach can be that target identification is more rapid. We are interested in the Sec pathway of protein secretion. Secretion via the Sec pathway is essential for *M. tuberculosis* viability and we previously confirmed that LepB itself is essential under standard growth conditions ^17^. We have previously developed and run a screen against a strain of *M. tuberculosis* which under expresses the essential signal peptidase LepB in an attempt to identify inhibitors of protein secretion via the Sec pathway ^17^. We report the exploration of a hit scaffold from this screen in this paper.

## RESULTS

### Identification of novel inhibitors of *M. tuberculosis* growth

We previously ran a high throughput screen against two strains of *M. tuberculosis* in order to find novel inhibitors of mycobacterial growth ^17,19^. Using a strain of *M. tuberculosis* which had lower expression of the signal peptidase LepB, we identified a single benzothiazole with modest anti-tubercular activity (Figure 1).

**Figure 1.**
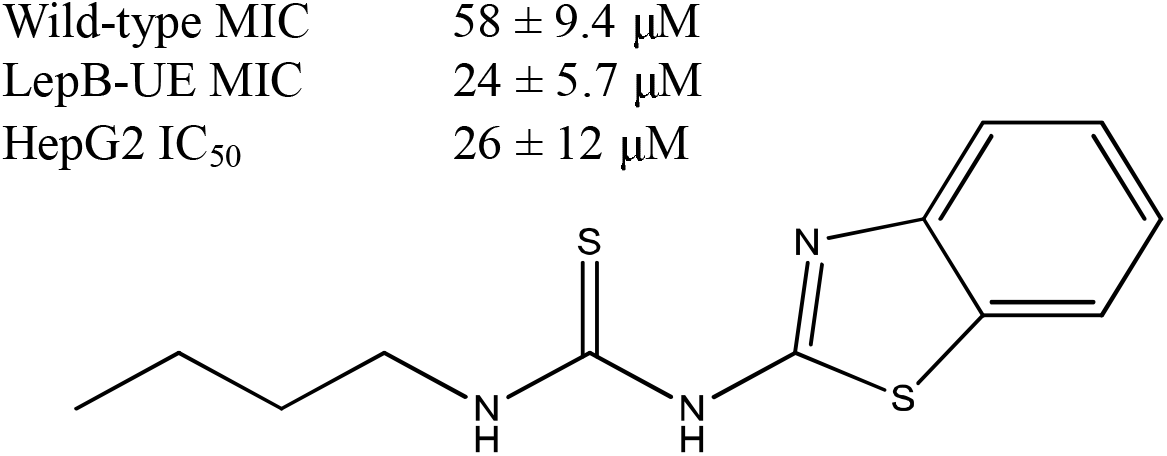
The seed benzothiazole molecule (1). The molecule was identified in a screen against *M. tuberculosis* under-expressing the sole signal peptidase. MICs were determined against this hypomorphic strain (LepB-UE) and the wild-type strain. Cytotoxicity was measured against HepG2 cells. Data are the average ± standard deviation (n = 4 for MICs).

This molecule had ∼two-fold higher activity against the LepB-UE strain (p<0.005). In order to determine if similar molecules had activity, we purchased a number of analogs and tested these for activity against the wild-type and LepB-UE strains (Figure 2 and Table 1). Only two additional molecules (**4** and **8**) had activity against the wild-type strain with modest MICs of 125 µM and 27 µM respectively. However, six of the molecules had reasonable activity against the LepB-UE, with five having MIC <100 µM ranging from 14 to 82 µM (molecules **1, 4, 5, 6** and **8**). We also measured cytotoxicity against the human HepG2 cell line. Nine of the molecules were cytotoxic, but the activity did not track with anti-tubercular activity suggesting that the two biological activities could be separated. Based on these data we decided to conduct hit assessment to determine the tractability of the series. We designed and synthesized a small set of analogs to explore the structure-activity relationship with a focus on improving potency against the wild-type strain and eliminating cytotoxicity.

**Figure 2.**
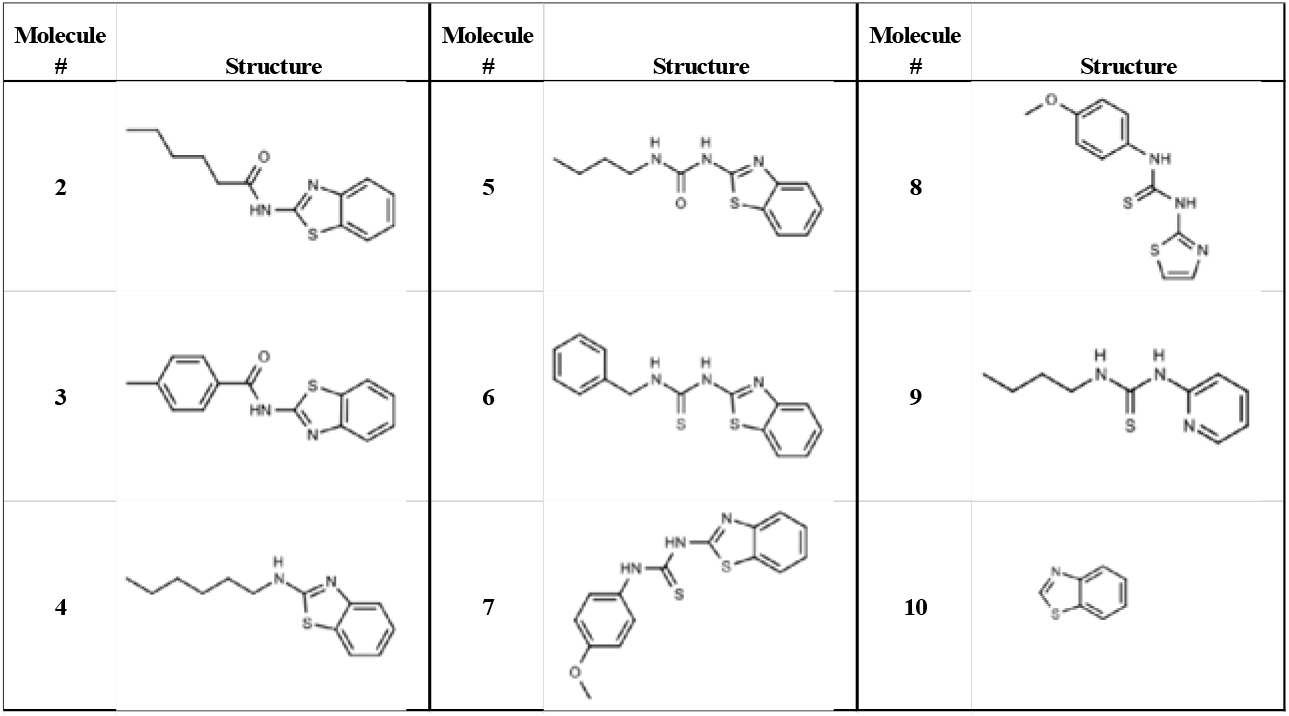
Molecules purchased for hit evaluation.

**Table 1.**
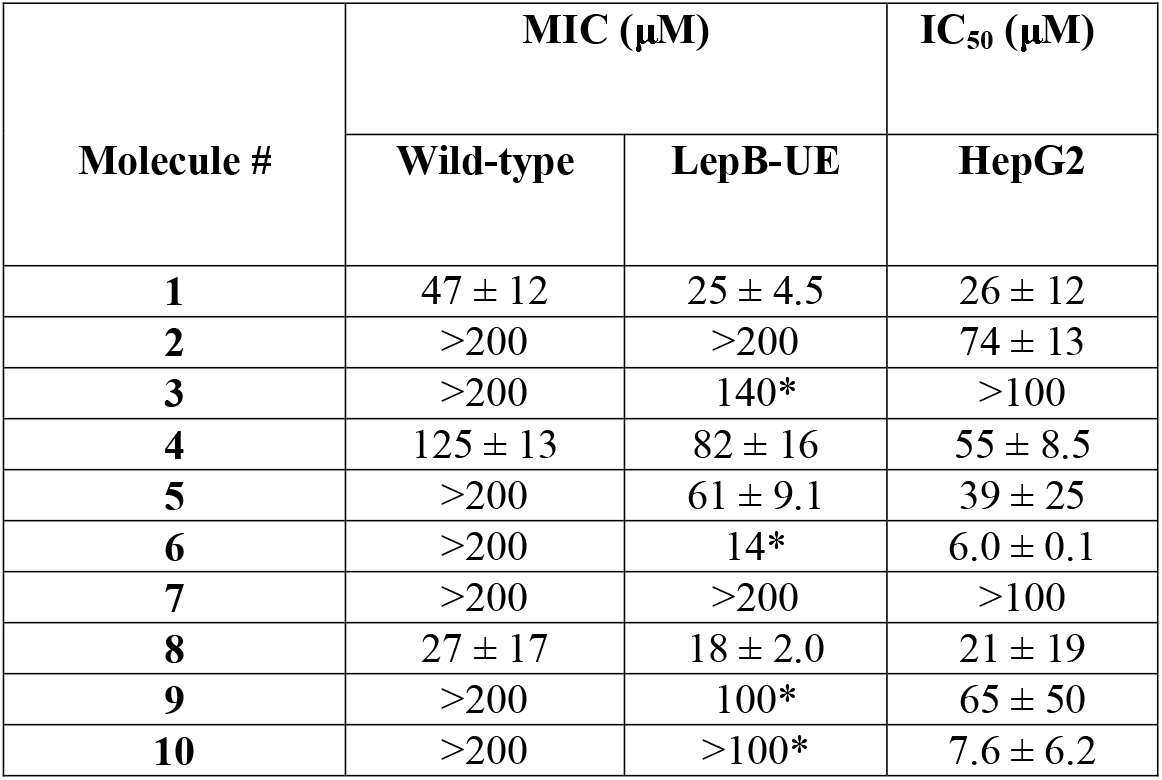
Activity of commercially available analogs. MIC = concentration required to inhibit growth by 90%. IC_50_ concentration required to inhibit growth by 50%. Data are average and standard deviation of a minimum of 2 biological replicates (except those marked with * from n=1).

| Molecule # | MIC (μM) |  | IC<sub>50</sub> (μM) |
| --- | --- | --- | --- |
|  | Wild-type | LepB-UE | HepG2 |
| 1 | 47 ± 12 | 25 ± 4.5 | 26 ± 12 |
| 2 | >200 | >200 | 74 ± 13 |
| 3 | >200 | 140* | >100 |
| 4 | 125 ± 13 | 82 ± 16 | 55 ± 8.5 |
| 5 | >200 | 61 ± 9.1 | 39 ± 25 |
| 6 | >200 | 14* | 6.0 ± 0.1 |
| 7 | >200 | >200 | >100 |
| 8 | 27 ± 17 | 18 ± 2.0 | 21 ± 19 |
| 9 | >200 | 100* | 65 ± 50 |
| 10 | >200 | >100* | 7.6 ± 6.2 |

### Structure activity relationship for the benzothiazole (BET) series

We first explored the benzothiazole core. In our first iteration, we synthesized three new molecules; all were tested for activity against the wild-type and LepB-UE strains, as well as for cytotoxicity for HepG2 cells (Table 2). Replacing the benzothiazole with a benzimidazole (**11**) or benzoxazole (**12**) retained good potency against the LepB-UE strain, while the benzoxazole led to improved activity against the wild-type strain that was equivalent to the LepB-UE strain (MIC of 23 - 32 µM), but both molecules were cytotoxic. Adding a chloro substituent (**13)** at position 4 increased activity against the LepB-UE strain (MIC = 7.9 µM), and reduced cytotoxicity (HepG2 IC_50_ >100 µM) but did not improve potency against the wild-type strain.

**Table 2.**
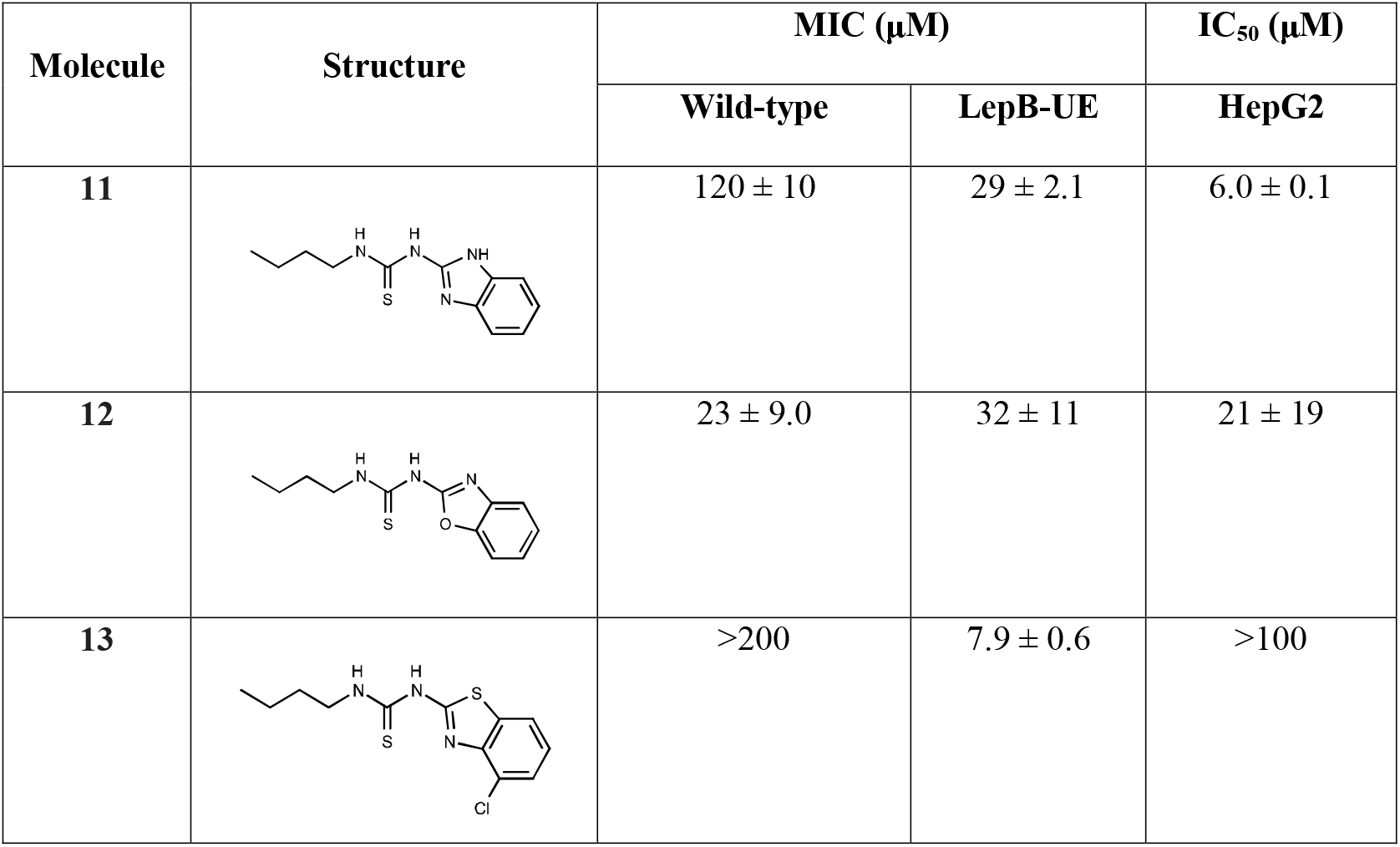
Activity of core analogs. MIC = concentration required to inhibit growth by 90%. IC_50_ concentration required to inhibit growth by 50%. Data are average and standard deviation of a minimum of 2 biological replicates.

| Molecule | Structure | MIC (μM) | | IC<sub>50</sub> (μM) |
| --- | --- | --- | --- | --- |
|  |  | Wild-type | LepB-UE | HepG2 |
| **11** | | 120 ± 10 | 29 ± 2.1 | 6.0 ± 0.1 |
| **12** | | 23 ± 9.0 | 32 ± 11 | 21 ± 19 |
| **13** | | >200 | 7.9 ± 0.6 | >100 |

Secondly, we simplified the benzothiazole to a single ring system. We synthesized 10 analogs and tested for biological activity (Table 3).

**Table 3.**
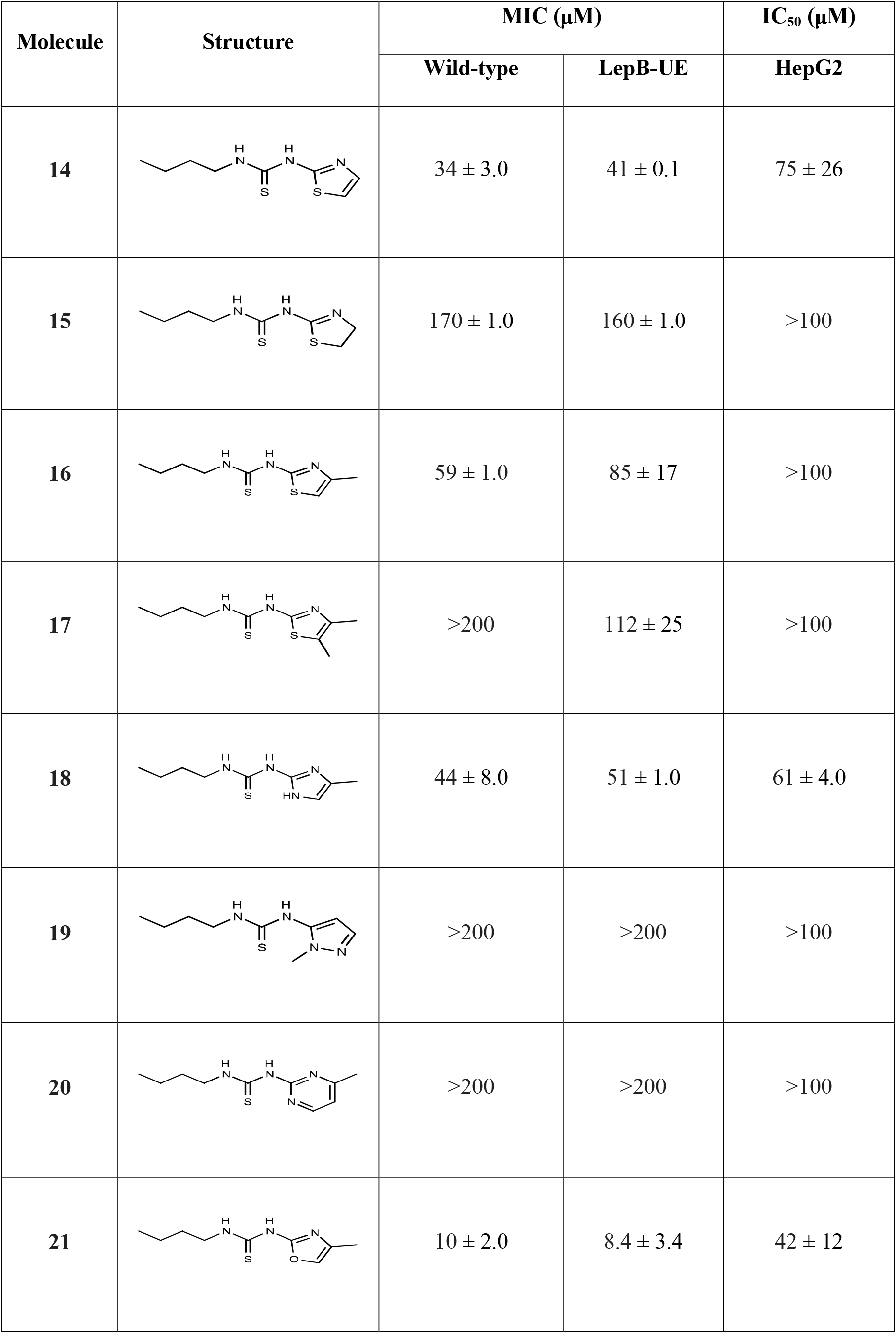
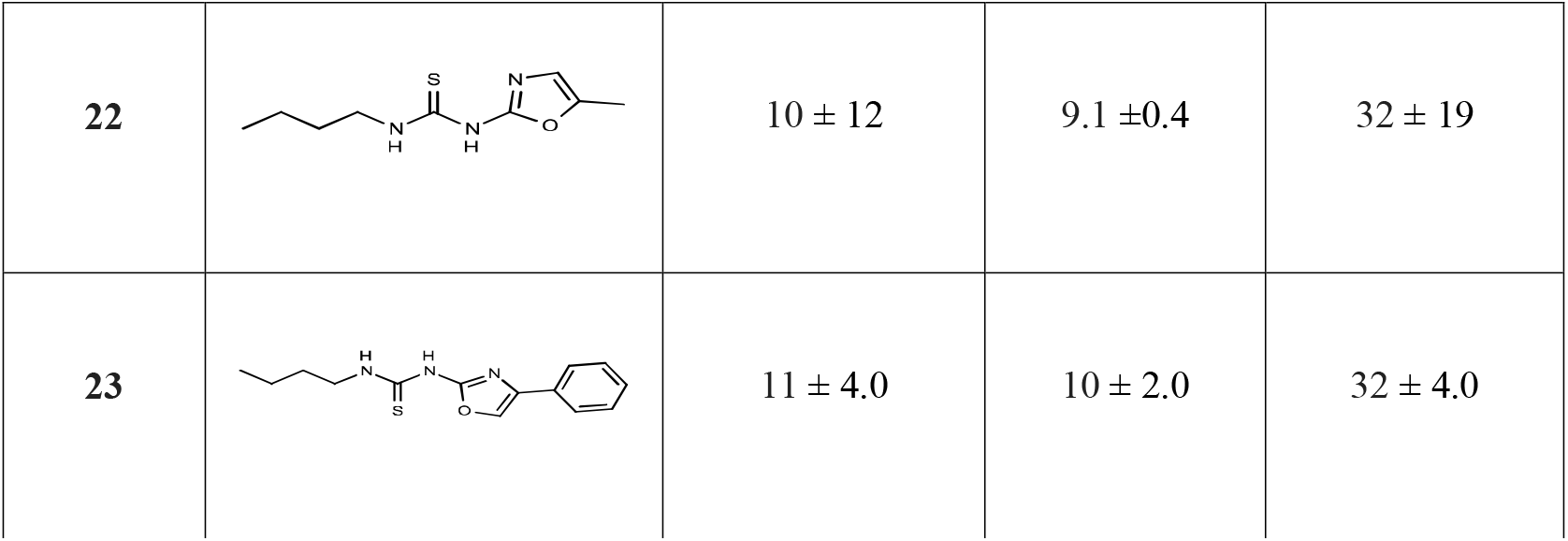
Activity of oxazole and thiazole core analogs. MIC = concentration required to inhibit growth by 90%. IC_50_ concentration required to inhibit growth by 50%. Data are average and standard deviation of a minimum of 2 biological replicates.

| Molecule | Structure | MIC ($\mu\text{M}$) | | IC<sub>50</sub> ($\mu\text{M}$) |
| --- | --- | --- | --- | --- |
|  |  | Wild-type | LepB-UE | HepG2 |
| 14 | | $34 \pm 3.0$ | $41 \pm 0.1$ | $75 \pm 26$ |
| 15 | | $170 \pm 1.0$ | $160 \pm 1.0$ | >100 |
| 16 | | $59 \pm 1.0$ | $85 \pm 17$ | >100 |
| 17 | | >200 | $112 \pm 25$ | >100 |
| 18 | | $44 \pm 8.0$ | $51 \pm 1.0$ | $61 \pm 4.0$ |
| 19 |  | >200 | >200 | >100 |
| 20 |  | >200 | >200 | >100 |
| 21 | | $10 \pm 2.0$ | $8.4 \pm 3.4$ | $42 \pm 12$ |
| **22** | | $10 \pm 12$ | $9.1 \pm 0.4$ | $32 \pm 19$ |
| **23** | | $11 \pm 4.0$ | $10 \pm 2.0$ | $32 \pm 4.0$ |

Compared to compound **1**, compound **14** with the 1,3 thiazole had improved potency against the wild-type strain (34 µM compared to 120 µM), but no increased activity against the LepB-UE strain (MICs of 29 µM and 41 µM). Removal of a double bond to give compound **15** resulted in a 4-5-fold loss of activity with no differential activity (MIC of 170 µM and 160 µM against wild-type and LepB-UE strains respectively). Adding a methyl group at position 5 (**16**) or 4,5 dimethyl group (**17**) did not improve activity. Overall, the thiazole derivatives were less cytotoxic than their benzothiazoles (HepG2 IC_50_>100 µM), with only **15** demonstrating any cytotoxicity (HepG2 IC_50_ = 75 µM). Replacing the sulphur (**17**) with nitrogen (**18**) retained activity against both strains (MIC = 44 µM and 51 µM for wild-type and LepB-UE respectively), but this molecule was cytotoxic (HepG2 IC_50_ = 61 µM). Replacing the thiazole with a methylated pyrazole (**19**) or methylated pyrimidine (**20**) resulted in loss of activity against both strains, but these molecules were not cytotoxic. Methylation of the oxazole at either position 4 or 5 (**21** and **22**) greatly improved activity against both strains (MICs of 10 µM against the wild-type; MICs of 8-9 µM LepB UE strains). Although both molecules had some cytotoxicity (HepG2 IC_50_ = 42 and 32 µM for **21** and **22** respectively), there was improved selectivity over earlier analogs. Addition of a phenyl group (**23**) had a similar effect with improved activity and some cytotoxicity (MICs 10-11 µM, HepG2 IC_50_ 32 µM).

Next, we explored the left-hand side of the molecule (Table 4). Inclusion of an oxygen or sulphur in the alkyl chain (**24** and **25**) resulted in some activity against the wild-type but resulted in increased cytotoxicity and loss of selectivity. Addition of a phenyl group (**26**) increased activity against the LepB-UE strain, but no activity against the wild-type strain. Addition of a methyl group on the thiourea (**27**) greatly increased cytotoxicity (HepG2 IC_50_ = 5.6 µM) without improving anti-tubercular activity (MIC = 155-195 µM). Replacing the alkyl chain with an isopropyl group resulted in good activity for both the benzothiazole (**28**) and oxazole derivatives (**29**), but no selectivity (MICs of 20-40 µM against wild-type; MICs of 12-21 µM against LepB-UE; HepG2 IC_50_ of 27-78 µM).

**Table 4.**
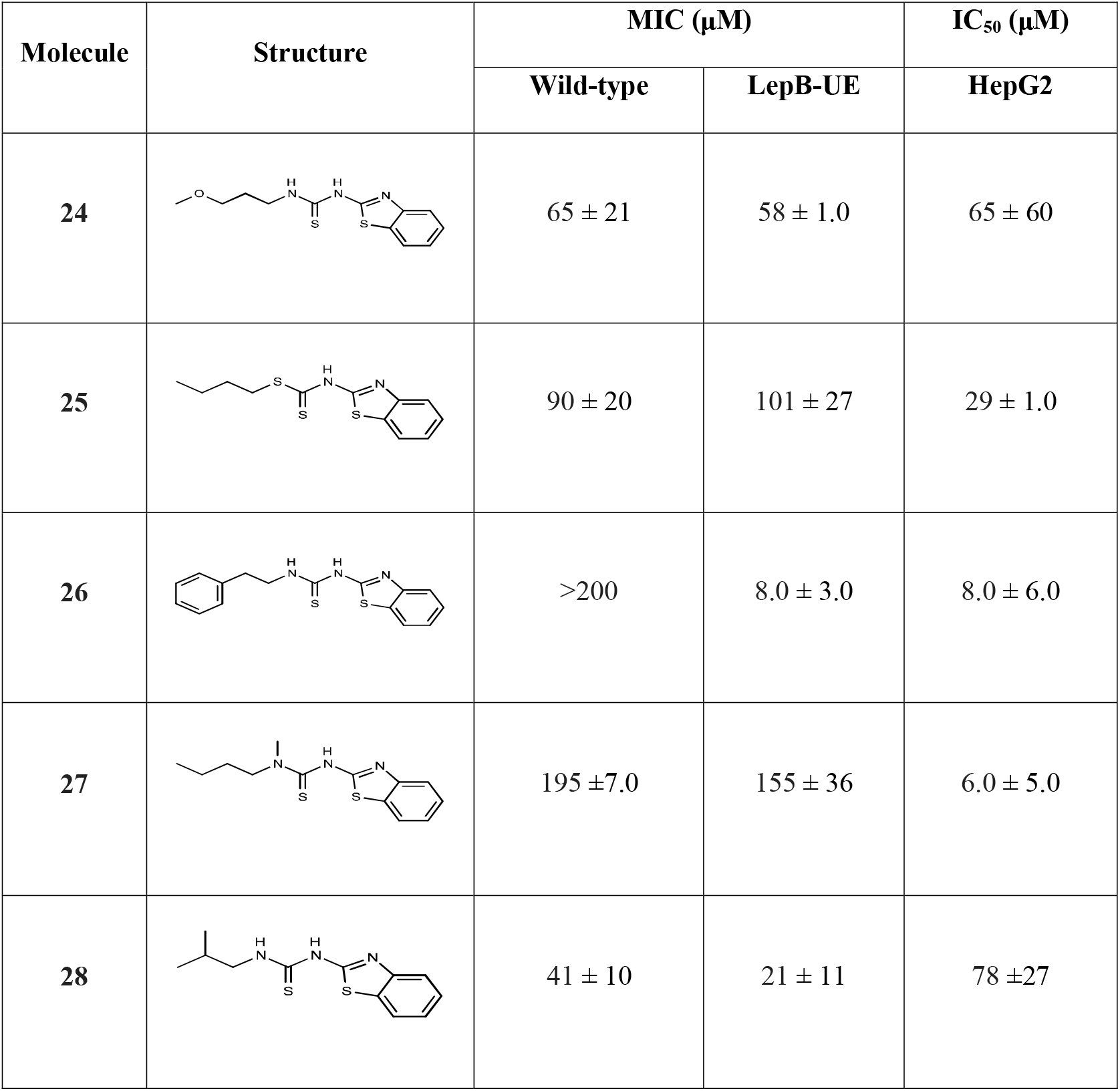
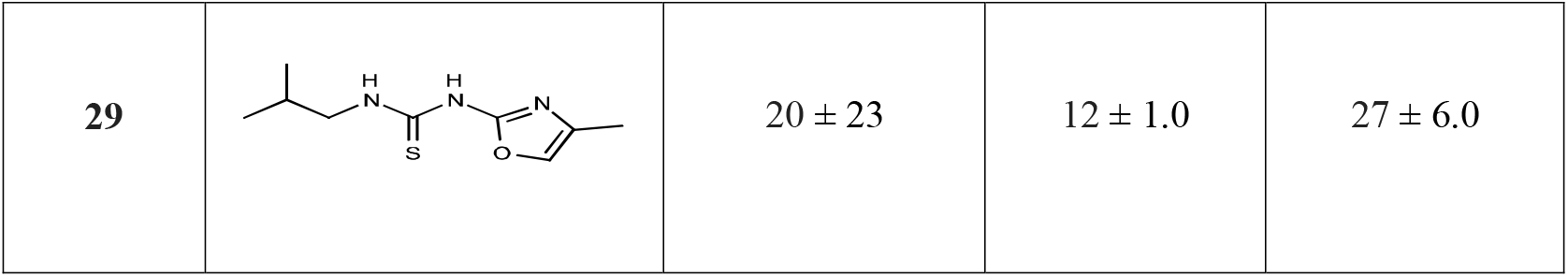
Activity of side chain core analogs. MIC = concentration required to inhibit growth by 90%. IC_50_ concentration required to inhibit growth by 50%. Data are average and standard deviation of a minimum of 2 biological replicates.

| Molecule | Structure | MIC (µM) |  | IC<sub>50</sub> (µM) |
| --- | --- | --- | --- | --- |
|  |  | Wild-type | LepB-UE | HepG2 |
| **24** |  | 65 ± 21 | 58 ± 1.0 | 65 ± 60 |
| **25** |  | 90 ± 20 | 101 ± 27 | 29 ± 1.0 |
| **26** |  | >200 | 8.0 ± 3.0 | 8.0 ± 6.0 |
| **27** |  | 195 ± 7.0 | 155 ± 36 | 6.0 ± 5.0 |
| **28** |  | 41 ± 10 | 21 ± 11 | 78 ± 27 |
| **29** | | $20 \pm 23$ | $12 \pm 1.0$ | $27 \pm 6.0$ |

Finally, we made further changes to the molecule on both the left- and right-hand sides. Switching the thiourea to the benzene ring from the thiazole (**30**) reduced cytotoxicity but lost ∼two-fold biological activity. Since we had seen an improvement in selectivity by adding a chloro-substituent in molecule **13**, we made another analog with chloro at the 4 position (**31**); this also abrogated cytotoxicity (HepG2 IC_50_ >100 µM), while retaining activity against the LepB-UE strain (MIC of 14 µM) but no activity against the wild-type strain. Since the oxazoles were generally more active, we synthesized benzoxazoles with chloro substituents on the 5-, 6- and 7-positions **(32**-**34**); however, this did not improve selectivity and reduced biological activity. Finally, we added a methoxy group to the benzoxazole (**35**), but this abrogated all activity, although the molecule was not toxic.

### BET molecules do not inhibit protein secretion

We originally identified the seed molecule for this series from a screen against the LepB-UE strain and confirmed that it had differential activity i.e. it was more effective against the the LepB-UE strain (Figure 1). However, our structure activity relationship studies did not see any increased activity against the LepB-UE strain (Figure 3). To determine if molecules target the secretory pathway, we tested a subset for inhibition of protein secretion via the Sec pathway in membrane fractions of *M. tuberculosis* using a fluorogenic peptide substrate (Table S1). Fourteen molecules representative of the series were tested, but none showed inhibition of secretion up to 200 µM, suggesting that they work via another mechanism.

**Figure 3.**
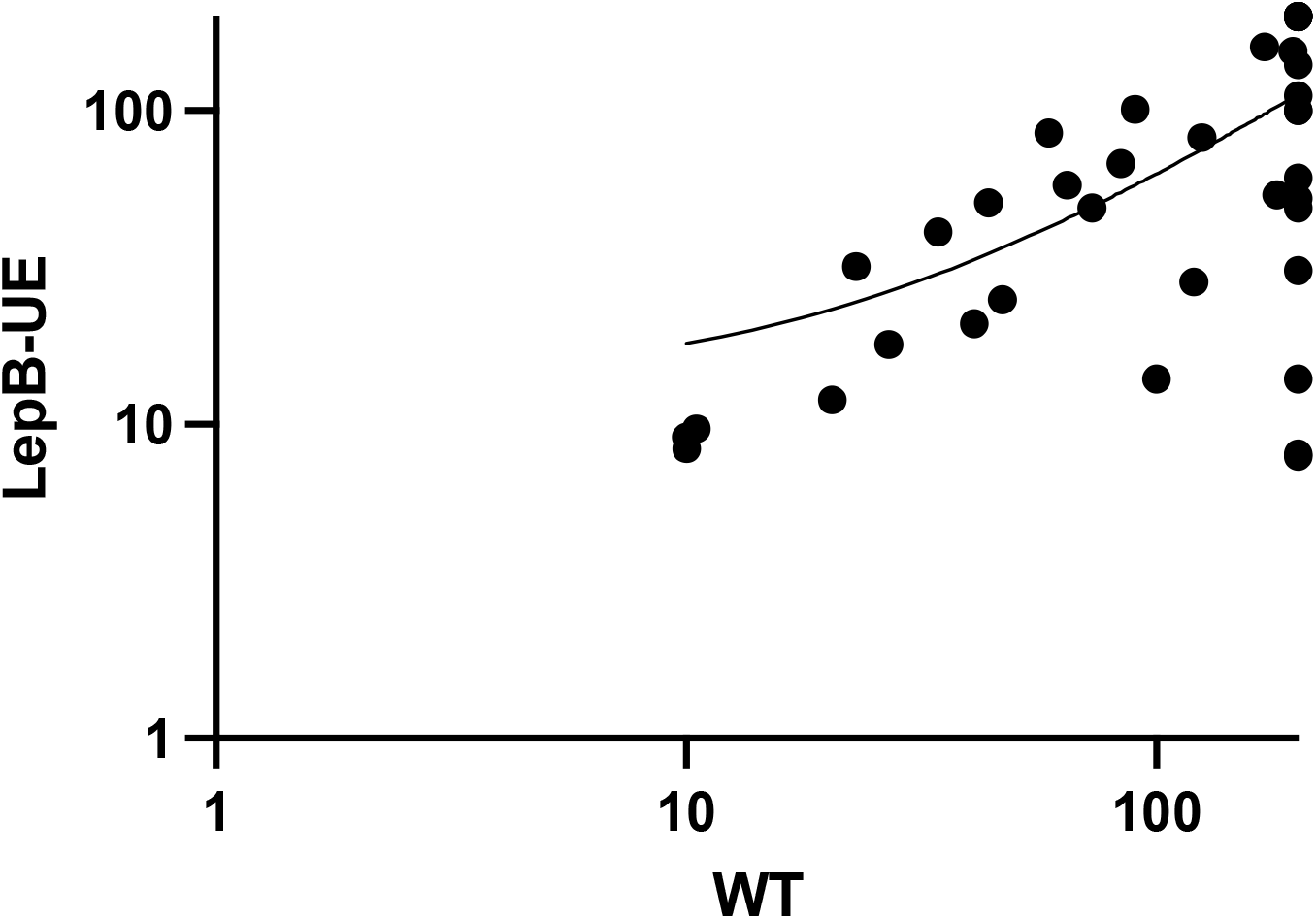
Correlation between activity against wild-type and LepB-UE strains. The average MICs for each strain are plotted. The Pearson correlation coefficient R^2^= 0.32. LepB-UE = *M. tuberculosis* strain under-expressing LepB. WT = *M. tuberculosis* wild-type strain.

### BET molecules are active against intracellular *M. tuberculosis*

We tested a small subset of molecules for activity against *M. tuberculosis* cultured in murine macrophage using a high content microscopy method. This enabled us to test cytotoxicity against infected macrophages simultaneously in order to reduce artefacts, since toxicity to the macrophages would also prevent mycobacterial growth (Table 6). Three molecules had activity against intracellular *M. tuberculosis*, two of these molecules (**1** and **23**) had excellent potency of < 10 µM. Interestingly, the activity against intracellular bacteria was far greater than that against aerobically cultured organisms. We were unable to measure intracellular activity for the remaining six molecules tested due to cytotoxicity against the murine macrophages. Activity against intracellular organisms is of particular interest since there is increasing evidence to link intracellular bacteria with both increased heterogeneity and antibiotic tolerance ^20,21^; thus targeting this population could be useful in a new regimen.

**Table 5.**
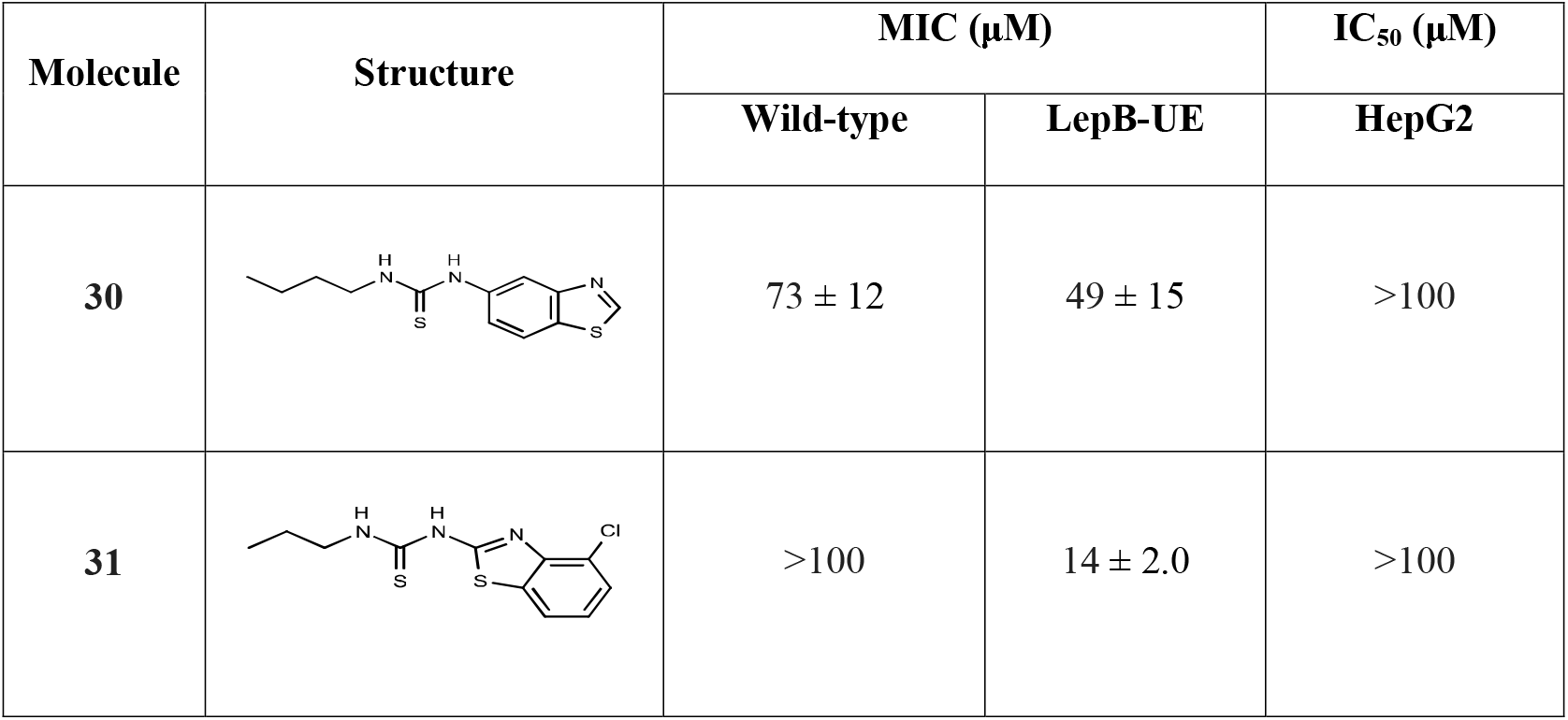
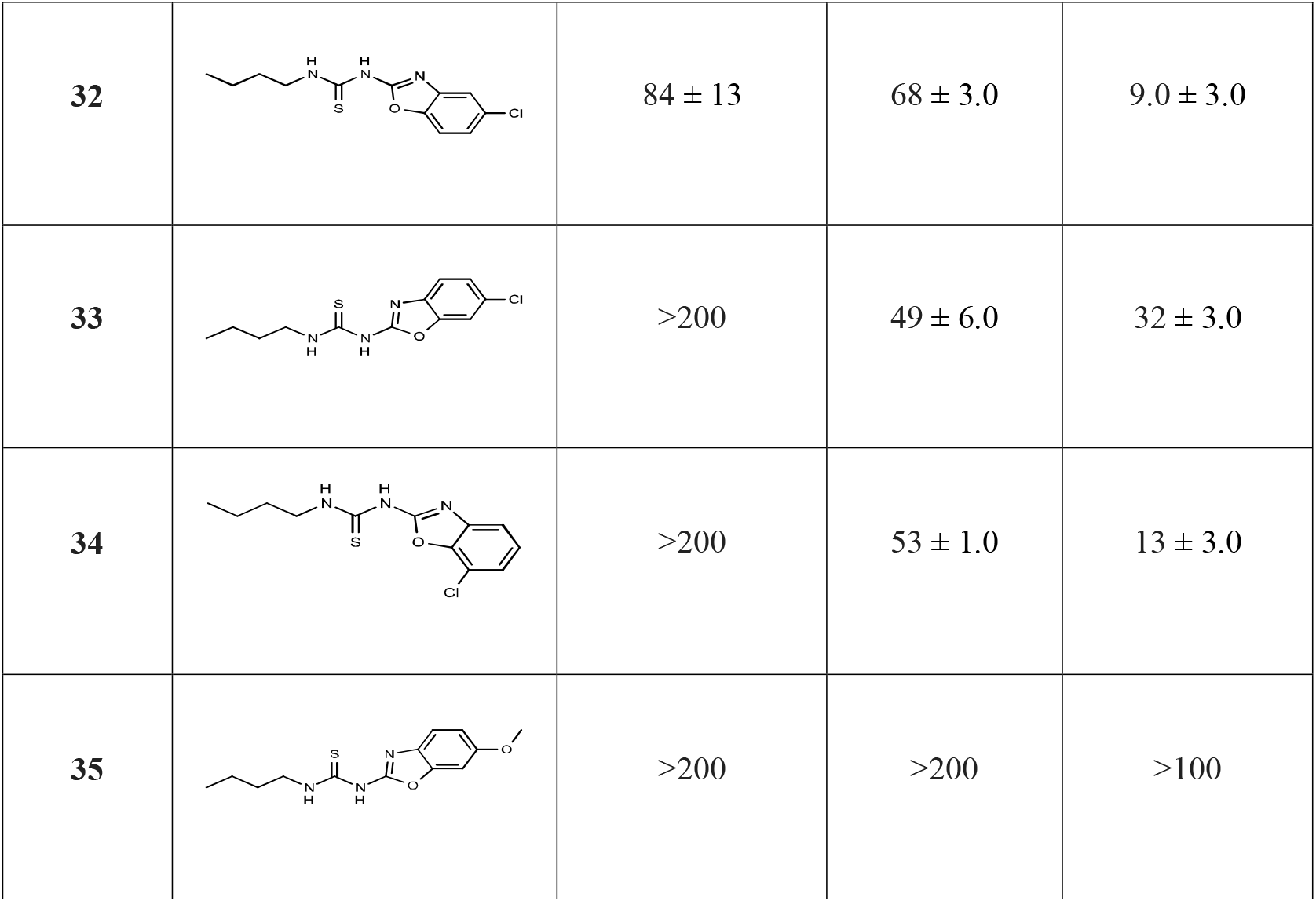
Activity of analogs. MIC = concentration required to inhibit growth by 90%. IC_50_ concentration required to inhibit growth by 50%. Data are average and standard deviation of a minimum of 2 biological replicates.

| Molecule | Structure | MIC (μM) | | IC<sub>50</sub> (μM) |
| --- | --- | --- | --- | --- |
|  |  | Wild-type | LepB-UE | HepG2 |
| **30** | | $73 \pm 12$ | $49 \pm 15$ | >100 |
| **31** | | >100 | $14 \pm 2.0$ | >100 |
| 32 | | $84 \pm 13$ | $68 \pm 3.0$ | $9.0 \pm 3.0$ |
| 33 | | >200 | $49 \pm 6.0$ | $32 \pm 3.0$ |
| 34 | | >200 | $53 \pm 1.0$ | $13 \pm 3.0$ |
| 35 |  | >200 | >200 | >100 |

**Table 6.**
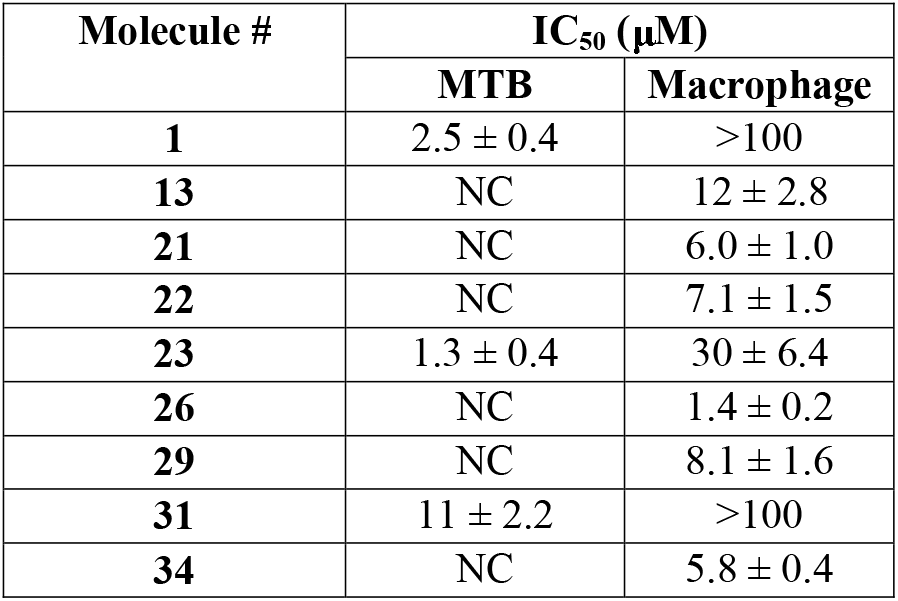
Activity against intracellular *M. tuberculosis*. IC_50_ is the concentration required to inhibit growth by 50% for either intracellular *M. tuberculosis* (MTB) or the RAW264:7 cells (macrophage). Data are average and standard deviation of a minimum of 2 biological replicates. NC = not calculated due to cytotoxicity against RAW264:7 cells.

### BET molecules have bactericidal activity against *M. tuberculosis*

We tested two molecules for their ability to kill *M. tuberculosis* under replicating (aerobic growth) and non-replicating (nutrient starvation) conditions (Figure 4). Both molecules were able to kill replicating organisms with complete sterilization of replicating cultures at 100 µM after 21 days. Compound **29** was more effective with bactericidal activity (defined as 3 log kill) at 27.5 µM. In contrast, compound **1** was more effective at killing non-replicating organisms with sterilization at the lowest concentration tested (20 µM), whereas compound **29** was only effective at the highest concentration (100 µM). This may indicate that the molecules have a different mode of action.

**Figure 4.**
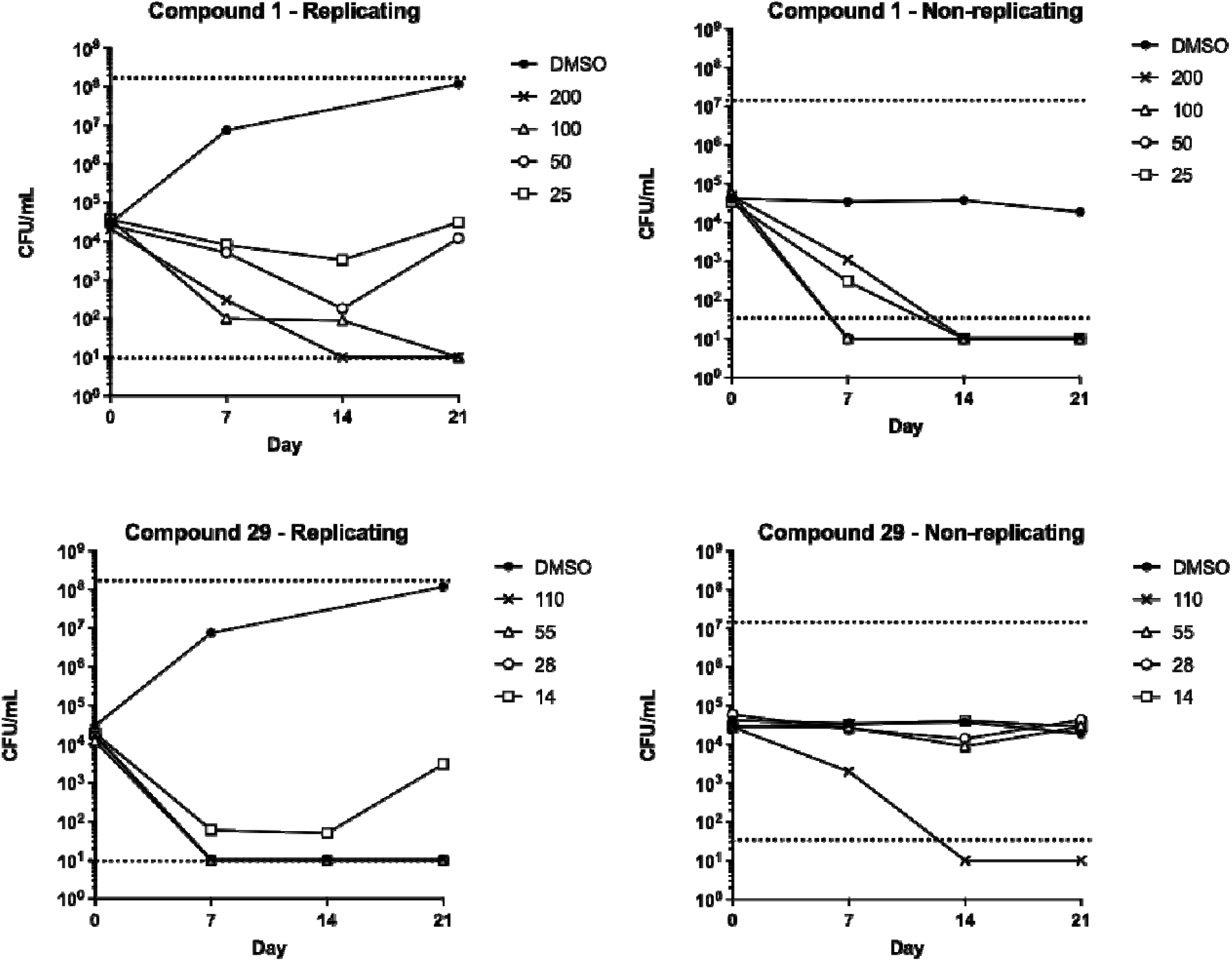
Kill kinetics for Compounds 1 and 29 against *M. tuberculosis.* Replicating bacteria were cultured in standard medium, non-replicating bacteria were in PBS-Tyloxapol. Compounds were added at the indicated concentrations (*μ*M). Viable bacteria were measured as colony forming units (CFU).

### ADME properties

We selected five compounds including the original seed compound for *in vitro* ADME studies. We prioritized compounds with the lowest cytotoxicity, while selecting compounds with different structural features. Three molecules showed good permeability determined using Caco-2 cells, and none of the five were subject to efflux (Table 8). Microsomal stability was generally low with half-life in the presence of human microsomes ranging from 8 to 26 minutes (Table 7 and Table S2). The molecules were generally clean in cytochrome P540 inhibition assays (Table S3), except that three molecules inhibited CYP2C19. Protein binding was variable ranging from 94.5% for compound **26** to 99.9% for compound **1** (Table S4). These data suggest that the key liability for the series is metabolic stability.

**Table 7.** Metabolic stability. NADPH-dependent clearance (CL_int_) and half-life (T_1/2_) was measured using human liver microsomes with test compounds at 1 µM. Verapamil and dextromethorphan were used as controls for high and low clearance respectively. CLint = ln(2) /(T1/2 [microsomal protein]). Data are average of two replicates.

| Molecule # |  |  |
| --- | --- | --- |
|  | CL<sub>int</sub><br>(μL min<sup>-1</sup><br>mg<sup>-1</sup>) | T<sub>1/2</sub><br>(min) |
| **1** | 96 | 24 |
| **12** | 218 | 11 |
| **14** | 89 | 26 |
| **21** | 179 | 13 |
| **24** | 282 | 8.2 |
| **Verapamil** | 128 | 18 |
| **Dextromethorphan** | 19.5 | 118 |

**Table 8.** Cell permeability. The permeability coefficient (Papp) was measured using Caco-2 cells. (A→B) is Apical to Basolateral; (B→A) is Basolateral to Apical. Efflux ratio (Re) was calculated as Papp(B→A)/Papp(A→B). Atenolol = low permeability control. Propanolol = high permeability control. Talinolol = P-glycoprotein efflux control. Data is expressed as permeability (10^−6^ cm s^-1^). Papp = (dQ/dt)/ C0*A. dQ/dt is rate of permeation, C0 is initial concentration of test agent, and A is the area of monolayer.

| Molecule # | Papp (10<sup>-6</sup> cm s<sup>-1</sup>) |  | Efflux ratio (Re) |
| --- | --- | --- | --- |
|  | mean A→B | mean B→A |  |
| 1 | 1.4 | 2.5 | 1.7 |
| 12 | 4.9 | 5.8 | 1.2 |
| 14 | 22 | 18 | 0.81 |
| 21 | 21 | 19 | 0.89 |
| 24 | 17 | 15 | 0.87 |
| Atenolol | 0.13 | 0.3 | 2.4 |
| Propanolol | 20 | 30 | 1.5 |
| Talinolol | 0.033 | 5.3 | 161 |

To our knowledge this is the first report of the activity and SAR against *M. tuberculosis* for benzothiazoles and benzoxazoles containing a urea or thiourea substituent, although similar molecules have been reported as active for different activities such as myorelaxants ^22^. Previous work has identified benzothiazole amide derivatives as active against both *M. tuberculosis* and non-tuberculous mycobacteria ^22,23^, and a single benzothiazole had weak activity against *M. tuberculosis* ^24^, but no urea derivatives were included, and the majority of the compounds were substituted on the cyclohexane ring. The related anti-tubercular thioalkyl benzoxazole is proposed to be metabolized by *M. tuberculosis* MymA to the active form ^25^. EthA and MymA are mono-oxygenases which can activate sulphur-containing prodrugs and it is possible that the molecules we describe are activated by the same mechanism ^26^. However, it is not clear why strains with lowered levels of the signal peptidase LepB would be more sensitive to prodrugs, and neither EthA nor MymA are secreted proteins. We determined that the molecules in this series did not work by inhibiting secretion in *M. tuberculosis*. There are other possibilities for the differential activity between the wild-type and LepB-UE strain. In particular, there could be changes in the cell wall which alter either molecule permeation and/or efflux thereby affecting the intracellular concentration.

We explored a series of molecules based around a benzothiazole seed molecule for their potential as new anti-tubercular agents. We conducted preliminary structure activity relationships and identified molecules with improved potency and reduced cytotoxicity, as well as molecules with excellent intracellular activity and selectivity. The key liability for the series is metabolic stability. The series does not appear to target protein secretion and further work to investigate the mode of action could assist in development of this series by allowing for structure-based design.

## ACKNOWLEDGEMENTS

We thank Lena Anoshchenko, Lina Castro, Lindsay Flint, James Johnson, Megha Gupta, Junitta Guzman, Anu Kumar, Alyssa Manning, Allison Morley, David Roberts and Anisa Tracy for technical assistance.

## MATERIALs and METHODS

### Minimum inhibitory concentration (MIC) and kill kinetics

*M. tuberculosis* was cultured in Middlebrook 7H9 medium supplements with 10% v/v OADC supplement (Becton Dickinson) and 0.05 % w/v Tween 80. Colony forming units (CFUs) were determined on Middlebrook 7H10 agar with 10 % v/v OADC afte 3-4 weeks incubation. Minimum inhibitory concentrations against *M. tuberculosis* were determined as described ^27^. Growth was measured by OD_590_ and RFU after 5 days culture. MIC was defined as the concentration of compound required to inhibit growth of *M. tuberculosis* by 90% and was determined using the Levenberg–Marquardt least-squares plot. Rifampicin controls (maximum inhibition) and a rifampicin dose response was included on every plate. Bacterial viability was measured in log phase cells, or in cells previously starved for >4 days in PBS, 0.05% w/v Tyloxapol ^28^. Compounds were added at multiples of the MIC (at the time the assay was initiated).

### HepG2 cytotoxicity

HepG2 cells were used to measure eukaryotic cytotoxicity. Viability of HepG2 was measured using CellTiter-Glo® reagent (Promega) after 3 days exposure to compounds. IC_50_ was calculated as the compound concentration required to reduce cell viability by 50% and was determined using the Levenberg–Marquardt least-squares plot. Staurosporine controls (maximum inhibition) and a staurosporine dose response was included on every plate.

### Activity against intracellular *M. tuberculosis*

Murine RAW 264.7 macrophages (ATCC TIB-71) were infected with *M. tuberculosis Ds*Red (DREAM8) ^29^ as described at an MOI of 1 for 24 h, washed, harvested and dispensed into 384-well plates ^30^. Infected macrophages were exposed to compounds for 72 hours; cell nuclei were stained with SYBR Green I. Plates were imaged with an ImageXpress Micro High Content Screening System (Molecular Devices) using a 4x objective and FITC and Texas Red channels. MetaXpress was used to determine the integrated intensity of bacteria or macrophages. IC_50_ was calculated as the compound concentration required to reduce bacterial growth by 50%. TC_50_ was calculated as the compound concentration required to reduce macrophage viability by 50%. Isoniazid and staurosporine controls (maximum inhibition) and dose responses were included on every plate.

### ADME

Microsomal stability was measured over 45 min (0, 5, 15, 30, 45 min samples) with 1 µM compound, 0.3 mg/mL human microsomes in 100 mM potassium phosphate buffer pH 7.4, 3 mM MgCl_2_ plus 2 mM NADPH where indicated. Intrinsic clearance was determined as ln(2) /(T_1/2_ [microsomal protein]), where T_1/2_ is the half-life. Permeability was measured across Caco-2 cells cultured in a Millipore 96 well Caco-2 plate. For Apical to Basolateral (A→B) permeability, 10 µM compound was added to the apical (A) side and amount of permeation was determined on the basolateral (B) side; for Basolateral to Apical (B→A) permeability, the compound was added to the B side and the amount of permeation was determine on the A side. Time points were taken at 2 hours.

### General synthetic procedures

Unless otherwise noted, all chemicals used were purchased pure, from commercially available sources such as Sigma Aldrich, VWR, Fisher or other chemical vendors. All reagents and solvents were used as received. ^1^H NMR (300MHz) spectra were recorded on a Bruker Biospin NMR spectrometer. Peak multiplicities are denoted as follows: s = singlet, d = doublet, t = triplet, p = pentet, m = multiplet. Thin layer chromatography was performed using Whatman silica gel 60 Å plates with florescent indicator and visualized using a UV lamp (254 nm) or KMnO_4_ stain. Flash chromatography was performed on Grace with GraceResolv Normal Phase disposable silica columns. High performance liquid chromatography (HPLC) was performed on a Gilson 322 HPLC pump with a Gilson UV/VIS-155 detector and a Phenomenex Gemini C18 column (10 µm, 250 mm x 10 mm). Liquid chromatography electrospray ionization mass spectroscopy (LC-MS/ESI-MS) were acquired on an Agilent LC/MSD-SL with a 1100 HPLC and G1956B mass spectrometer with a Phenomenex Gemini 5 μm C18 110Å 50×3 mm column. High-resolution mass spectra (HRMS-ESI) were acquired by the Mass Spectrometry Laboratory at the University of Michigan on an Agilent Q-TOF HPLC-MS. Purity for molecules was determined by HPLC (as above). In general molecules were >95% pure, except as stated for molecules **17 and 20**. Molecules used in key assays were all >95% pure.

### Reaction scheme for synthesis

General procedure for the synthesis of compounds (excluding **15, 25**, and **27**): Appropriate amine (1.0 eq), alkyl isothiocyanate (1.1 eq - 1.2 eq), triethylamine (2.0 eq), and anhydrous dimethylformamide (DMF) was stirred. The reaction mixture was heated to 90 °C to 100 °C and monitored by TLC and LC-MS. Saturated sodium chloride solution and ethyl acetate were added to the reaction mixture. The aqueous layer was extracted with ethyl acetate. The combined organic layer was dried over anhydrous sodium sulfate and concentrated in vacuo. The crude reaction mixture was purified via column chromatography and recrystallization.

### (1) IDR-0167255/TPN-0002034

#### 1-(benzo[d]thiazol-2-yl)-3-butylthiourea (1)

Triethylamine (0.75 mL, 5.38 mmol) and 2-aminobenzothiazole (0.3998 g, 2.6618 mmol) were added to a stirred solution of butyl isothiocyanate (0.39 mL, 3.23 mmols) and DMF (5.4 mL). The reaction mixture was heated to 90 °C and stirred for 18 h. The reaction mixture was transferred to the separatory funnel with EtOAc and sat. NaCl (aq) solution. The aqueous layer was extracted three times with EtOAc. The combined organic layer was dried over anhydrous Na_2_SO_4_, and the solvent was removed in vacuo. The product was purified by flash column chromatography (SiO_2_; hexanes:EtOAc 95:5 to 0:1) followed by recrystallization in MeOH to yield **1** as colorless crystals (0.0167 g, 2.4%). ^1^H NMR (300 MHz, DMSO-*d*_6_) δ 0.97 (t, *J* = 7.3 Hz, 3H), 1.42 (m, 2H), 1.53 – 1.74 (m, 2H), 3.62 (m, 2H), 7.21 – 7.53 (m, 2H), 7.72 (s, 1H), 7.96 (s, 1H), 10.16 (s, 1H), 11.80 (s, 1H). HRMS-ESI (*m*/*z*): [M+H]^+^ calcd for C_12_H_15_N_3_S_2_, 266.0790, found 266.0784. Purity 99%.

### (11) IDR-0532302/TPN-0002029

#### 1-(benzo[d]imidazol-2-yl)-3-butylthiourea (11)

Triethylamine (0.56 mL, 4 mmol) and butyl isothiocyanate (0.26 mL, 2.2 mmol) were added to a stirred solution of the 2-aminobenzimidazole (0.266 g, 2 mmol) and DMF (10 mL). The reaction mixture was heated to 100 °C and stirred for 15 h. The reaction mixture was transferred to the separatory funnel with EtOAc and sat. NaCl (aq) solution. The aqueous layer was extracted three times with EtOAc. The combined organic layer was dried over anhydrous Na_2_SO_4_, and the solvent was removed in vacuo. The product was purified by recrystallization in ethanol to yield **11** as white crystals (0.042 g, 8.5 %). ^1^H NMR (300 MHz, DMSO-*d*_6_) δ 0.97 (t, *J* = 7.3 Hz, 3H), 1.35 – 1.47 (m, 2H), 1.58 – 1.69 (m, 2H), 3.62 – 3.67 (m, 2H), 7.07 – 7.13 (m, 2H), 7.42 – 7.47 (m, 2H), 10.97 (s, 1H), 11.14 (s, 1H), 11.26 (s, 1H). HRMS-ESI (*m*/*z*): [M+H]^+^ calcd for C_12_H_16_N_4_S, 249.1168, found 249.1170. Purity >95%.

### (12) IDR-0532358/TPN-0002027

#### 1-(benzo[d]oxazol-2-yl)-3-butylthiourea (12)

Triethylamine (0.56 mL, 4 mmol) and butyl isothiocyanate (0.26 mL, 2.2 mmol) were added to a stirred solution of the 2-aminobenzoxazole (0.268 g, 2 mmol) and DMF (10 mL). The reaction mixture was heated to 100 °C and stirred for 15 h. The reaction mixture was transferred to the separatory funnel with EtOAc and sat. NaCl (aq) solution. The aqueous layer was extracted three times with EtOAc. The combined organic layer was dried over anhydrous Na_2_SO_4_, and the solvent was removed in vacuo. The product was purified by recrystallization in ethanol to yield **12** as white crystals (0.044 g, 9 %). ^1^H NMR (300 MHz, DMSO-*d*_6_) δ 0.94 (t, *J* = 7.3 Hz, 3H),1.33 – 1.46 (m, 2H), 1.60 – 1.70 (m, 2H), 3.65 – 3.70 (m, 2H), 7.25 – 7.35 (m, 2H), 7.56 – 7.63 (m, 2H), 10.51 (s, 1H), 12.28 (s, 1H). HRMS-ESI (*m*/*z*): [M+H]^+^ calcd for C_12_H_15_N_3_OS, 250.1020, found 250.1012. Purity 99%.

### (13) IDR-0532303/TPN-0002028

#### 1-butyl-3-(4-chlorobenzo[d]thiazol-2-yl)thiourea (13)

Triethylamine (0.28 mL, 2 mmol) and butyl isothiocyanate (0.13 mL, 1.1 mmol) were added to a stirred solution of the 2-amino-4-chlorobenzthiazole (0.184 g, 1 mmol) and DMF (5 mL). The reaction mixture was heated to 100 °C and stirred for 15 h. The reaction mixture was transferred to the separatory funnel with EtOAc and sat. NaCl (aq) solution. The aqueous layer was extracted three times with EtOAc. The combined organic layer was dried over anhydrous Na_2_SO_4_, and the solvent was removed in vacuo. The product was purified by recrystallization in ethanol to yield **13** as white crystals (0.042 g, 14 %). ^1^H NMR (300 MHz, DMSO-*d*_6_) δ 0.96 (t, *J* = 7.3 Hz, 3H), 1.36 – 1.48 (m, 2H), 1.56 – 1.66 (m, 2H), 3.53 – 3.60 (m, 2H), 7.24 – 7.31 (m, 1H), 7.50 – 7.54 (m, 1H), 7.91 – 7.94 (m, 1H), 9.63 (s, 1H), 12.14 (s, 1H). HRMS-ESI (*m*/*z*): [M+H]^+^ calcd for C_12_H_14_ClN_3_S_2,_ 300.0400, found 300.0393. Purity 99%.

### (14) IDR-0532394/TPN-0002022

#### 1-butyl-3-(thiazol-2-yl)thiourea (14)

Triethylamine (0.56 mL, 4 mmol) and butyl isothiocyanate (0.26 mL, 2.2 mmol) were added to a stirred solution of the 2-aminothiazole (0.200 g, 2 mmol) and DMF (10 mL). The reaction mixture was heated to 100 °C and stirred for 15 h. The reaction mixture was transferred to the separatory funnel with EtOAc and sat. NaCl (aq) solution. The aqueous layer was extracted three times with EtOAc. The combined organic layer was dried over anhydrous Na_2_SO_4_, and the solvent was removed in vacuo. The product was purified by flash column chromatography (SiO_2_; hexanes:EtOAc 1:0 to 0:1) to yield **14** as white powder (0.086 g, 20 %). ^1^H NMR (300 MHz, DMSO-*d*_6_) δ 0.91 (t, *J* = 7.3 Hz, 3H), 1.28 – 1.40 (m, 2H), 1.51 – 1.60 (m, 2H), 3.49 – 3.55 (m, 2H), 7.09 – 7.12 (m, 1H), 7.39 – 7.43 (m, 1H), 9.66 (s, 1H), 11.52 (s, 1H). HRMS-ESI (*m*/*z*): [M+H]^+^ calcd for C_8_H_13_N_3_S_2,_ 216.0630, found 216.0623. Purity 99%.

### (15) IDR-0541252/TPN-0002015

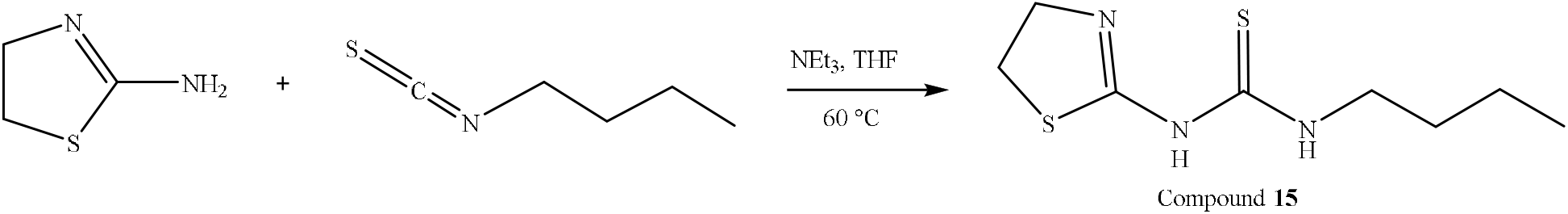

#### 1-butyl-3-(4,5-dihydrothiazol-2-yl)thiourea (15)

Triethylamine (0.20 mL, 1.43 mmol) and 2-amino-2-thiazoline (0.1923 g, 1.387 mmol) were added to a stirred solution of butyl isothiocyanate (0.20 mL, 1.38 mmols) and tetrahydrofuran (2.8 mL). The reaction mixture was heated to 60 °C and stirred overnight. The reaction mixture was extracted three times with CH_2_Cl_2_ and once with saturated NaCl (aq). The organic layers were combined and dried over anhydrous Na_2_SO_4_ and the solvent was removed in vacuo. The product was purified by flash column chromatography (SiO_2_; hexanes:EtOAc 4:1 and CH_2_Cl_2_:MeOH 95:5) to yield **15** as white solids (0.0221 g, 7.4%). 1H NMR (300 MHz, Methylene Chloride-*d*_2_) δ 0.92 (t, *J* = 7.4 Hz, 3H), 1.19 – 1.45 (m, 2H), 1.45 – 1.67 (m, 2H), 3.12 – 3.43 (m, 2H), 3.54 (d, *J* = 6.6 Hz, 2H), 3.84 (overlapping t, 2H), 4.02 (s), 8.20 (s, 1H), 11.02 (s, 1H). HRMS-ESI (m/z): [M+H]+ calcd for C_8_H_15_N_3_S_2_, 218.0790, found 218.0786. Purity 95%.

### (16) IDR-0566795/TPN-0002011

#### 1-butyl-3-(4-methylthiazol-2-yl)thiourea (16)

Triethylamine (0.73 mL, 5.24 mmol) and 2-amino-4-methyl-thiazole (0.3087 g, 2.7039 mmol) were added to a stirred solution of butyl isothiocyanate (0.40 mL, 3.32 mmols) and DMF (5.3 mL). The reaction mixture was heated to 90 °C and stirred for 5 h. The reaction mixture was transferred to the separatory funnel with EtOAc and H_2_O. The aqueous layer was extracted three times with EtOAc. The combined organic layer was extracted once with saturated NaCl (aq), dried over anhydrous Na_2_SO_4_, and the solvent was removed in vacuo. The product was purified by flash column chromatography (SiO_2_; hexanes:EtOAc 9:1 to 0:1) followed by recrystallization in EtOH to yield **16** as colorless crystals (0.0268 g, 4.5%). ^1^H NMR (300 MHz, DMSO-*d*_6_) δ 0.95 (t, *J* = 7.3 Hz, 3H), (h, 7.4 Hz, 2H), 1.59 (p, *J* = 7.1 Hz, 2H), 2.26 (s, 3H), 3.56 (q, *J* = 6.5 Hz, 2H), 6.70 (s, 1H), 9.98 (s, 1H), 11.49 (s, 1H). HRMS-ESI (*m*/*z*): [M+H]^+^ calcd for C_9_H_15_N_3_S_2_, 230.0790, found 230.0784. Purity 99%.

### (17) IDR-0566794/TPN-0002012

#### 1-butyl-3-(4,5-dimethylthiazol-2-yl)thiourea (17)

Triethylamine (0.65 mL, 4.66 mmol) and 4,5-dimethyl-1,3-thiazol-2-amine (0.3060 g, 2.3869 mmol) were added to a stirred solution of butyl isothiocyanate (0.35 mL, 2.90 mmols) and DMF (4.7 mL). The reaction mixture was heated to 90 °C and stirred for 4 h. The reaction mixture was transferred to the separatory funnel with EtOAc and H_2_O. The aqueous layer was extracted three times with EtOAc. The combined organic layer was extracted once with saturated NaCl (aq), dried over anhydrous Na_2_SO_4_, and the solvent was removed in vacuo. The product was purified by flash column chromatography (SiO_2_; hexanes:EtOAc 9:1 to 0:1) followed by recrystallization in EtOH to yield **17** as colorless crystals (0.0255 g, 4.5%). ^1^H NMR (300 MHz, DMSO-*d*_6_) δ 0.94 (t, *J* = 7.3 Hz, 3H), 1.38 (h, *J* = 7.2 Hz, 2H), 1.58 (p, *J* = 7.2 Hz, 2H), 2.15 (s, 3H), 2.22 (s, 3H), 3.54 (m, 2H), 10.00 (s, 1H), 11.35 (s, 1H). HRMS-ESI (*m*/*z*): [M+H]^+^ calcd for C_10_H_17_N_3_S_2_, 244.0940, found 244.0941. Purity 93%.

### (18) IDR-0563572/TPN-0002013

#### 1-butyl-3-(4-methyl-1H-imidazol-2-yl)thiourea (18)

Triethylamine (0.72 mL, 5.2 mmol) and 5-methyl-1H-imidazol-2-ylamine (0.2504 g, 2.578 mmol) were added to a stirred solution of butyl isothiocyanate (0.37 mL, 3.1 mmols) and DMF (5.0 mL). The reaction mixture was heated to 90 °C and stirred for 18 h. The reaction mixture was transferred to the separatory funnel with EtOAc and sat. NaCl (aq) solution. The aqueous layer was extracted three times with EtOAc. The combined organic layer was dried over anhydrous Na_2_SO_4_, and the solvent was removed in vacuo. The product was purified by flash column chromatography (SiO_2_; hexanes:EtOAc 95:5 to 0:1) followed by HPLC (water:MeCN 0 to 100) to yield **18** as a yellow oil (0.0184 g, 3.3%). ^1^H NMR ^1^H NMR (300 MHz, DMSO-*d*_6_) δ 0.94 (t, *J* = 7.3 Hz, 3H), 1.39 (h, 7.1 Hz, 2H), 1.58 (p, *J* = 7.1 Hz, 2H), 2.10 (s, 3H), 3.58 (q, *J* = 6.7 Hz, 2H), 6.49 (s, 1H), 10.05 – 11.20 (m, 3H). LC-MS-ESI (*m*/*z*): [M+H]^+^ calcd for C_12_H_14_N_2_OS_2_, 267.0630, found 267.0627. Purity 99%.

### (19) IDR-0532542/TPN-0002016

#### 1-butyl-3-(1-methyl-1H-pyrazol-5-yl)thiourea (19)

Triethylamine (0.28 mL, 2 mmol) and butyl isothiocyanate (0.13 mL, 1.1 mmol) were added to a stirred solution of the 5-amino-1-methylpyrazole (0.097 g, 1 mmol) and DMF (5 mL). The reaction mixture was heated to 100 °C and stirred for 8 h. The reaction mixture was transferred to the separatory funnel with EtOAc and sat. NaCl (aq) solution. The aqueous layer was extracted three times with EtOAc. The combined organic layer was dried over anhydrous Na_2_SO_4_, and the solvent was removed in vacuo. The product was purified by flash column chromatography (SiO_2_; hexanes:EtOAc 1:0 to 0:1) to yield **19** as white powder (0.046 g, 22 %). ^1^H NMR (300 MHz, Chloroform-*d*) δ 0.93 (t, *J* = 7.3 Hz, 3H), 1.26 – 1.38 (m, 2H), 1.50 – 1.60 (m, 2H), 3.58 – 3.64 (m, 2H), 3.79 (s, 3H), 5.80 – 6.10 (m, 1H), 6.16 – 6.17 (m, 1H), 7.52 (s, 1H), 7.96 (s, 1H). HRMS-ESI (*m*/*z*): [M+H]^+^ calcd for C_9_H_16_N_4_S, 213.1180, found 213.1171. Purity 99%.

### (20) IDR-0571506/TPN-0002004

#### 1-butyl-3-(4-methylpyrimidin-2-yl)thiourea (20)

Triethylamine (1.0 mL, 7.2 mmol) and 2-amino-4-methylpyrimidine (0.4004 g, 3.6690 mmol) were added to a stirred solution of butyl isothiocyanate (0.52 mL, 4.31 mmols) and DMF (7.2 mL). The reaction mixture was heated to 90 °C and stirred for 19 h. The reaction mixture was transferred to the separatory funnel with EtOAc and sat. NaCl (aq) solution. The aqueous layer was extracted three times with EtOAc. The combined organic layer was dried over anhydrous Na2SO4, and the solvent was removed in vacuo. The product was purified by flash column chromatography (SiO2; hexanes:EtOAc 95:5 to 0:1) followed by recrystallization in MeOH to yield **20** as white solids (0.0165 g, 2.0%). ^1^H NMR (300 MHz, DMSO-*d*_6_) δ 0.96 (t, *J* = 7.3 Hz, 3H), 1.42 (h, *J* = 7.4 Hz, 2H), 1.65 (p, *J* = 7.0 Hz, 2H), 3.64 (q, *J* = 6.9, 2H), 7.08 (d, *J* = 5.1 Hz, 1H), 8.53 (d, *J* = 5.1 Hz, 1H), 10.49 (s, 1H), 11.40 (s, 1H). HRMS-ESI (m/z): [M+H]^+^ calcd for C_10_H_16_N_4_S, 225.1180, found 225.1169. Purity 90%.

### (21) IDR-0571504/TPN-0002005

#### 1-butyl-3-(4-methyloxazol-2-yl)thiourea (21)

Triethylamine (1.15 mL, 8.25 mmol) and 2-amino-4-methyloxazole (0.36 mL, 4.29 mmol) were added to a stirred solution of butyl isothiocyanate (0.60 mL, 4.97 mmols) and DMF (8.2 mL). The reaction mixture was heated to 90°C and stirred for 19 h. The reaction mixture was transferred to the separatory funnel with EtOAc and sat. NaCl (aq) solution. The aqueous layer was extracted three times with EtOAc. The combined organic layer was dried over anhydrous Na_2_SO_4_, and the solvent was removed in vacuo. The product was purified by flash column chromatography (SiO_2_; hexanes:EtOAc 95:5 to 0:1) to yield **21** as light yellow solids (0.0465 g, 5.1%). ^1^H NMR (300 MHz, DMSO-*d*_6_) δ 0.95 (t, *J* = 7.3 Hz, 3H), 1.39 (h, *J* = 7.4 Hz, 2H), 1.61 (p, *J* = 7.2 Hz, 2H), 2.08 (s, 3H), 3.62 (q, *J* = 6.6 Hz, 2H), 7.53 (s, 1H), 10.34 (s, *J* = 5.6 Hz, 1H), 11.80 (s, 1H). HRMS-ESI (*m*/*z*): [M+H]^+^ calcd for C_9_H_15_N_3_OS, 214.1020, found 214.1011. Purity 99%.

### (22) IDR-0578471/TPN-0002000

#### 1-butyl-3-(5-methyloxazol-2-yl)thiourea (22)

Triethylamine (1.1 mL, 7.9 mmol) and 5-methyl-1,3-oxazole-2-amine (0.3930 g, 4.006 mmol) were added to a stirred solution of butyl isothiocyanate (0.60 mL, 5.0 mmols) and DMF (8.0 mL). The reaction mixture was heated to 90 °C and stirred for 24 h. The reaction mixture was transferred to the separatory funnel with EtOAc and sat. NaCl (aq) solution. The aqueous layer was extracted three times with EtOAc. The combined organic layer was dried over anhydrous Na_2_SO_4_, and the solvent was removed in vacuo. The product was purified by flash column chromatography (SiO_2_; hexanes:EtOAc 95:5 to 0:1 and 95:5 CH_2_Cl_2_:MeOH) followed by recrystallization in MeOH to yield **22** as beige colored solids (0.0851 g, 9.8%). ^1^H NMR ^1^H NMR (300 MHz, DMSO-*d*_6_) δ 0.94 (t, *J* = 7.3 Hz, 3H), 1.37 (h, *J* = 7.3 Hz, 2H), 1.60 (p, *J* = 7.1 Hz, 2H), 2.26 (s, 3H), 3.61 (q, *J* = 6.6 Hz, 2H), 6.78 (s, 1H), 10.30 (s, 1H), 11.76 (s, 1H). HRMS-ESI (*m*/*z*): [M+H]^+^ calcd for C_9_H_15_N_3_OS, 214.1020, found 214.1011. Purity 99%.

### (23) IDR-0578334/TPN-0002001

#### butyl-3-(4-phenyloxazol-2-yl)thiourea (23)

Triethylamine (0.80 mL, 5.74 mmol) and 4-phenyl-1,3-oxazol-2-amine (0.2475 g, 1.545 mmol) were added to a stirred solution of butyl isothiocyanate (0.42 mL, 3.48 mmols) and DMF (5.6 mL). The reaction mixture was heated to 90 °C and stirred for 28 h. The reaction mixture was transferred to the separatory funnel with EtOAc and sat. NaCl (aq) solution. The aqueous layer was extracted three times with EtOAc. The combined organic layer was dried over anhydrous Na_2_SO_4_, and the solvent was removed in vacuo. The product was purified by flash column chromatography (SiO_2_; hexanes:EtOAc 95:5 to 0:1 and CH_2_Cl_2_:MeOH 9:1 to 0:1) followed by HPLC (water:MeCN 0 to 100) to yield **23** as white solids (0.0042 g, 1%). ^1^H NMR (300 MHz, DMSO-*d*_6_) δ 1.00 (t, *J* = 7.3 Hz, 3H), 1.48 (h, *J* = 7.9 Hz, 2H), 1.69 (p, *J* = 7.2, 6.8 Hz, 2H), 3.68 (q, *J* = 6.4 Hz, 2H), 7.43 (m, 3H), 7.77 (d, *J* = 7.6 Hz, 2H), 8.36 (s, 1H), 10.46 (s, 1H), 11.99 (s, 1H). HRMS-ESI (*m*/*z*): [M+H]^+^ calcd for C_14_H_17_N_3_OS, 276.1170, found 276.1170. Purity 99%

### (24) IDR-0532359/TPN-0002026

#### 1-(benzo[d]thiazol-2-yl)-3-(3-methoxypropyl)thiourea (24)

Triethylamine (0.28 mL, 2 mmol) and 3-methoxypropyl isothiocyanate (0.14 mL, 1.1 mmol) were added to a stirred solution of the 2-aminobenzothiazole (0.150 g, 1 mmol) and DMF (5 mL). The reaction mixture was heated to 100 °C and stirred for 15 h. The reaction mixture was transferred to the separatory funnel with EtOAc and sat. NaCl (aq) solution. The aqueous layer was extracted three times with EtOAc. The combined organic layer was dried over anhydrous Na_2_SO_4_, and the solvent was removed in vacuo. The product was purified by recrystallization in ethanol to yield **24** as white crystals (0.029 g, 10 %). ^1^H NMR (300 MHz, DMSO-*d*_6_) δ 1.82 – 1.90 (m, 2H), 3.27 (s, 3H), 3.41 – 3.46 (m, 2H), 3.61 – 3.70 (m, 2H), 7.26 – 7.93 (m, 4H), 10.16 (s, 1H), 10.81 (s, 1H). HRMS-ESI (*m*/*z*): [M+H]^+^ calcd for C_12_H_15_N_3_OS_2,_ 282.0740, found 282.0734. Purity >95%.

### (25) IDR-0563553/TPN-0002014

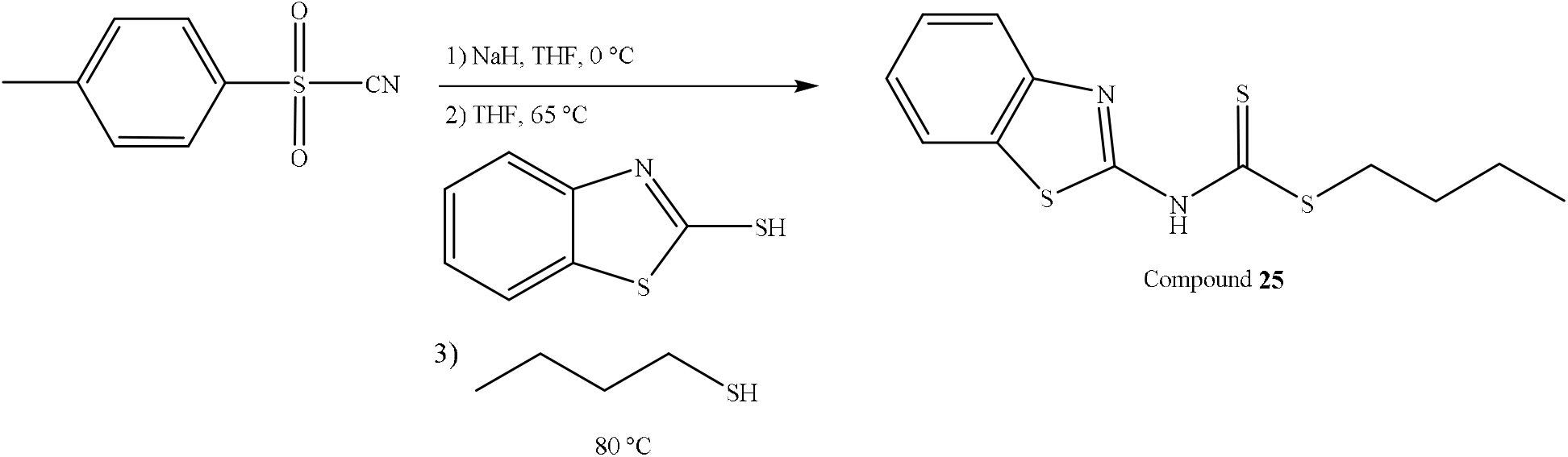

#### 2-thiocyanatobenzo[d]thiazole (25a)

Sodium hydride (0.4482 g, 18.68 mmol) was added to a solution of *p*-toluenesulfonyl cyanide (1.64 g, 9.05 mmol) in THF (6.0 mL) while stirring in an ice-water bath. A solution of 2-mercaptobenzothiazole (1.0112 g, 6.0460 mmol) in THF (6.0 mL) was added to the reaction mixture. The reaction mixture was heated to 65 °C and stirred for 2.5 h. The reaction mixture was cooled in an ice-water bath and quenched with water. The aqueous layer was extracted three times with EtOAc. The combined organic layers were dried over anhydrous Na_2_SO_4_ and the solvent was removed in vacuo to yield 1.65 g of solid.

### butyl benzo[d]thiazol-2-ylcarbamodithioate (25)

A solution of **25a** (0.4013 g, 2.0873 mmol) and 1-butanethiol (3.0 mL, 28 mmol) was heated to 80 °C while stirring. The reaction mixture was stirred for 3 h. The reaction mixture was extracted three times with CH_2_Cl_2_ and once with saturated NaCl (aq). The organic layers were combined and dried over anhydrous Na_2_SO_4_ and the solvent was removed in vacuo. The product was purified by flash column chromatography (SiO_2_; hexanes:EtOAc 4:1) to yield **25** as yellow solids (0.0269 g, 4.6%). ^1^H NMR (300 MHz, Chloroform-*d*) δ 0.97 (t, *J* = 7.3 Hz, 3H), 1.47 (h, *J* = 14.5, 7.4 Hz, 2H), 1.74 (p, *J* = 15.2, 7.0 Hz, 2H), 3.33 (t, *J* = 7.5 Hz, 2H), 7.40 (t, *J* = 7.9 Hz, 1H), 7.51 (t, *J* = 7.7 Hz, 1H), 7.77 (dd, *J* = 21.2, 8.0 Hz, 2H). HRMS-ESI (*m*/*z*): [M+H]^+^ calcd for C_12_H_14_N_2_S_3_, 283.0400, found 283.0397. Purity 95%.

### (26) IDR-0532360/TPN-0002025

#### 1-(benzo[d]thiazol-2-yl)-3-phenethylthiourea (26)

Triethylamine (0.28 mL, 2 mmol) and 2-phenylethyl isothiocyanate (0.16 mL, 1.1 mmol) were added to a stirred solution of the 2-aminobenzothiazole (0.150 g, 1 mmol) and DMF (5 mL). The reaction mixture was heated to 100 °C and stirred for 15 h. The reaction mixture was transferred to the separatory funnel with EtOAc and sat. NaCl (aq) solution. The aqueous layer was extracted three times with EtOAc. The combined organic layer was dried over anhydrous Na_2_SO_4_, and the solvent was removed in vacuo. The product was purified by recrystallization in ethanol to yield **29** as white crystals (0.035 g, 11 %). ^1^H NMR (300 MHz, DMSO-*d*_6_) δ 2.94 – 2.99 (m, 2H), 3.84 – 3.86 (m, 2H), 7.24 – 7.44 (m, 7H), 7.57 – 7.60 (m, 1H), 7.89 – 7.92 (m, 1H), 10.27 (s, 1H), 11.86 (s, 1H). HRMS-ESI (*m*/*z*): [M+H]^+^ calcd for C_16_H_15_N_3_S_2,_ 314.0790, found 314.0783. Purity 99%.

### (27) IDR-0579434/TPN-0001998

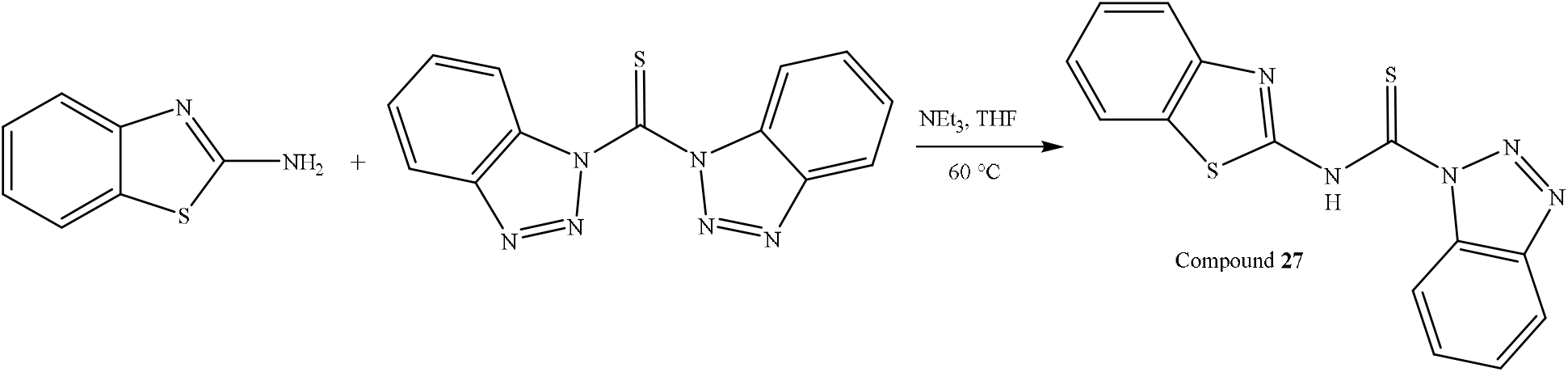

#### 3-(benzo[d]thiazol-2-yl)-1-butyl-1-methylthiourea (27)

Bis(1-benzotriazolyl)methanethione (0.5051 g, 1.802 mmol) was added to a stirred solution of 2-aminobenzothiazole (0.2199 g, 1.464 mmols) and THF (3.0 mL). The reaction mixture was stirred at ambient temperature for 21 h. Triethylamine (0.41 mL, 2.9 mmol) and N-methylbutylamine (0.21 mL, 1.8 mmol) were added to the reaction mixture and the reaction was left to stir for another 32 hours at ambient temperature, and at 50 °C for 42 hours. The reaction was monitored by LC-MS and TLC. The reaction mixture was transferred to the separatory funnel with EtOAc and water. The aqueous layer was extracted three times with EtOAc. The combined organic layer was dried over anhydrous Na_2_SO_4_, and the solvent was removed in vacuo. The product was purified by flash column chromatography (SiO_2_; hexanes:EtOAc 95:5 to 0:1) followed by HPLC (water:MeCN 0 to 100) to yield **27** as a white solid (0.1149 g, 28.1%). ^1^H NMR ^1^H NMR (300 MHz, DMSO-*d*_6_) δ 0.96 (t, *J* = 7.4 Hz, 3H), 1.34 (m, 2H), 1.61 (p, *J* = 9.0 Hz, 2H), 3.33 (s, 3H), 3.89 (t, *J* = 7.5 Hz, 2H), 7.13 – 7.48 (m, 3H), 7.76 (d, *J* = 6.3 Hz, 1H), 12.71 (s, 1H). LC-MS-ESI (*m*/*z*): [M+H]^+^ calcd for C_13_H_18_N_3_S_2_, 280.1, found 280.0. Purity 99%.

### (28) IDR-0532392/TPN-0002024

#### 1-(benzo[d]thiazol-2-yl)-3-isobutylthiourea (28)

Triethylamine (0.56 mL, 4 mmol) and isobutyl isothiocyanate (0.28 mL, 2.2 mmol) were added to a stirred solution of the 2-aminobenzothiazole (0.3 g, 2 mmol) and DMF (10 mL). The reaction mixture was heated to 100 °C and stirred for 15 h. The reaction mixture was transferred to the separatory funnel with EtOAc and sat. NaCl (aq) solution. The aqueous layer was extracted three times with EtOAc. The combined organic layer was dried over anhydrous Na_2_SO_4_, and the solvent was removed in vacuo. The product was purified by recrystallization in ethanol to yield **28** as white crystals (0.052 g, 10 %). ^1^H NMR (300 MHz, DMSO-*d*_6_) δ 0.96 – 0.98 (d, *J* = 6.6 Hz, 6H), 1.94 – 2.03 (m, 1H), 3.20 – 3.44 (m, 2H), 7.26 – 7.41 (m, 2H), 7.65 – 7.91 (m, 2H), 10.25 (s, 1H), 11.81 (s, 1H). HRMS-ESI (*m*/*z*): [M+H]^+^ calcd for C_12_H_15_N_3_S_2,_ 266.0790, found 266.0796. Purity >95%.

### (29) IDR-0578333/TPN-0002002

#### 1-isobutyl-3-(4-methyloxazol-2-yl)thiourea (29)

Triethylamine (1.4 mL, 10 mmol) and 2-amino-4-methyloxazole (0.43 mL, 5.1 mmol) were added to a stirred solution of isobutyl isothiocyanate (0.75 mL, 6.1 mmols) and DMF (10.0 mL). The reaction mixture was heated to 90 °C and stirred for 23 h. The reaction mixture was transferred to the separatory funnel with EtOAc and sat. NaCl (aq) solution. The aqueous layer was extracted three times with EtOAc. The combined organic layer was dried over anhydrous Na_2_SO_4_, and the solvent was removed in vacuo. The product was purified by flash column chromatography (SiO_2_; hexanes:EtOAc 95:5 to 0:1 and CH_2_Cl_2_:MeOH 95:5 to 0:1) followed by HPLC (water:MeCN 0 to 100) to yield **29** as white solids (0.0132 g, 1.2%). ^1^H NMR (300 MHz, DMSO-*d*_6_) δ 0.98 (d, *J* = 6.7 Hz, 6H), 1.87-2.15 (m, 4H), 3.48 (t, *J* = 6.0 Hz, 2H), 7.54 (s, 1H), 10.46 (s, 1H), 11.83 (s,

1H). HRMS-ESI (*m*/*z*): [M+H]^+^ calcd for C_9_H_15_N_3_OS, 214.1020, found 214.1007. Purity 99%.

### (30) IDR-0571381/TPN-0002010

#### 1-(benzo[d]thiazol-5-yl)-3-butylthiourea (30)

Triethylamine (0.65 mL, 4.66 mmol) and 5-aminobenzothiazole (0.3494 g, 2.3262 mmol) were added to a stirred solution of butyl isothiocyanate (0.34 mL, 2.82 mmols) and DMF (4.5 mL). The reaction mixture was heated to 90 °C and stirred for 6.5 h. The reaction mixture was transferred to the separatory funnel with EtOAc and H_2_O. The aqueous layer was extracted three times with EtOAc. The combined organic layer was extracted once with saturated NaCl (aq), dried over anhydrous Na_2_SO_4_, and the solvent was removed in vacuo. The product was purified by flash column chromatography (SiO_2_; CH_2_Cl_2_:MeOH 1:0 to 9:1) followed by recrystallization in acetone to yield **30** as white solids (0.0479 g, 7.7%). ^1^H NMR (300 MHz, DMSO-*d*_6_) δ 0.95 (t, *J* = 7.3 Hz, 3H), 1.37 (h, *J* = 7.6 Hz, 2H), 1.57 (p, *J* = 7.4 Hz, 2H), 3.51 (q, *J* = 6.8 Hz, 2H), 7.48 (d, *J* = 8.9 Hz, 1H), 7.90 (s, 1H), 8.11 (d, *J* = 8.6 Hz, 1H), 8.23 (s, 1H), 9.42 (s, 1H), 9.67 (s, 1H). HRMS-ESI (*m*/*z*): [M+H]^+^ calcd for C_12_H_15_N_3_S_2_, 266.0790, found 266.0786. Purity 99%.

### (31) IDR-0532393/TPN-0002023

#### 1-(4-chlorobenzo[d]thiazol-2-yl)-3-propyl thiourea (31)

Triethylamine (0.28 mL, 2 mmol) and propyl isothiocyanate (0.11 mL, 1.1 mmols) were added to a stirred solution of the 2-amino-4-chlorobenzthiazole (0.184 g, 1 mmol) and DMF (5 mL). The reaction mixture was heated to 100 °C and stirred for 15 h. The reaction mixture was transferred to the separatory funnel with EtOAc and sat. NaCl (aq) solution. The aqueous layer was extracted three times with EtOAc. The combined organic layer was dried over anhydrous Na_2_SO_4_, and the solvent was removed in vacuo. The product was purified by recrystallization in ethanol to yield **31** as white crystals (0.030 g, 10.5 %). ^1^H NMR (300 MHz, DMSO-*d*_6_) δ 0.97 (t, *J* = 7.3 Hz, 3H), 1.57 – 1.69 (m, 2H), 3.50 – 3.56 (m, 2H), 7.25 – 7.30 (m, 1H), 7.49 – 7.53 (m, 1H), 7.91 – 7.94 (m, 1H), 9.56 (s, 1H), 12.13 (s, 1H). HRMS-ESI (*m*/*z*): [M+H]^+^ calcd for C_11_H_12_ClN_3_S_2,_ 286.0240, found 286.0241. Purity 98%.

### (32) IDR-0571501/TPN-0002008

#### 1-butyl-3-(5-chlorobenzo[d]oxazol-2-yl) thiourea (32)

Triethylamine (0.33 mL, 2.37 mmol) and 2-amino-5-chlorobenzoxazole (0.3998 g, 2.3716 mmol) were added to a stirred solution of butyl isothiocyanate (0.30 mL, 2.49 mmols) and DMF (4.6 mL). The reaction mixture was heated to 90 °C and stirred for 23 h. The reaction mixture was transferred to the separatory funnel with EtOAc and sat. NaCl (aq) solution. The aqueous layer was extracted three times with EtOAc. The combined organic layer was dried over anhydrous Na_2_SO_4_, and the solvent was removed in vacuo. The product was purified by flash column chromatography (SiO_2_; hexanes:EtOAc 95:5 to 0:1) followed by recrystallization in CHCl_3_ to yield **32** as white solids (0.0382 g, 5.7%). ^1^H NMR (300 MHz, DMSO-*d*_6_) δ 0.97 (t, *J* = 7.3 Hz, 3H), 1.43 (h, *J* = 7.9 Hz, 2H), 1.67 (p, *J* = 7.3 Hz, 2H), 3.69 (q, *J* = 6.7 Hz, 2H), 7.35 (d, *J* = 6.2 Hz, 1H), 7.69 (d, *J* = 8.3 Hz, 2H), 10.42 (s, 1H), 12.45 (s, 1H). HRMS-ESI (*m*/*z*): [M+H]^+^ calcd for C_12_H_14_ClN_3_OS, 284.0630, found 284.0622. Purity 98%.

### (33) IDR-0571502/TPN-0002007

#### 1-butyl-3-(6-chlorobenzo[d]oxazol-2-yl)thiourea (33)

Triethylamine (0.33 mL, 2.37 mmol) and 6-chloro-1,3-benzoxazole-2-amine (0.4005 g, 2.3757 mmol) were added to a stirred solution of butyl isothiocyanate (0.30 mL, 2.49 mmols) and DMF (4.6 mL). The reaction mixture was heated to 90 °C and stirred for 26 h. The reaction mixture was transferred to the separatory funnel with EtOAc and sat. NaCl (aq) solution. The aqueous layer was extracted three times with EtOAc. The combined organic layer was dried over anhydrous Na_2_SO_4_, and the solvent was removed in vacuo. The product was purified by flash column chromatography (SiO_2_; hexanes:EtOAc 95:5 to 0:1) to yield **33** as white solids (0.0439 g, 6.5%). ^1^H NMR (300 MHz, DMSO-*d*_6_) δ 0.97 (t, *J* = 7.3 Hz, 3H), 1.42 (h, *J* = 7.4 Hz, 2H), 1.67 (p, *J* = 7.3 Hz, 2H), 3.69 (q, *J* = 6.6 Hz, 2H), 7.41 (d, *J* = 8.5, 1H), 7.61 (d, *J* = 8.4 Hz, 1H), 7.88 (s, 1H), 10.43 (s, 1H), 12.42 (s, 1H). HRMS-ESI (*m*/*z*): [M+H]^+^ calcd for C_12_H_14_ClN_3_OS, 284.0630, found 284.0621. Purity 99%.

### (34) IDR-0571503/TPN-0002006

#### 1-butyl-3-(7-chlorobenzo[d]oxazol-2-yl)thiourea (34)

Triethylamine (0.33 mL, 2.37 mmol) and 7-chloro-1,3-benzoxazole-2-amine (0.4003 g, 2.3745 mmol) were added to a stirred solution of butyl isothiocyanate (0.30 mL, 2.49 mmols) and DMF (4.6 mL). The reaction mixture was heated to 90 °C and stirred for 24 h. The reaction mixture was transferred to the separatory funnel with EtOAc and sat. NaCl (aq) solution. The aqueous layer was extracted three times with EtOAc. The combined organic layer was dried over anhydrous Na_2_SO_4_, and the solvent was removed in vacuo. The product was purified by flash column chromatography (SiO_2_; hexanes:EtOAc 95:5 to 0:1) to yield **34** as white solids (0.0529 g, 7.9%). ^1^H NMR (300 MHz, DMSO-*d*_6_) δ 0.97 (t, *J* = 7.3 Hz, 3H), 1.43 (h, *J* = 7.5 Hz, 2H), 1.68 (p, *J* = 7.3 Hz, 2H), 3.69 (q, *J* = 6.6 Hz, 2H), 7.31 – 7.46 (m, 2H), 7.58 (d, *J* = 7.2 Hz, 1H), 10.40 (s, 1H), 12.45 (s, 1H). HRMS-ESI (*m*/*z*): [M+H]^+^ calcd for C_12_H_14_ClN_3_OS, 284.0630, found 284.0621. Purity 98%.

### (35) IDR-0576402/TPN-0002003

#### 1-butyl-3-(5-methoxybenzo[d]oxazol-2-yl)thiourea (35)

Triethylamine (0.70 mL, 5.0 mmol) and 5-methoxybenzo[d]oxazol-2-amine (0.3995 g, 2.4336 mmol) were added to a stirred solution of butyl isothiocyanate (0.35 mL, 2.90 mmols) and DMF (5.0 mL). The reaction mixture was heated to 90 °C and stirred for 39 h. The reaction mixture was transferred to the separatory funnel with EtOAc and sat. NaCl (aq) solution. The aqueous layer was extracted three times with EtOAc. The combined organic layer was dried over anhydrous Na_2_SO_4_, and the solvent was removed in vacuo. The product was purified by flash column chromatography (SiO_2_; hexanes:EtOAc 95:5 to 0:1) to yield **35** as white solids (0.0810 g, 11.9%). ^1^H NMR (300 MHz, DMSO-*d*_6_) δ 0.97 (t, *J* = 7.3 Hz, 3H), 1.42 (h, *J* = 6.7 Hz, 2H), 1.67 (p, 6.9 Hz, 2H), 3.58 – 3.73 (q, *J* = 6.7 Hz, 2H), 3.83 (s, 3H), 6.95 (d, *J* = 6.9 Hz, 1H), 7.33 (s, 1H), 7.49 (d, *J* = 8.7 Hz, 1H), 10.46 (s, 1H), 12.23 (s, 1H). HRMS-ESI (*m*/*z*): [M+H]^+^ calcd for C_13_H_17_N_3_O_2_S, 280.1120, found 280.1118. Purity 99%.

## REFERENCES

1 World Health Organization, Global tuberculosis report 2021.

2 J. A. Maddry, S. Ananthan, R. C. Goldman, J. V. Hobrath, C. D. Kwong, C. Maddox, L. Rasmussen, R. C. Reynolds, J. A. Secrist, M. I. Sosa, E. L. White and W. Zhang, Tuberculosis (Edinb), 2009, 89, 354–363.

3 S. Ananthan, E. R. Faaleolea, R. C. Goldman, J. V. Hobrath, C. D. Kwong, B. E. Laughon, J. A. Maddry, A. Mehta, L. Rasmussen, R. C. Reynolds, J. A. Secrist, N. Shindo, D. N. Showe, M. I. Sosa, W. J. Suling and E. L. White, Tuberculosis (Edinb), 2009, 89, 334–353.

4 R. C. Reynolds, S. Ananthan, E. Faaleolea, J. V. Hobrath, C. D. Kwong, C. Maddox, L. Rasmussen, M. I. Sosa, E. Thammasuvimol, E. L. White, W. Zhang and J. A. Secrist, Tuberculosis (Edinb), 2012, 92, 72–83.

5 M. Silva-Miranda, E. Ekaza, A. Breiman, K. Asehnoune, D. Barros-Aguirre, K. Pethe, F. Ewann, P. Brodin, L. Ballell-Pages and F. Altare, Antimicrob Agents Chemother, 2015, 59, 693–697.

6 L. Ballell, R. H. Bates, R. J. Young, D. Alvarez-Gomez, E. Alvarez-Ruiz, V. Barroso, D. Blanco, B. Crespo, J. Escribano, R. González, S. Lozano, S. Huss, A. Santos-Villarejo, J. J. Martín-Plaza, A. Mendoza, M. J. Rebollo-Lopez, M. Remuiñan-Blanco, J. L. Lavandera, E. Pérez-Herran, F. J. Gamo-Benito, J. F. García-Bustos, D. Barros, J. P. Castro and N. Cammack, ChemMedChem, 2013, 8, 313–321.

7 T. Parish, Expert Opin Drug Discov, 2020, 15, 349–358.

8 J. V. Early, A. Casey, M. A. Martinez-Grau, I. C. Gonzalez Valcarcel, M. Vieth, J. Ollinger, M. A. Bailey, T. Alling, M. Files, Y. Ovechkina and T. Parish, Antimicrob Agents Chemother, 2016, 60, 3608–3616.

9 J. Early, J. Ollinger, C. Darby, T. Alling, S. Mullen, A. Casey, B. Gold, J. Ochoada, T. Wiernicki, T. Masquelin, C. Nathan, P. A. Hipskind and T. Parish, ACS Infect Dis, 2019, 5, 272–280.

10 B. Gold, T. Warrier and C. Nathan, Methods Mol Biol, 2015, 1285, 293–315.

11 C. Deb, C.-M. Lee, V. S. Dubey, J. Daniel, B. Abomoelak, S. Pawar, L. Rogers and P. E. Kolattukudy, PLoS ONE, 2009, 4, 15.

12 S. S. Grant, T. Kawate, P. P. Nag, M. R. Silvis, K. Gordon, S. A. Stanley, E. Kazyanskaya, R. Nietupski, A. Golas, M. Fitzgerald, S. Cho, S. G. Franzblau and D. T. Hung, ACS Chem Biol, 2013, 8, 2224–2234.

13 T. Warrier, M. Martinez-Hoyos, M. Marin-Amieva, G. Colmenarejo, E. Porras-De Francisco, A. I. Alvarez-Pedraglio, M. T. Fraile-Gabaldon, P. A. Torres-Gomez, L. Lopez-Quezada, B. Gold, J. Roberts, Y. Ling, S. Somersan-Karakaya, D. Little, N. Cammack, C. Nathan and A. Mendoza-Losana, ACS Infect Dis, 2015, 1, 580–585.

14 C. M. Darby, H. I. Ingólfsson, X. Jiang, C. Shen, M. Sun, N. Zhao, K. Burns, G. Liu, S. Ehrt, J. D. Warren, O. S. Andersen, O. S. Anderson, S. J. Brickner and C. Nathan, PLoS One, 2013, 8, e68942.

15 B. Gold and C. Nathan, Microbiol Spectr,, DOI:10.1128/microbiolspec.TBTB2-0031-2016.

16 D. J. Payne, M. N. Gwynn, D. J. Holmes and D. L. Pompliano, Nat Rev Drug Discov, 2007, 6, 29–40.

17 S. A. Bonnett, J. Ollinger, S. Chandrasekera, S. Florio, T. O’Malley, M. Files, J.-A. Jee, J. Ahn, A. Casey, Y. Ovechkina, D. Roberts, A. Korkegian and T. Parish, ACS Infect Dis, 2016, 2, 893–902.

18 G. L. Abrahams, A. Kumar, S. Savvi, A. W. Hung, S. Wen, C. Abell, C. E. Barry, D. R. Sherman, H. I.M. Boshoff and V. Mizrahi, Chem Biol, 2012, 19, 844–854.

19 S. A. Bonnett, D. Dennison, M. Files, A. Bajpai and T. Parish, PLoS One, 2018, 13, e0198059.

20 E. S. Chung, W. C. Johnson and B. B. Aldridge, Nat Rev Microbiol,, DOI:10.1038/s41579-022-00721-0.

21 S. Vijay, D. N. Vinh, H. T. Hai, V. T. N. Ha, V. T. M. Dung, T. D. Dinh, H. N. Nhung, T. T. B. Tram, B. B. Aldridge, N. T. Hanh, D. D. A. Thu, N. H. Phu, G. E. Thwaites and N. T. T. Thuong, Front Microbiol, 2017, 8, 2296.

22 J. Graham, C. E. Wong, J. Day, E. McFaddin, U. Ochsner, T. Hoang, C. L. Young, W. Ribble, M. A. DeGroote, T. Jarvis and X. Sun, Bioorg Med Chem Lett, 2018, 28, 3177–3181.

23 M. A. De Groote, T. C. Jarvis, C. Wong, J. Graham, T. Hoang, C. L. Young, W. Ribble, J. Day, W. Li, M. Jackson, M. Gonzalez-Juarrero, X. Sun and U. A. Ochsner, Front Microbiol, 2018, 9, 2231.

24 H. M. Abdel-Rahman and M. A. Morsy, J Enz Inhib Med Chem, 2007, 22, 57–64.

25 A. L. Moure, G. Narula, F. Sorrentino, A. Bojang, C. K. M. Tsui, C. Sao Emani, E. Porras-De Francisco, B. Díaz, M. J. Rebollo-López, P. A. Torres-Gómez, E. M. López-Román, I. Camino, P. Casado Castro, L. Guijarro López, F. Ortega, L. Ballell, D. Barros-Aguirre, M. Remuiñán Blanco and Y. Av-Gay, J Med Chem, 2020, 63, 4732–4748.

26 S. S. Grant, S. Wellington, T. Kawate, C. A. Desjardins, M. R. Silvis, C. Wivagg, M. Thompson, K. Gordon, E. Kazyanskaya, R. Nietupski, N. Haseley, N. Iwase, A. M. Earl, M. Fitzgerald and D. T. Hung, Cell Chem Biol, 2016, 23, 666–677.

27 J. Ollinger, M. A. Bailey, G. C. Moraski, A. Casey, S. Florio, T. Alling, M. J. Miller and T. Parish, PLoS One, 2013, 8, e60531.

28 J. Early and T. Alling, Methods Mol Biol, 2015, 1285, 269–279.

29 P. Carroll, J. Muwanguzi-Karugaba and T. Parish, BMC Res Notes, 2018, 11, 685.

30 A. J. Manning, Y. Ovechkina, A. McGillivray, L. Flint, D. M. Roberts and T. Parish, Methods, 2017, 127, 3–11.

