## Supplemental data for "Identification of 2-amino benzothiazoles with bactericidal activity against *Mycobacterium tuberculosis*"

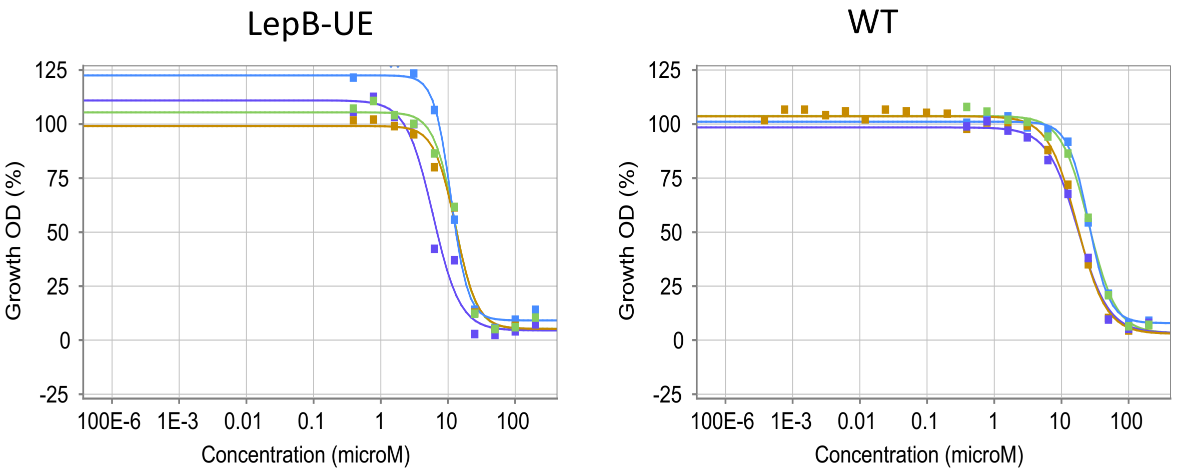


**Figure S1.** Compound response curves for a single batch of the seed molecule against the LepB under-expressing strain (LepB-UE) and the wild-type strain (WT). Data are from four independent experiments with the same batch of compound.

| Molecule # | Secretion $IC_{50}$ (µM) | Molecule # | Secretion $IC_{50}$ (µM) |
|---|---|---|---|
| **1** | >200 | **19** | >200 |
| **8** | >200 | **23** | >200 |
| **11** | >200 | **26** | >200 |
| **12** | >200 | **27** | >200 |
| **13** | >200 | **33** | >200 |
| **14** | >200 | **37** | >200 |
| **16** | >200 | **37** | >200 |

**Table S1. Molecules were tested for the ability to inhibit secretion using membrane fractions.**

Cell membrane fractions were isolated as described (Bonnett SA, Ollinger J, Chandrasekera S, Florio S, O'Malley T, Files M, Jee JA, Ahn J, Casey A, Ovechkina Y, Roberts D, Korkegian A, Parish T. 2016. A target-based whole cell screen approach to identify potential inhibitors of Mycobacterium tuberculosis signal peptidase. ACS Infectious Diseases 2:893–902).

**Method**

Cell membrane fractions were isolated from γ-irradiated whole *M. tuberculosis* H37Rv (NR-14819; BEI resources). Cells (10 g) were washed once in 100 mL of 20 mM Tris (pH 8.0), 250 mM NaCl, and 0.05% w/v Tween 80 buffer and suspended in 80 mL of 20 mM Tris, pH 8, and 1% Triton X-114. The suspension was incubated for 1 h with rotation, and the pellet was recovered by centrifugation. One hundred milligrams of Triton X-114 membrane fraction pellet was solubilized in 1 mL of XTractor buffer (Clontech) and transferred to a 2 mL screw-cap tube containing 0.1 mm silica. The tubes were agitated in a BeadBeater (BioSpec) for five 30 s pulses. The resulting cell membrane fraction was diluted 1:5 with 50 mM Tris-HCl, pH 8.0, and assayed for SPase activity using the fluorogenic peptide substrate 3-nitrotyrosine (NO)-YFSASALA~KI-aminobenzoic acid-OH (ABZ) (California Peptide). Assays were performed in 96-well black plates with 2 µM substrate, 200 µM test compound, and 3% DMSO in 50 mM Tris-HCl, pH 8.0 (total volume = 100 µL). Reactions were initiated by the addition of 10 µL of cell membrane fraction and incubated at 37 °C for 4 h. Activity was analyzed by fluorescence (Ex 315 nm/Em 410 nm). Each assay plate contained controls with no compound or no cell membrane fraction.

**Table S2. Metabolic stability**. Metabolic stability was measured using human liver microsomes. Compound **1**

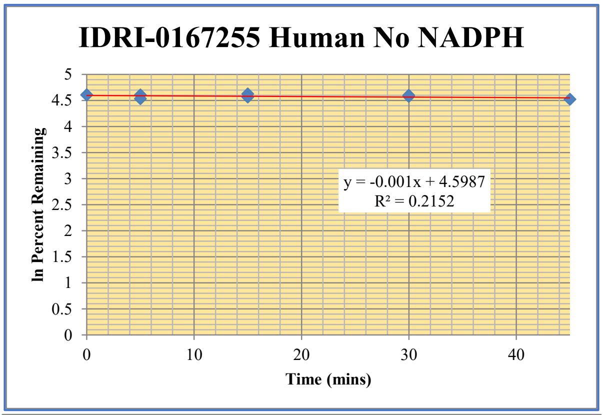


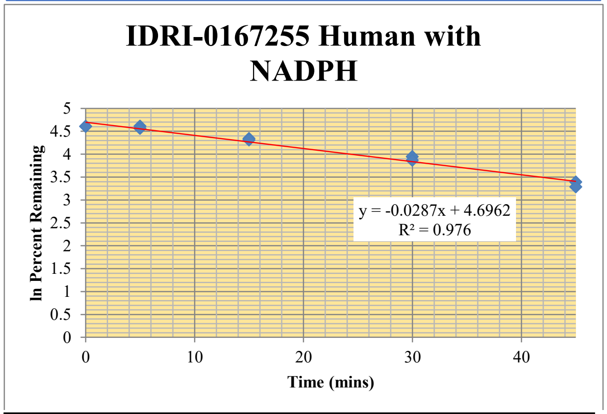


| Time (min) | $\%RE_1$ | $\%RE_2$ | $\%RE_{avg}$ |
|---|---|---|---|
| 0 | 100% | 100% | 100% |
| 5 | 100% | 93.1% | 96.6% |
| 15 | 102% | 95.9% | 99.0% |
| 30 | 97.2% | 99.2% | 98.2% |
| 45 | | 92.1% | 92.1% |

| Time (min) | $\%RE_1$ | $\%RE_2$ | $\%RE_{avg}$ |
|---|---|---|---|
| 0 | 100% | 100% | 100% |
| 5 | 100% | 95.8% | 98.1% |
| 15 | 76.8% | 74.4% | 75.6% |
| 30 | 51.7% | 47.8% | 49.7% |
| 45 | 29.7% | 26.7% | 28.2% |

Compound **12**

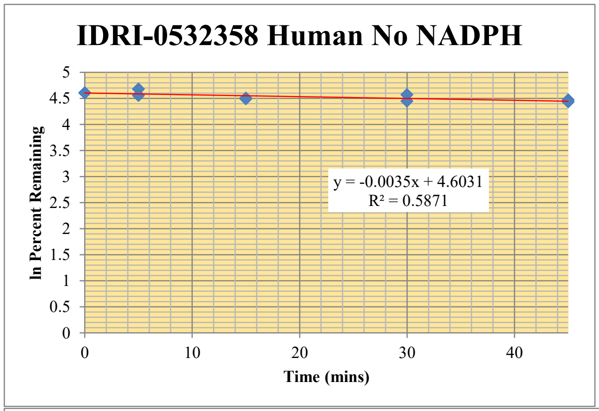


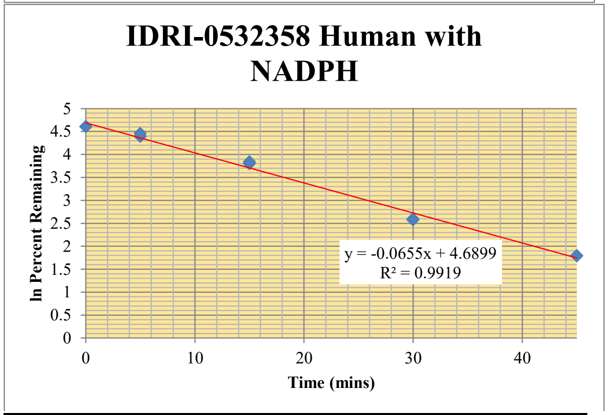


| Time (min) | %$RE_1$ | %$RE_2$ | %$RE_{avg}$ |
|---|---|---|---|
| 0 | 100% | 100% | 100% |
| 5 | 108% | 95.4% | 102% |
| 15 | 91% | 89.3% | 89.9% |
| 30 | 96.6% | 85.7% | 91.1% |
| 45 | 87.5% | 84.2% | 85.9% |

| Time (min) | %$RE_1$ | %$RE_2$ | %$RE_{avg}$ |
|---|---|---|---|
| 0 | 100% | 100% | 100% |
| 5 | 81% | 86.2% | 83.7% |
| 15 | 44.7% | 46.6% | 45.6% |
| 30 | 13.4% | 13.1% | 13.3% |
| 45 | 6.0% | 6.0% | 6.0% |

Compound **24**

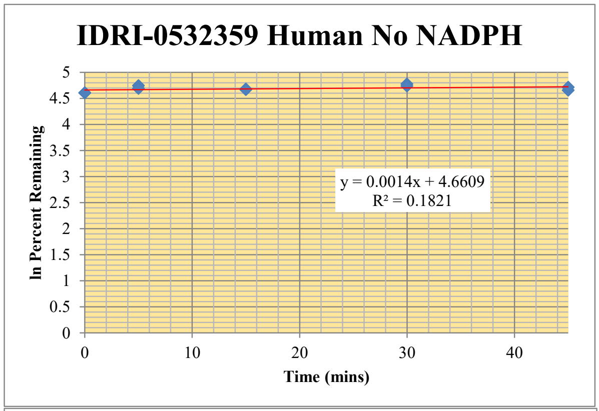


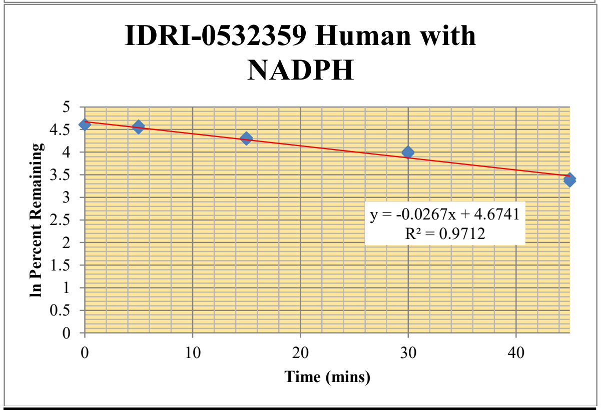


| Time (min) | $\%RE_1$ | $\%RE_2$ | $\%RE_{avg}$ |
|---|---|---|---|
| 0 | 100% | 100% | 100% |
| 5 | 115% | 109% | 112% |
| 15 | 108% | 106.4% | 107.0% |
| 30 | 119% | 114% | 116% |
| 45 | 105% | 111% | 108% |

| Time (min) | $\%RE_1$ | $\%RE_2$ | $\%RE_{avg}$ |
|---|---|---|---|
| 0 | 100% | 100% | 100% |
| 5 | 92.7% | 97.5% | 95.1% |
| 15 | 72.6% | 74.9% | 73.8% |
| 30 | 53.0% | 55.0% | 54.0% |
| 45 | 28.7% | 30.4% | 29.5% |

Compound 1**4**

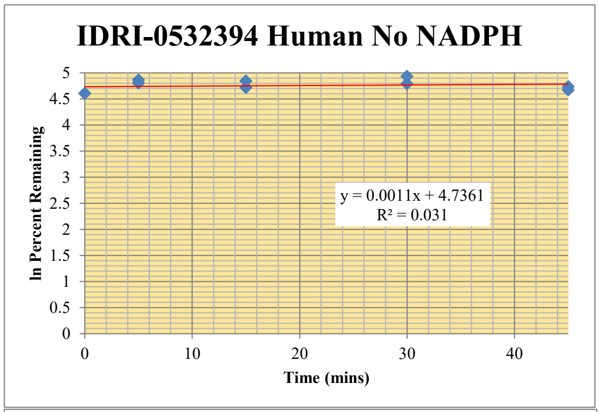


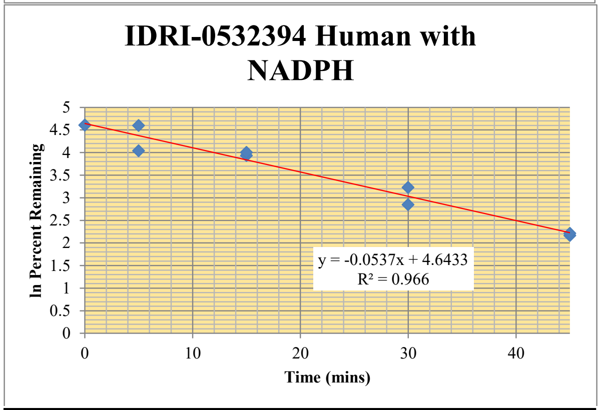


| Time (min) | %$RE_1$ | %$RE_2$ | %$RE_{avg}$ |
|---|---|---|---|
| 0 | 100% | 100% | 100% |
| 5 | 130% | 122% | 126% |
| 15 | 127% | 112% | 119% |
| 30 | 139% | 121% | 130% |
| 45 | 107% | 114% | 111% |

| Time (min) | %$RE_1$ | %$RE_2$ | %$RE_{avg}$ |
|---|---|---|---|
| 0 | 100% | 100% | 100% |
| 5 | 56.8% | 99.1% | 78.0% |
| 15 | 51.0% | 54.9% | 53.0% |
| 30 | 17.3% | 25.2% | 21.2% |
| 45 | 8.7% | 9.1% | 8.9% |

Compound **21**

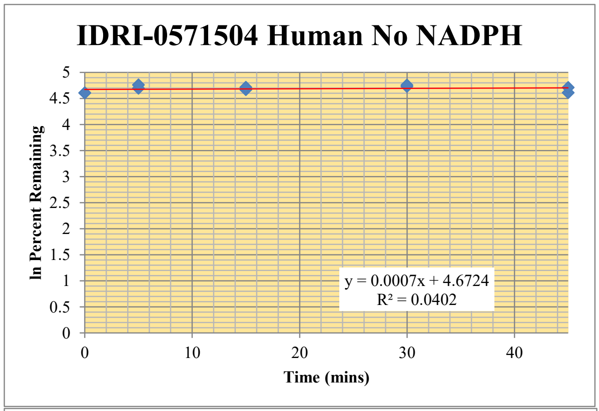


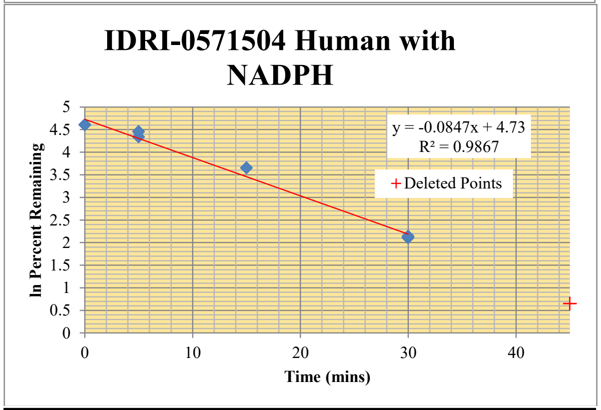


| Time (min) | %$RE_1$ | %$RE_2$ | %$RE_{avg}$ |
|---|---|---|---|
| 0 | 100% | 100% | 100% |
| 5 | 109% | 117% | 113% |
| 15 | 106% | 111% | 109% |
| 30 | 113% | 116% | 115% |
| 45 | 101% | 111% | 106% |

| Time (min) | %$RE_1$ | %$RE_2$ | %$RE_{avg}$ |
|---|---|---|---|
| 0 | 100% | 100% | 100% |
| 5 | 76.8% | 86.0% | 81.4% |
| 15 | 38.7% | | 38.7% |
| 30 | 8.2% | 8.5% | 8.4% |
| 45 | 1.9% | 1.9% | 1.9% |

**Table S3. Permeability.** Permeability was measured using Caco-2 cells after 1 and 2 hours incubation with compound. (A→B) is Apical to Basolateral; (B→A) is Basolateral to Apical. Data is expressed as permeability ($10^{-6}$ cm $s^{-1}$). dQ/dt is rate of permeation, C0 is initial concentration of test agent.

| *Test Article* | *Assay Duration* | *direction* | *value* | *1st replicate* | *2nd replicate* | *mean* | *recovery* |
|---|---|---|---|---|---|---|---|
| **IDRI-0571504 (Cpd 21)** | **1 hr** | A→B | dQ/dt[a] | 3.7E-04 | 4.6E-04 | 4.2E-04 | 59% |
| | | A→B | **$C_0$[b]** | 125 | 134 | 129 | |
| | | B→A | dQ/dt[a] | 3.2E-04 | 4.5E-04 | 3.8E-04 | 77% |
| | | B→A | **$C_0$[b]** | 138 | 144 | 141 | |
| **IDRI-0167255 (Cpd 1)** | **1 hr** | A→B | dQ/dt[a] | 9.1E-06 | 1.0E-05 | 9.7E-06 | 40% |
| | | A→B | **$C_0$[b]** | 168 | 177 | 172 | |
| | | B→A | dQ/dt[a] | 1.8E-05 | 3.0E-05 | 2.4E-05 | 72% |
| | | B→A | **$C_0$[b]** | 154 | 145 | 150 | |
| **IDRI-0532358 (Cpd 12)** | **1 hr** | A→B | dQ/dt[a] | 2.5E-05 | 4.2E-05 | 3.4E-05 | 48% |
| | | A→B | **$C_0$[b]** | 80.6 | 88.2 | 84.4 | |
| | | B→A | dQ/dt[a] | 2.6E-05 | 2.9E-05 | 2.7E-05 | 86% |
| | | B→A | **$C_0$[b]** | 74.3 | 60.8 | 67.5 | |
| **IDRI-0532359 Cpd (24)** | **1 hr** | A→B | dQ/dt[a] | 1.1E-04 | 1.1E-04 | 1.1E-04 | 45% |
| | | A→B | **$C_0$[b]** | 53.4 | 56.5 | 55.0 | |
| | | B→A | dQ/dt[a] | 7.2E-05 | 8.8E-05 | 8.0E-05 | 69% |
| | | B→A | **$C_0$[b]** | 50.2 | 58.6 | 54.4 | |
| **IDRI-0532394 (Cpd 14)** | **1 hr** | A→B | dQ/dt[a] | 2.4E-04 | 2.8E-04 | 2.6E-04 | 57% |
| | | A→B | **$C_0$[b]** | 75.0 | 80.4 | 77.7 | |
| | | B→A | dQ/dt[a] | 2.1E-04 | 2.4E-04 | 2.3E-04 | 72% |
| | | B→A | **$C_0$[b]** | 77.0 | 86.7 | 81.9 | |

| *Test Article* | *Assay Duration* | *direction* | *value* | *1st replicate* | *2nd replicate* | *mean* | *recovery* |
|---|---|---|---|---|---|---|---|
| **IDRI-0571504 (Cpd 21)** | **2 hr** | A→B | dQ/dt[a] | 3.4E-04 | 2.6E-04 | 3.0E-04 | 56% |
| | | A→B | $\mathbf{C_0}$[b] | 129 | 128 | 129 | |
| | | B→A | dQ/dt[a] | 2.3E-04 | 3.2E-04 | 2.8E-04 | 67% |
| | | B→A | $\mathbf{C_0}$[b] | 136.4 | 127.5 | 131.9 | |
| **IDRI-0167255 (Cpd 1)** | **2 hr** | A→B | dQ/dt[a] | 1.9E-05 | 2.2E-05 | 2.1E-05 | **22%** |
| | | A→B | $\mathbf{C_0}$[b] | 137 | 128 | 132 | |
| | | B→A | dQ/dt[a] | 3.8E-05 | 3.8E-05 | 3.8E-05 | 55% |
| | | B→A | $\mathbf{C_0}$[b] | 146.5 | 134.3 | 140.4 | |
| **IDRI-0532358 (Cpd 12)** | **2 hr** | A→B | dQ/dt[a] | 3.4E-05 | 3.1E-05 | 3.2E-05 | **33%** |
| | | A→B | $\mathbf{C_0}$[b] | 62.8 | 56.6 | 59.7 | |
| | | B→A | dQ/dt[a] | 2.7E-05 | 3.3E-05 | 3.0E-05 | 61% |
| | | B→A | $\mathbf{C_0}$[b] | 48.4 | 46.1 | 47.3 | |
| **IDRI-0532359 Cpd (24)** | **2 hr** | A→B | dQ/dt[a] | 9.1E-05 | 8.4E-05 | 8.8E-05 | 54% |
| | | A→B | $\mathbf{C_0}$[b] | 46.3 | 46.1 | 46.2 | |
| | | B→A | dQ/dt[a] | 9.6E-05 | 1.3E-04 | 1.1E-04 | 69% |
| | | B→A | $\mathbf{C_0}$[b] | 66.3 | 67.9 | 67.1 | |
| **IDRI-0532394 (Cpd 14)** | **2 hr** | A→B | dQ/dt[a] | 1.6E-04 | 1.4E-04 | 1.5E-04 | 59% |
| | | A→B | $\mathbf{C_0}$[b] | 63.2 | 62.2 | 62.7 | |
| | | B→A | dQ/dt[a] | 1.4E-04 | 1.8E-04 | 1.6E-04 | 65% |
| | | B→A | $\mathbf{C_0}$[b] | 83.5 | 83.3 | 83.4 | |

| CYP Isoform | Substrate | Substrate Concentration | HLM Concentration | Incubation Time | Positive Control |
|---|---|---|---|---|---|
| CYP2B6 | Bupropion | 100 µM | 0.25 mg/ml | 10 min | Ticlopidine |
| CYP2C8 | Amodiaquine | 5 µM | 0.25 mg/ml | 10 min | Montelukast |
| CYP2C9 | Tolbutamide | 100 µM | 0.5 mg/ml | 15 min | Sulfaphenazole |
| CYP2C19 | Mephenytoin | 100 µM | 0.25 mg/ml | 60 min | Tranylcypromine |
| CYP2D6 | Dextromethorphan | 5 µM | 0.5 mg/ml | 10 min | Quinidine |
| CYP3A4 | Midazolam | 2.5 µM | 0.25 mg/ml | 10 min | Ketoconazole |
| CYP3A4 | Testosterone | 50 µM | 0.25 mg/ml | 10 min | Ketoconazole |

| Test Article/ | | | $IC_{50}$ (µM) | | | | | | |
|---|---|---|---|---|---|---|---|---|---|
| Control Inhibitor | TPN | Molecule | CYP2B6 | CYP2C8 | CYP2C9 | CYP2C19 | CYP2D6 | CYP3A4 - Testosterone | CYP3A4 - Midazolam |
| IDRI-0167255 | TPN-0002034 | **1** | > 20 | > 20 | > 20 | 4.6 | > 20 | > 20 | > 20 |
| IDRI-0532358 | TPN-0002027 | **12** | > 20 | > 20 | > 20 | 12.7 | > 20 | > 20 | > 20 |
| IDRI-0532359 | TPN-0002022 | **14** | > 20 | > 20 | 13.2 | 13.5 | > 20 | > 20 | > 20 |
| IDRI-0532394 | TPN-0002005 | **21** | > 20 | > 20 | > 20 | > 20 | > 20 | 11.9 | > 20 |
| IDRI-0571504 | TPN-0002026 | **24** | > 20 | > 20 | > 20 | > 20 | > 20 | > 20 | > 20 |
| | | Ticlopidine | 0.61 | | | | | | |
| | | Montelukast | | 0.16 | | | | | |
| | | Sulfaphenazole | | | 0.11 | | | | |
| | | Tranylcypromine | | | | 7.6 | | | |
| | | Quinidine | | | | | 0.028 | | |
| | | Ketoconazole | | | | | | 0.011 | 0.024 |

**Table S3. Inhibition of cytochrome P450s.**
Compounds were tested for inhibition of P450s using the enzyme substrate pairs and conditions noted. $IC_{50}$s ($\mu$M) were calculated. Control P450 inhibitors were included for each isoform.

#### Method

For each compound, the analytical method was developed as follows. Analyte signal was optimized for each compound by ESI positive or negative ionization mode. An MS2 scan or an SIM scan was used to optimize fragmenter voltage and a product ion analysis was used to identify the best fragment for analysis. The collision energy was optimized using a product ion or MRM scan. An ionization ranking was assigned indicating the compound's ease of ionization. Samples were analyzed by LC/MS/MS using an Agilent 6410 mass spectrometer coupled with an Agilent 1200 HPLC and a CTC PAL chilled autosampler, all controlled by MassHunter software (Agilent). After separation on a C18 reverse phase HPLC column (Agilent Zorbax StableBond 3.5$\mu$m, 2.1 x 30 mm) using an acetonitrile-water gradient system, peaks were analyzed by mass spectrometry (MS) using ESI ionization in MRM mode.

Serial dilutions of a 10 mM stock solution were prepared in acetonitrile:DMSO (9:1). The final DMSO concentration was < 0.2%. Compounds were incubated with human liver microsomes in the presence of 2 mM NADPH in 100 mM potassium phosphate (pH 7.4) containing 5 mM magnesium chloride and a probe substrate, in a 200 $\mu$l assay final volume in duplicate. The probe substrate concentrations including bupropion (CYP2B6), amodiaquine (CYP2C8), tolbutamide (CYP2C9), mephenytoin (CYP2C19), dextromethorphan (CYP2D6) and midazolam/testosterone (CYP3A4/5) were used as summarized in the Table. The selective CYP inhibitors were screened alongside the test agents as a positive control including ticlopidine (CYP2B6), montelukast (CYP2C8), sulfaphenazole (CYP2C9), tranylcypromine (CYP2C19), quinidine (CYP2D6) and ketoconazole (CYP3A4/5). After incubation at 37oC, reactions were terminated by addition of methanol containing internal standard (propranolol) for analytical quantification. The quenched samples were kept on ice for 10 min and centrifuged at 4 °C for 10 min. The supernatant was removed and the probe substrate metabolite was analyzed by LC-MS/MS (Agilent Technologies Triple Quad LC/MS). A decrease in the formation of the metabolite compared to vehicle control was used to calculate an IC50 value (the test concentration which produces 50% inhibition).

| Molecule | Mean plasma fraction Unbound Fuplasma (%) | Mean plasma fraction Bound (%) | Post-Assay Recovery |
|---|---|---|---|
| Propranolol | 28.00% | 72.% | 89.4% |
| Warfarin | 0.67% | 99.3% | 88.6% |
| **1** | 0.10% | 99.9% | 84.3% |
| **12** | 0.20% | 99.8% | 80.1% |
| **14** | 0.98% | 99.0% | 83.8% |
| **21** | 3.7% | 96.3% | 83.9% |
| **24** | 5.5% | 94.5% | 76.5% |

**Table S4. Plasma protein binding**
Protein binding was measured in human plasma using Rapid Equilibrium Dialysis (RED) devices (Pierce) according to the manufacturers' instructions; 5 µM compounds in plasma were dialysed against PBS for 4 h at 37C. Compounds were detected by LC-MS/MS in plasma and PBS

### High Resolution Mass Spectrometry (HRMS) Spectra

#### (1) IDR-0167255/TPN-0002034

[M+H]+ calcd for $C_{12}H_{15}N_3S_2$, 266.0790, found 266.0784.

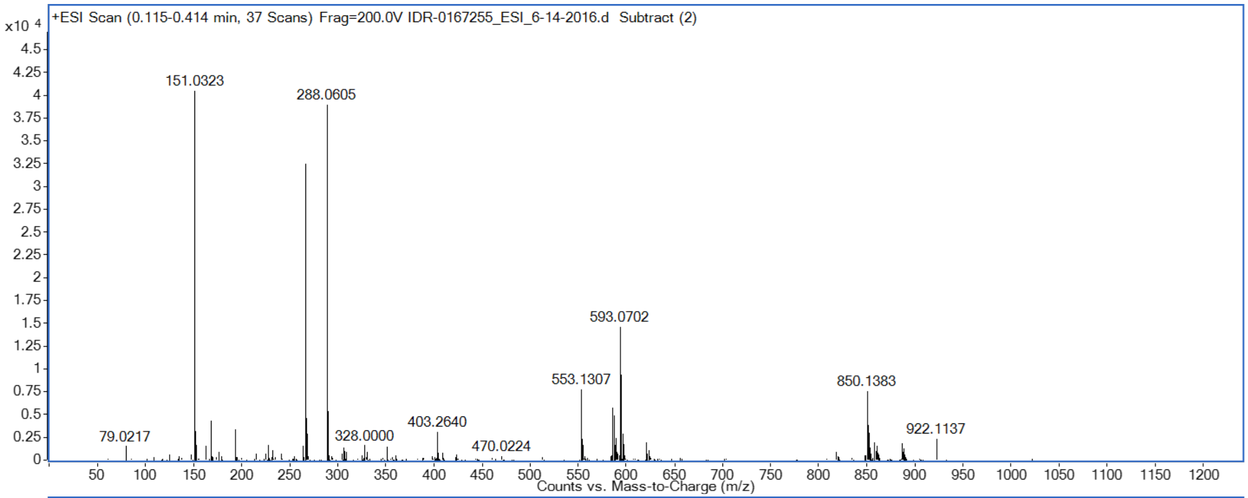


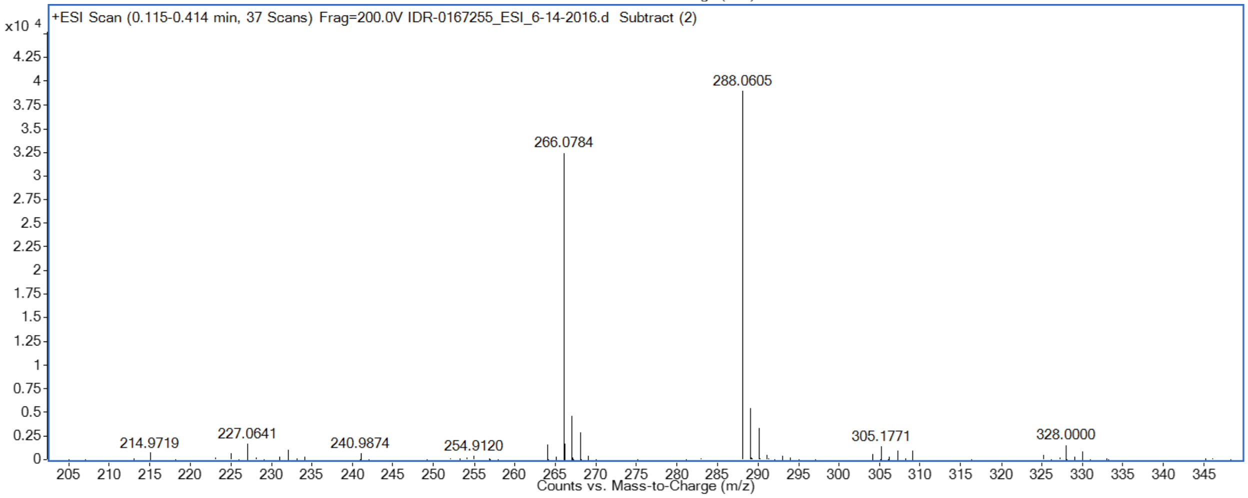


#### (11) IDR-0532302/TPN-0002029

[M+H]+ calcd for $C_{12}H_{16}N_4S$: 249.1168

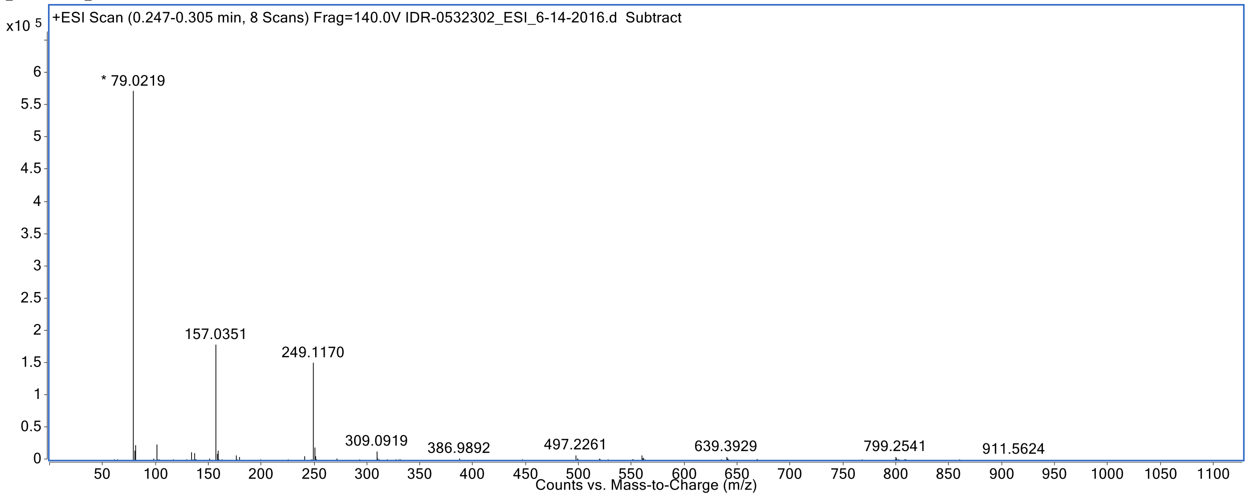

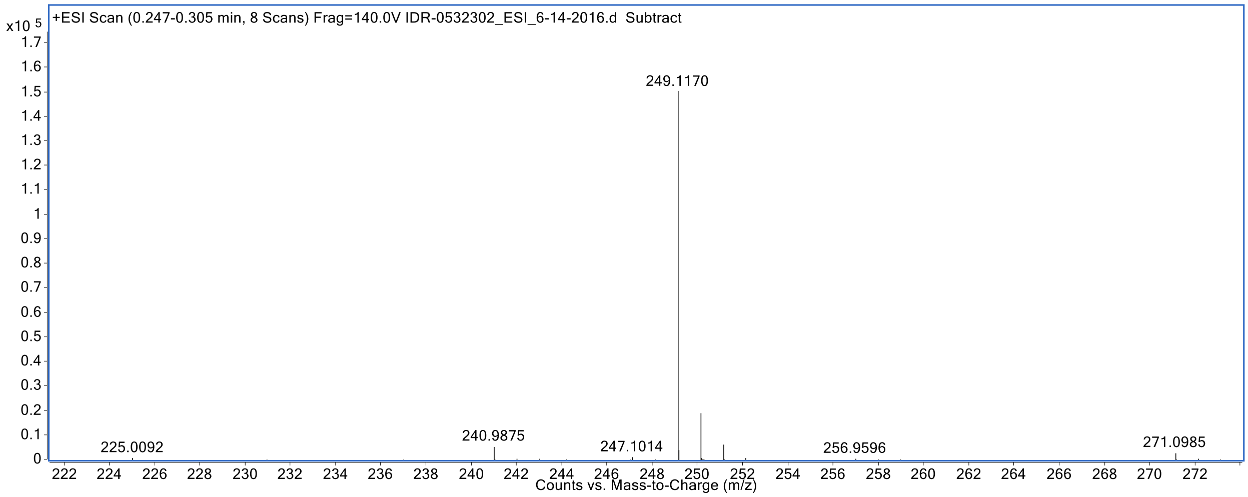


**(12) IDR-0532358/TPN-0002027**

$[M+H]^+$ calcd for $C_{12}H_{15}N_3OS$: 250.1009

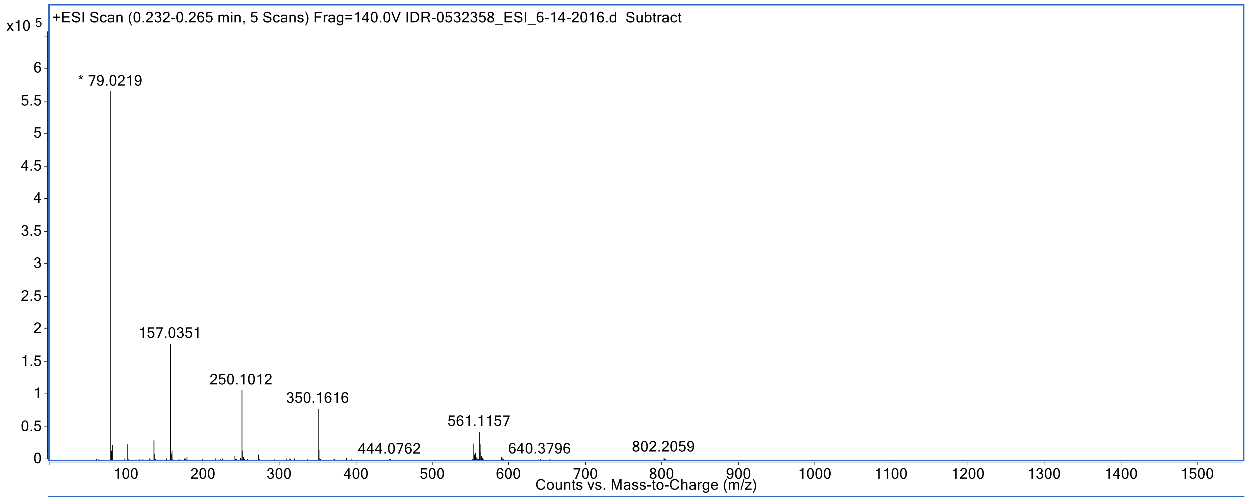


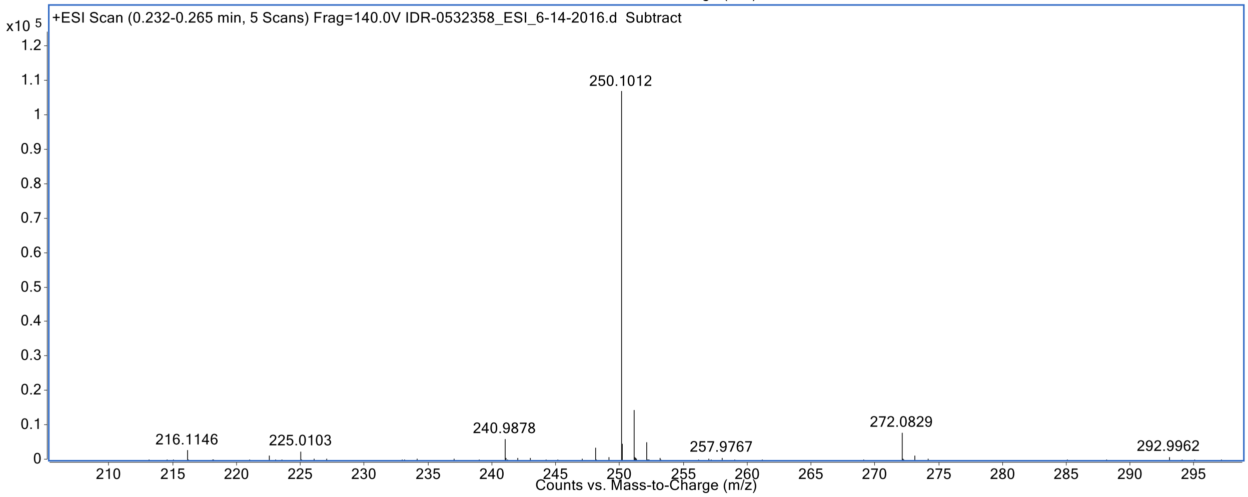

#### (13) IDR-0532303/TPN-0002028

$[M+H]^+$ calcd for $C_{12}H_{14}ClN_3S_2$: 300.0390

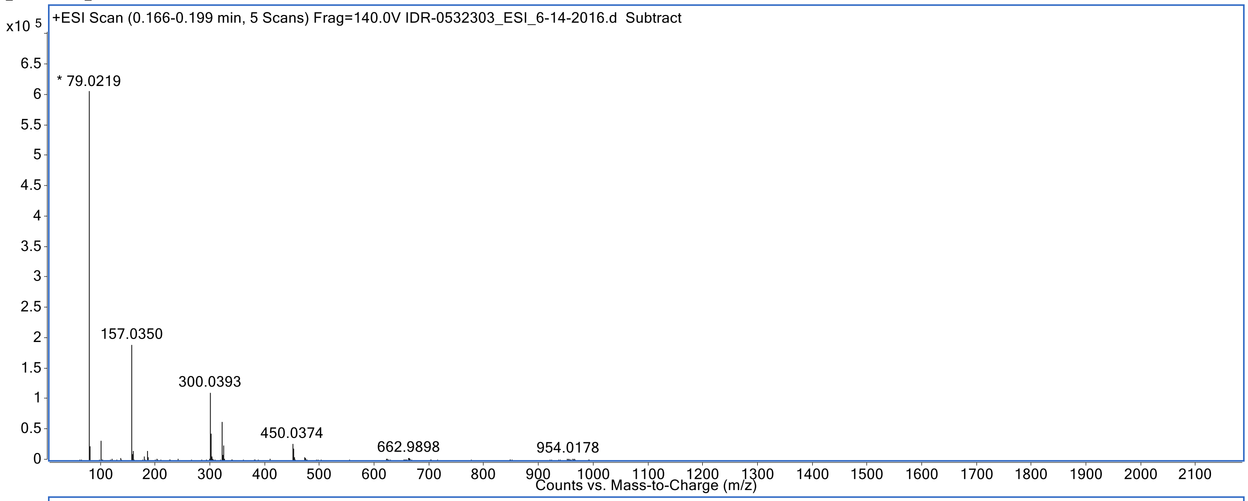


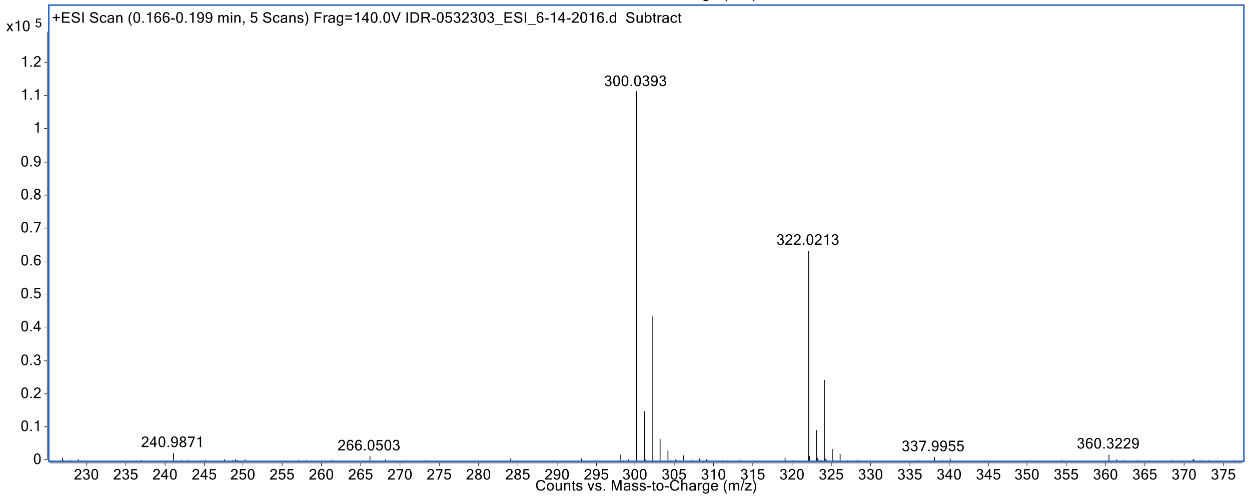


#### (14) IDR-0532394/TPN-0002022

$[M+H]^+$ calcd for $C_8H_{13}N_3S_2$: 216.0624

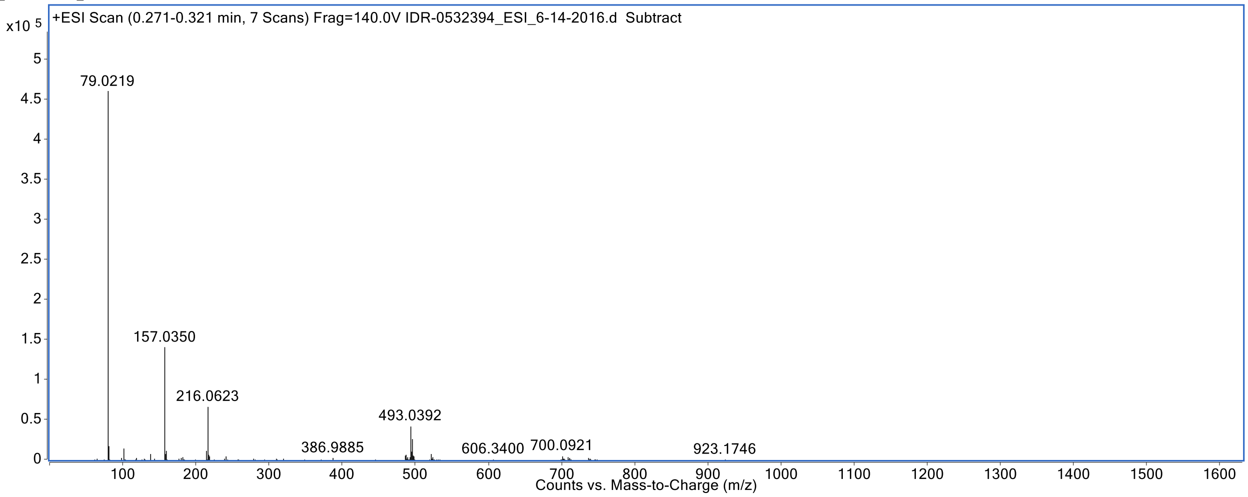

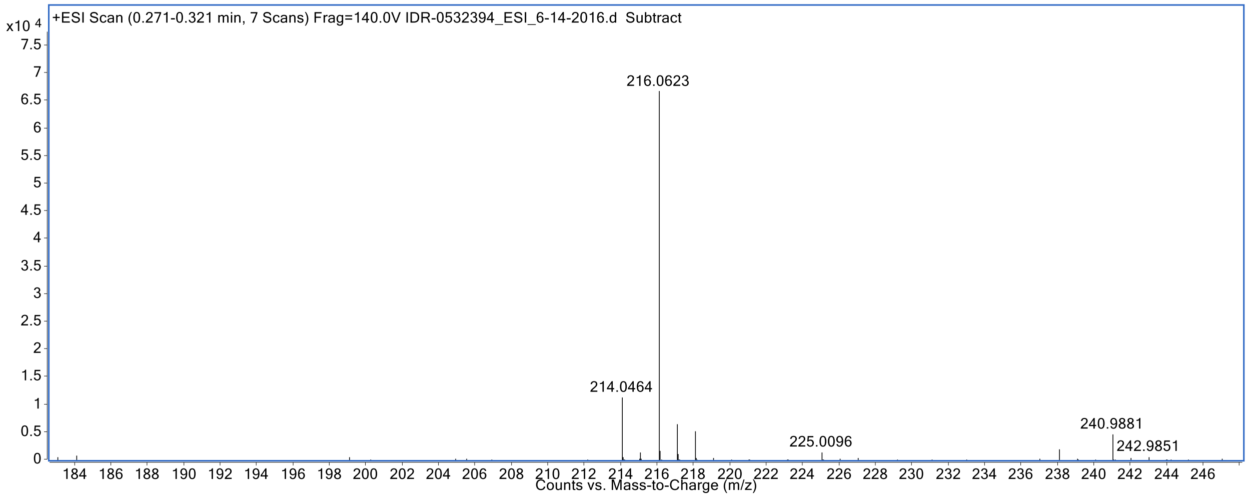


**(15) IDR-0541252/TPN-0002015**

[M+H]+ calcd for C8H15N3S2, 218.0790, found 218.0786.

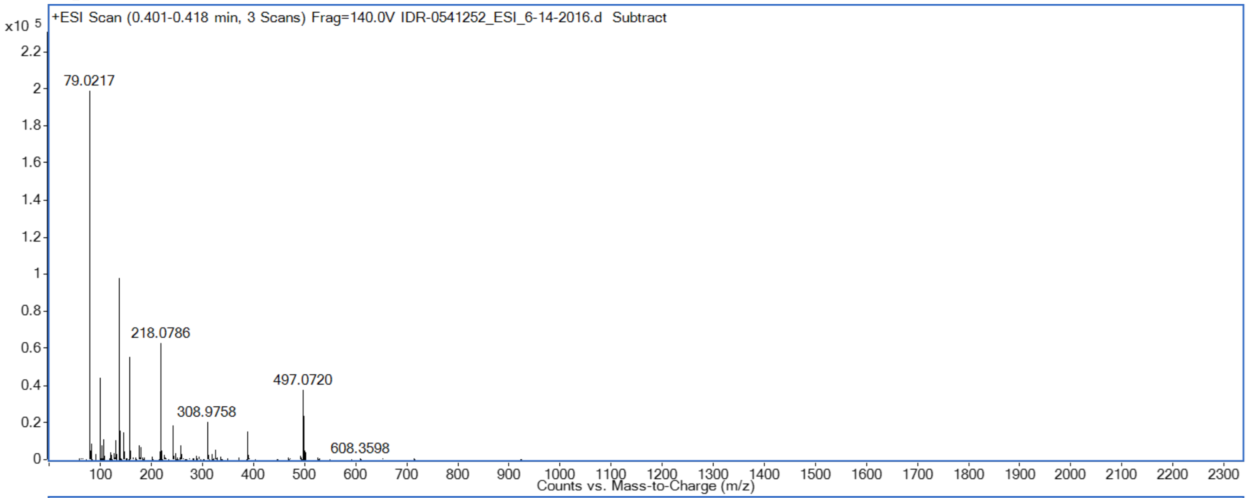


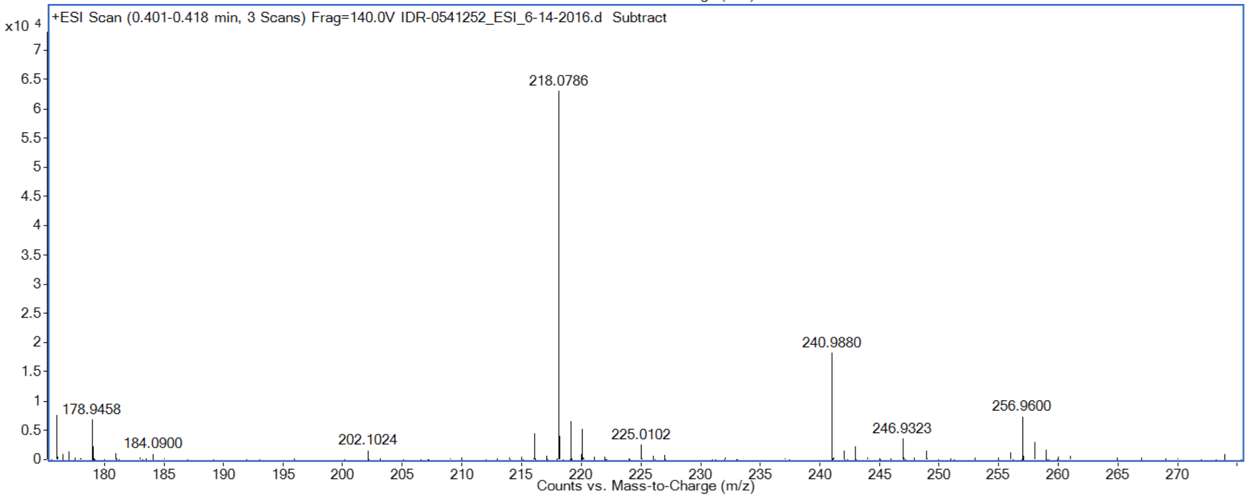


**(16) IDR-0566795/TPN-0002011**

$[M+H]^+$ calcd for $C_9H_{15}N_3S_2$, 230.0790, found 230.0784

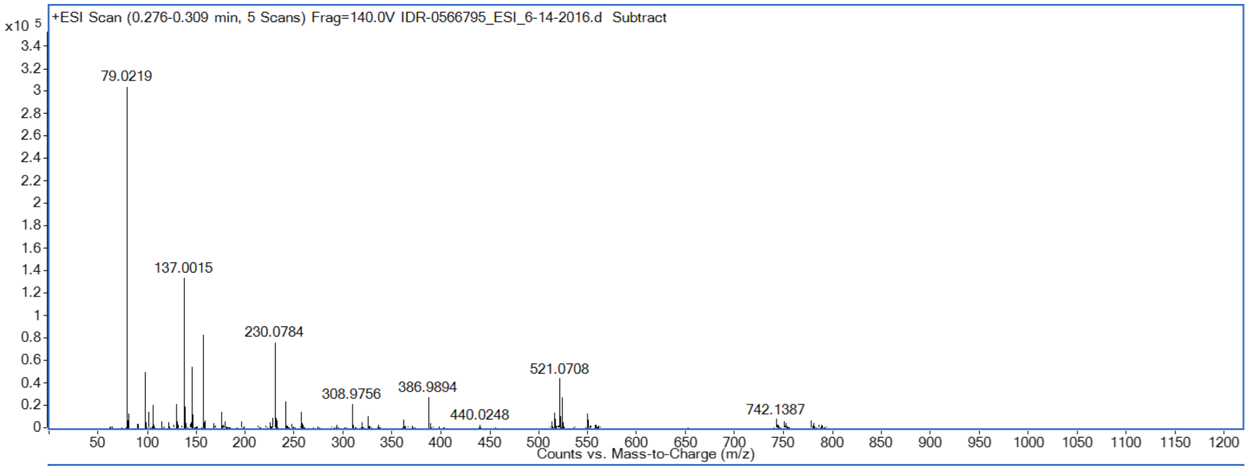


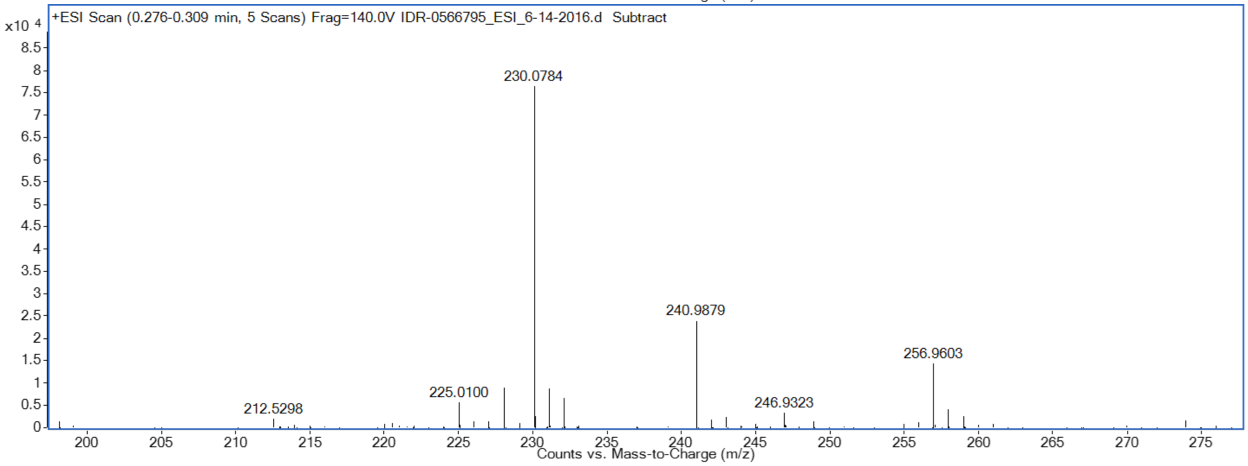


#### (17) IDR-0566794/TPN-0002012

$[M+H]^+$ calcd for $C_{10}H_{17}N_3S_2$, 244.0940, found 244.0941

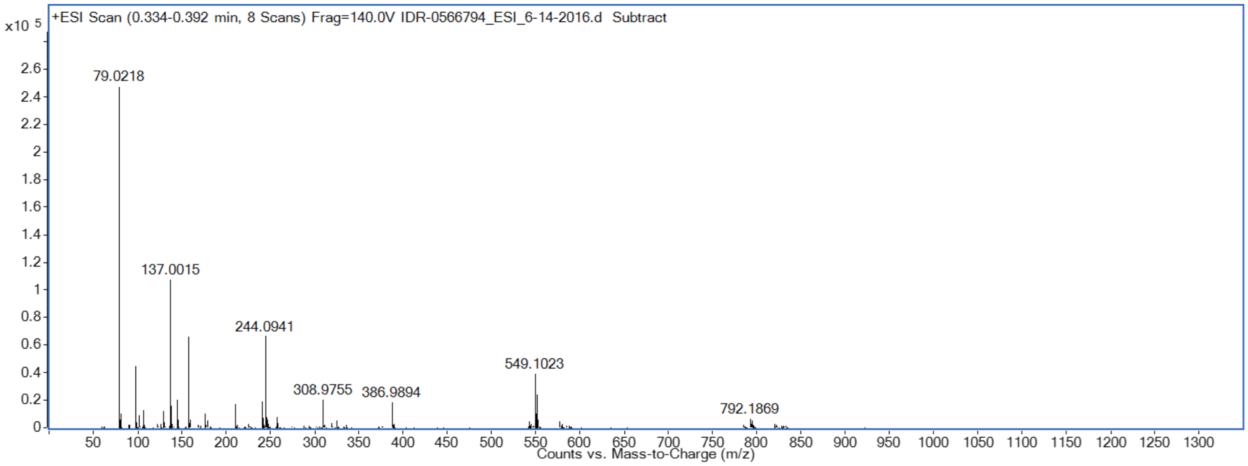

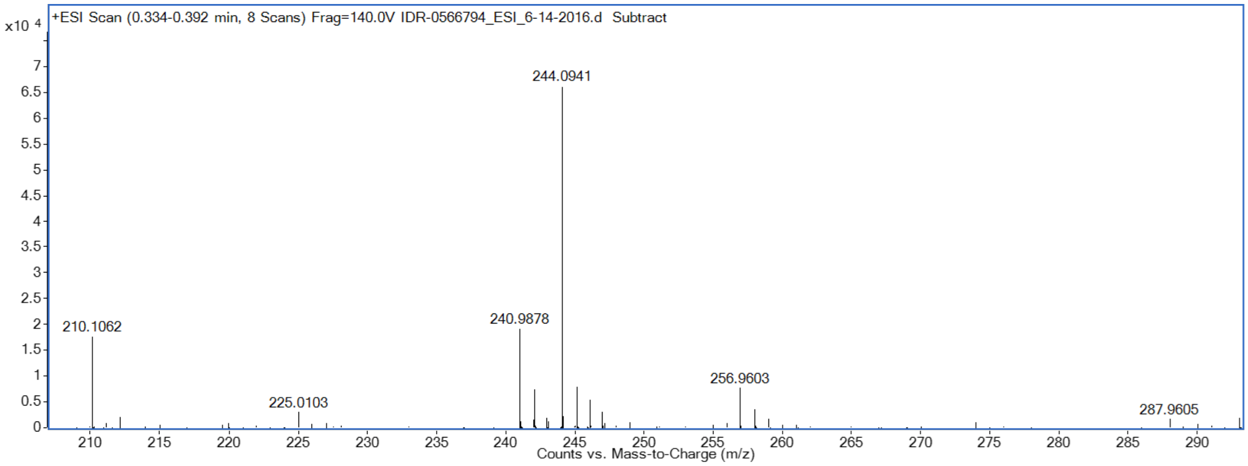


**(19) IDR-0532542/TPN-0002016**

$[M+H]^+$ calcd for $C_9H_{16}N_4S$: 213.1168

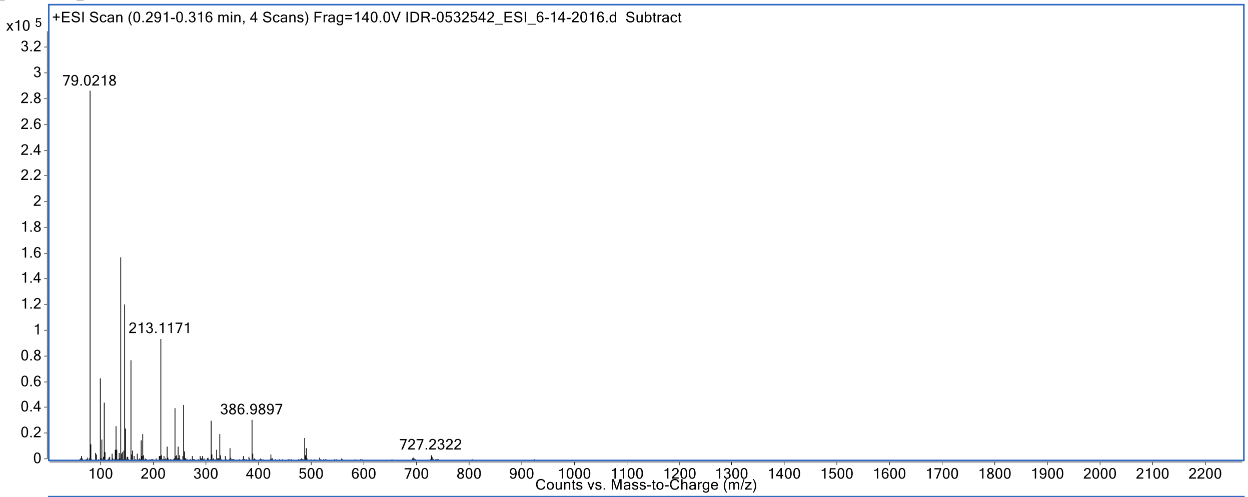


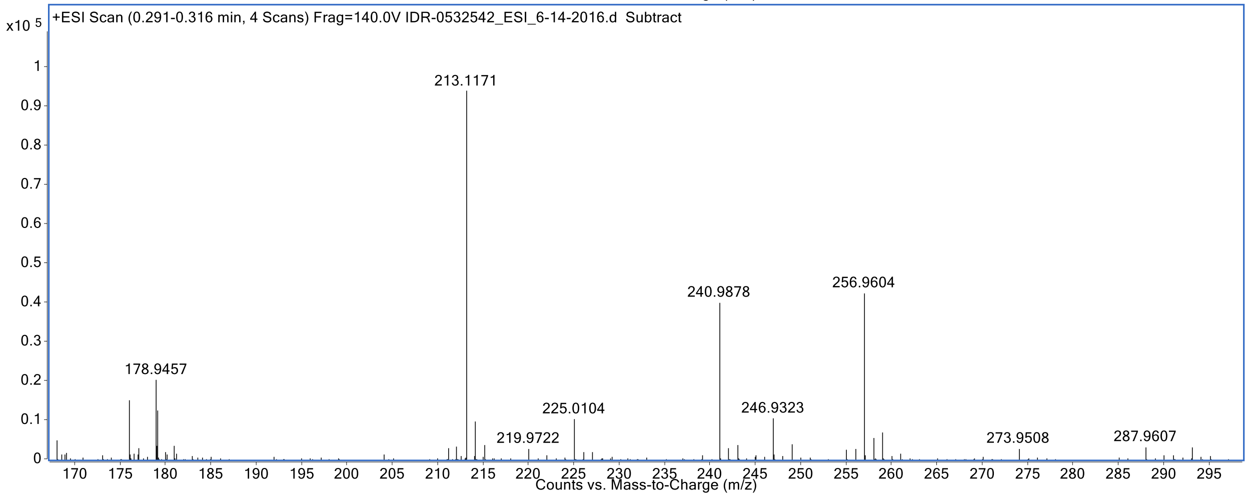


**(20) IDR-0571506/TPN-0002004**

$[M+H]^+$ calcd for C10H16N4S, 225.1180, found 225.1169

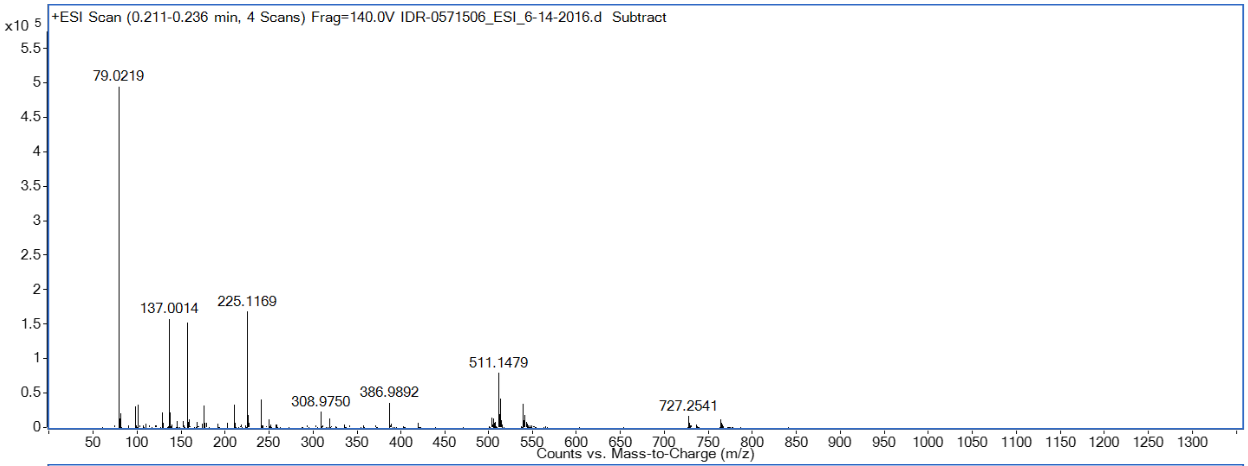


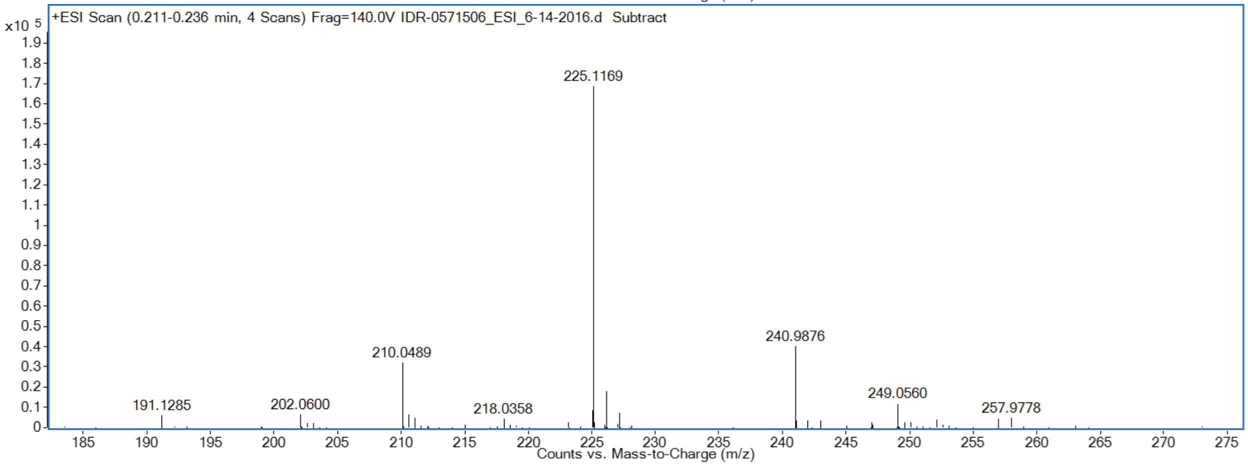


**(21) IDR-0571504/TPN-0002005**

$[M+H]^+$ calcd for $C_9H_{15}N_3OS$, 214.1020, found 214.1011

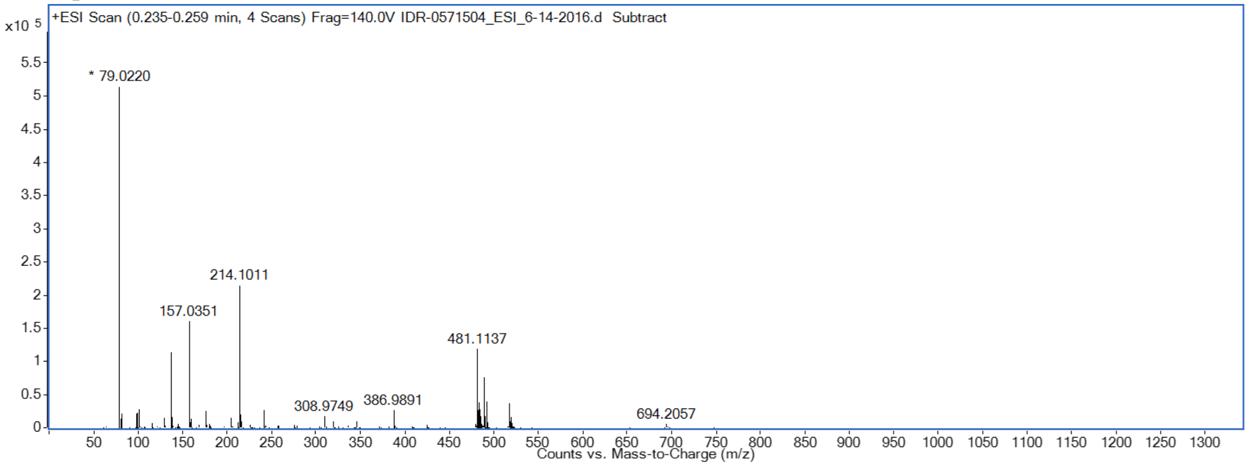

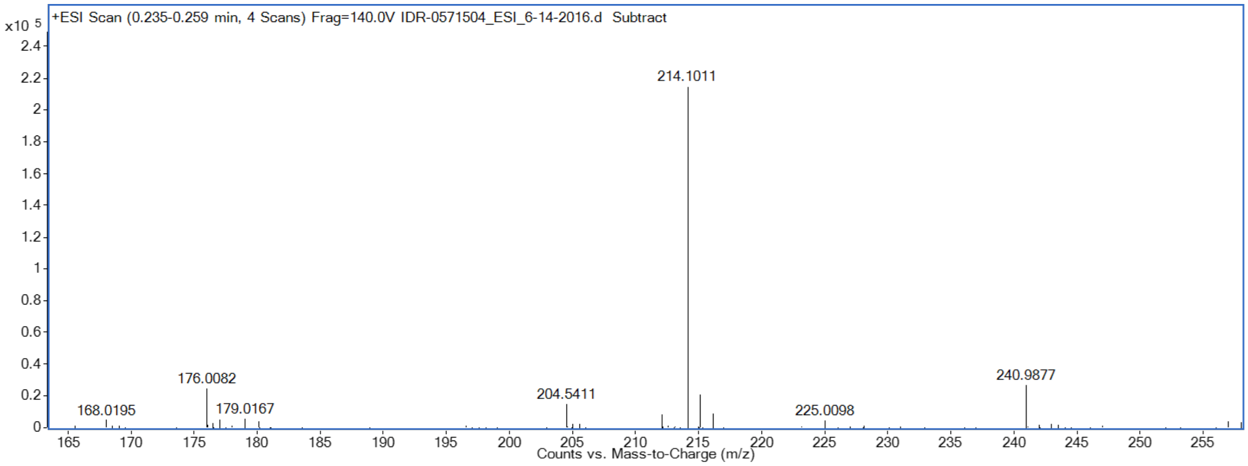


#### (22) IDR-0578471/TPN-0002000

$[M+H]^+$ calcd for $C_9H_{15}N_3OS$, 214.1020, found 214.1011

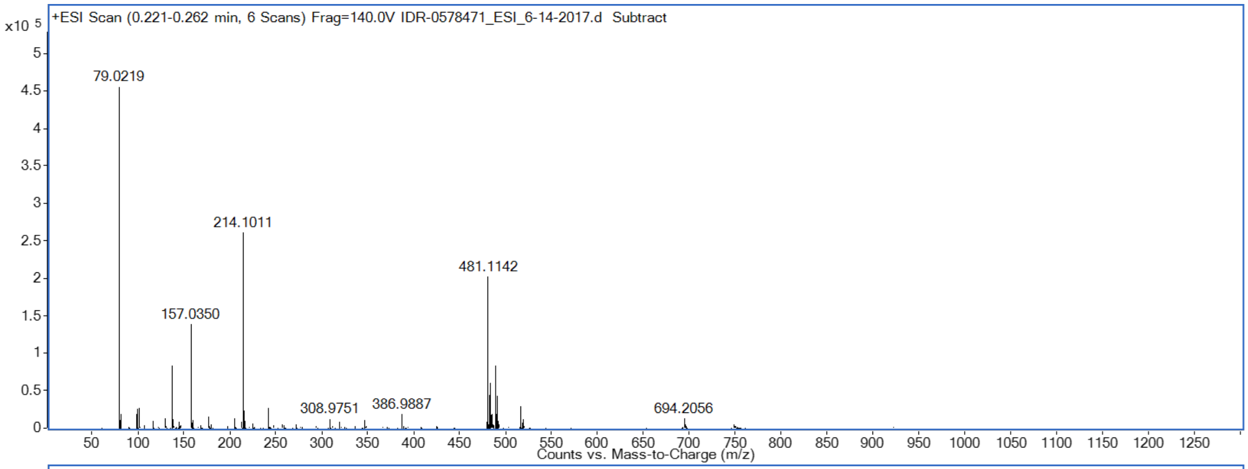


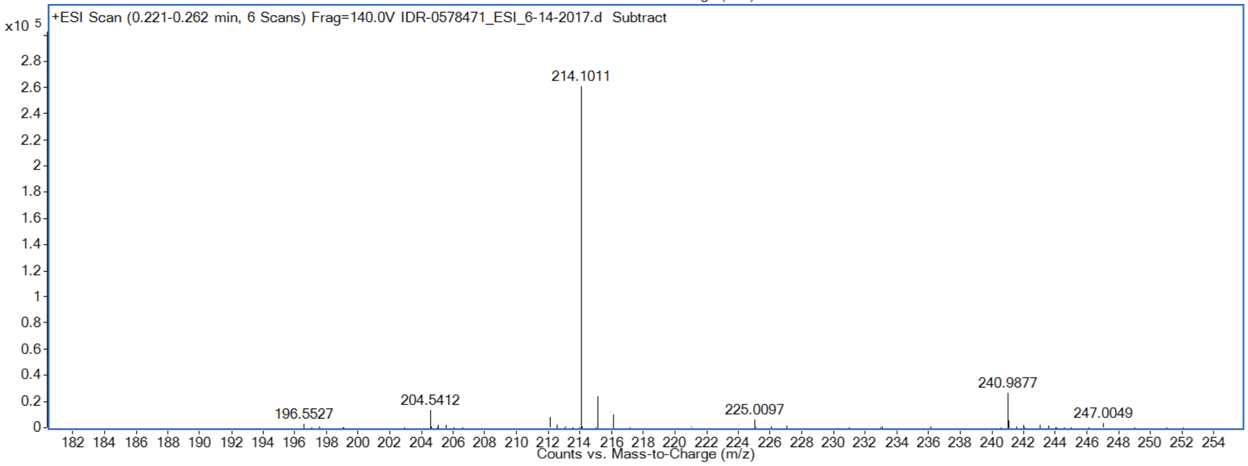


#### (23) IDR-0578334/TPN-0002001

$[M+H]^+$ calcd for $C_{14}H_{17}N_3OS$, 276.1170, found 276.1170

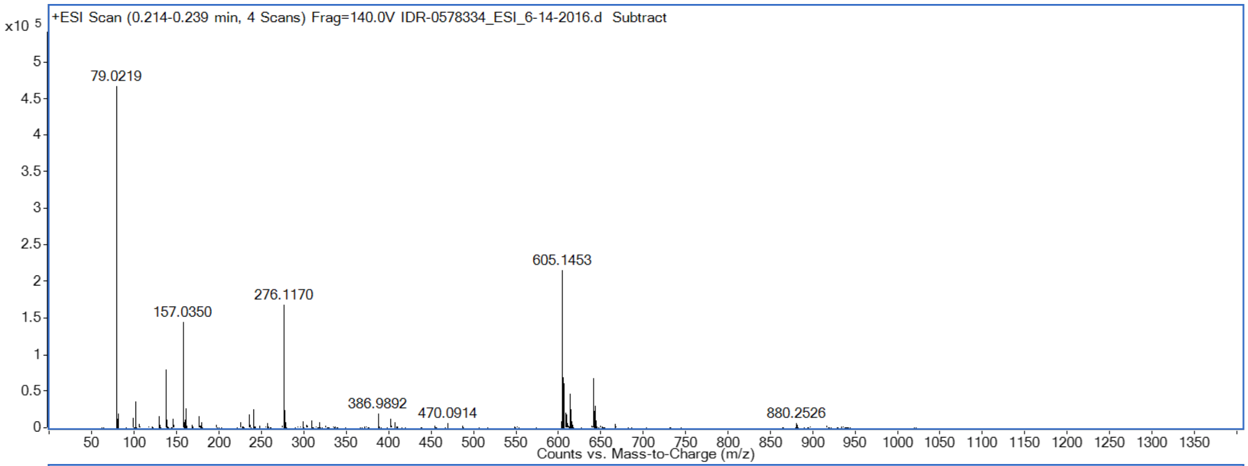


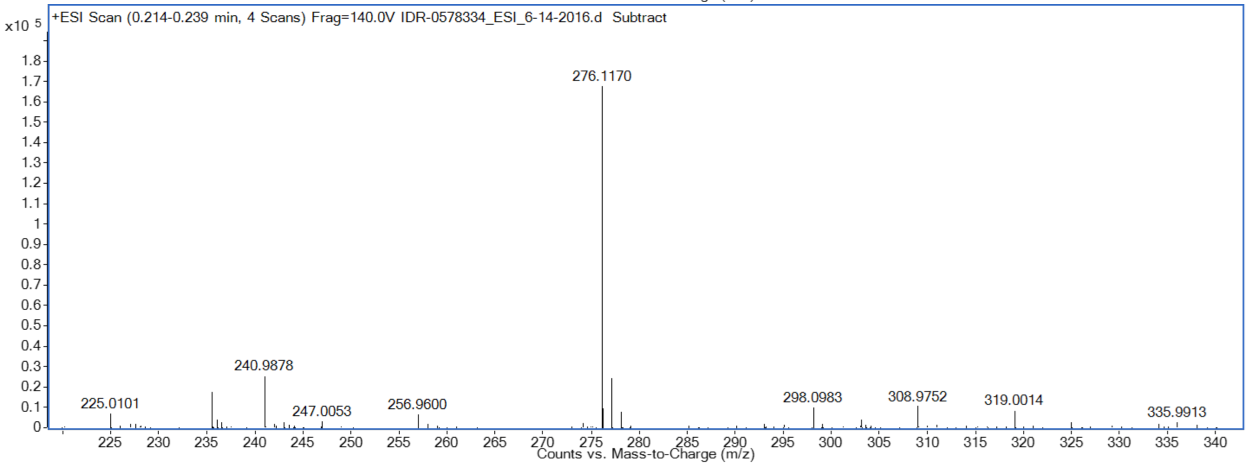


#### (24) IDR-0532359/TPN-0002026

$[M+H]^+$ calcd for $C_{12}H_{15}N_3OS_2$: 282.0729

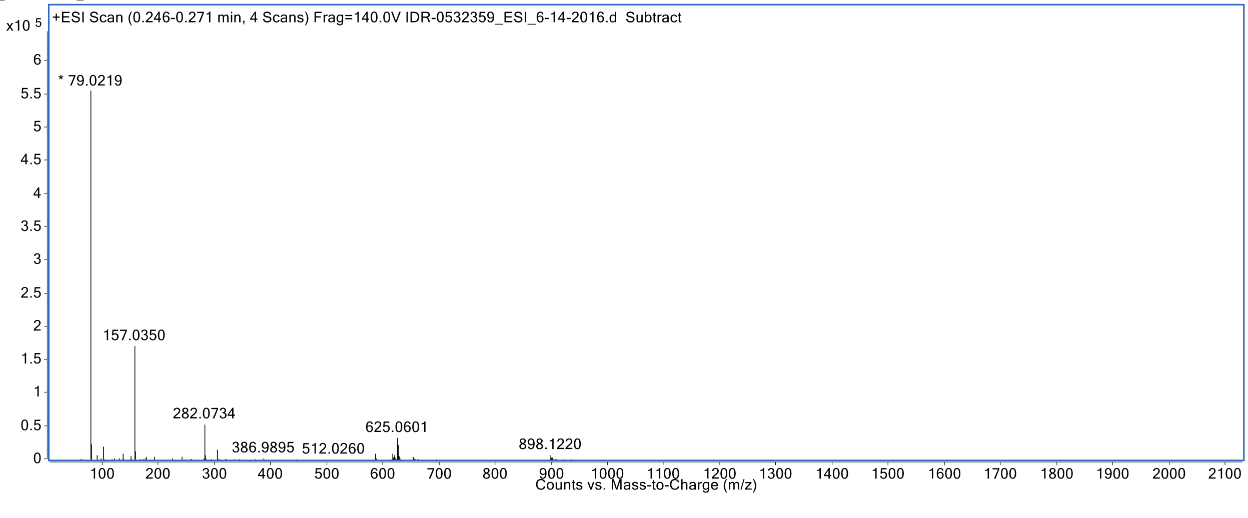

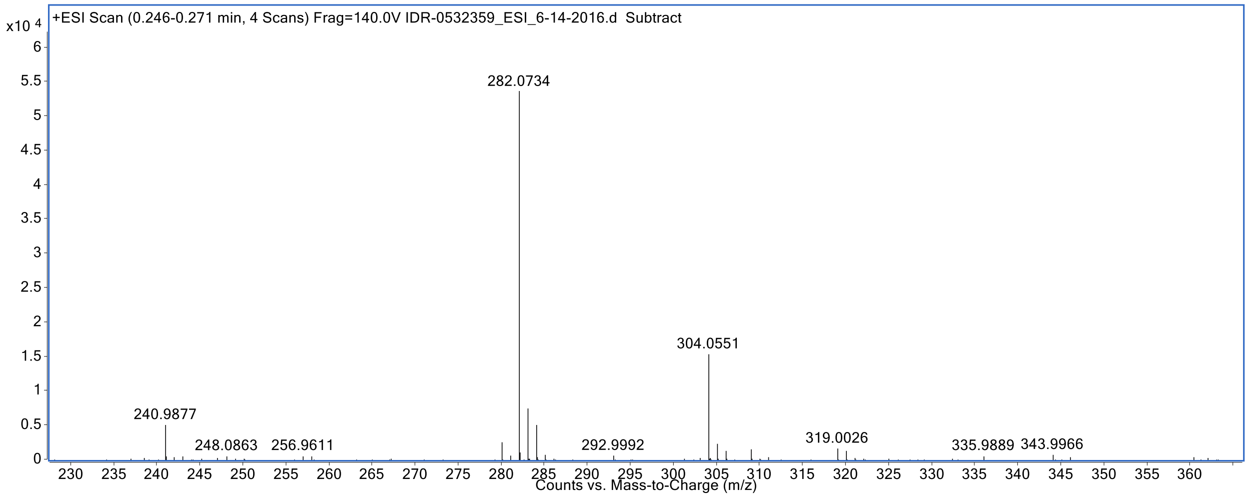


#### (25) IDR-0563553/TPN-0002014

$[M+H]^+$ calcd for $C_{12}H_{14}N_2S_3$, 283.0400, found 283.0397

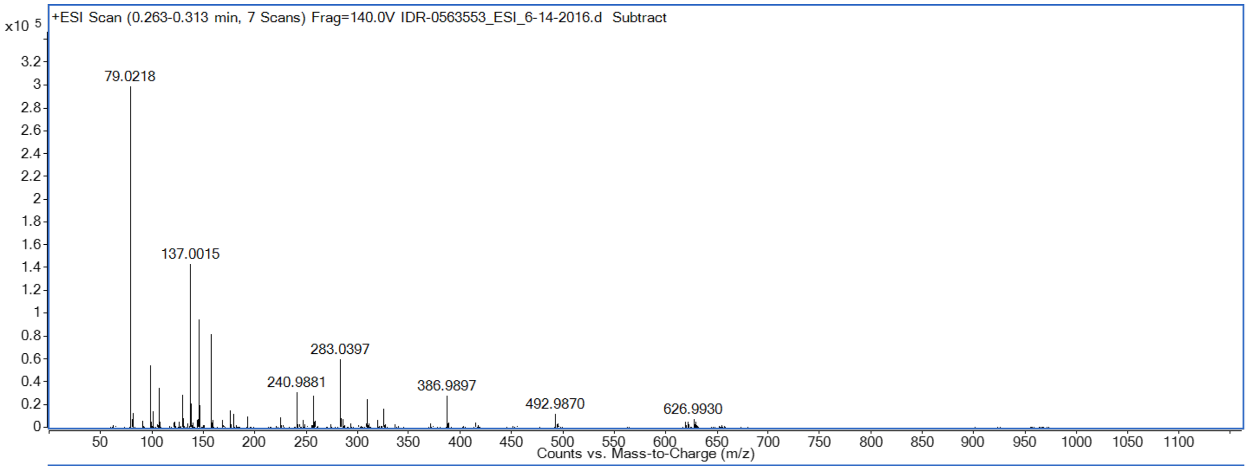


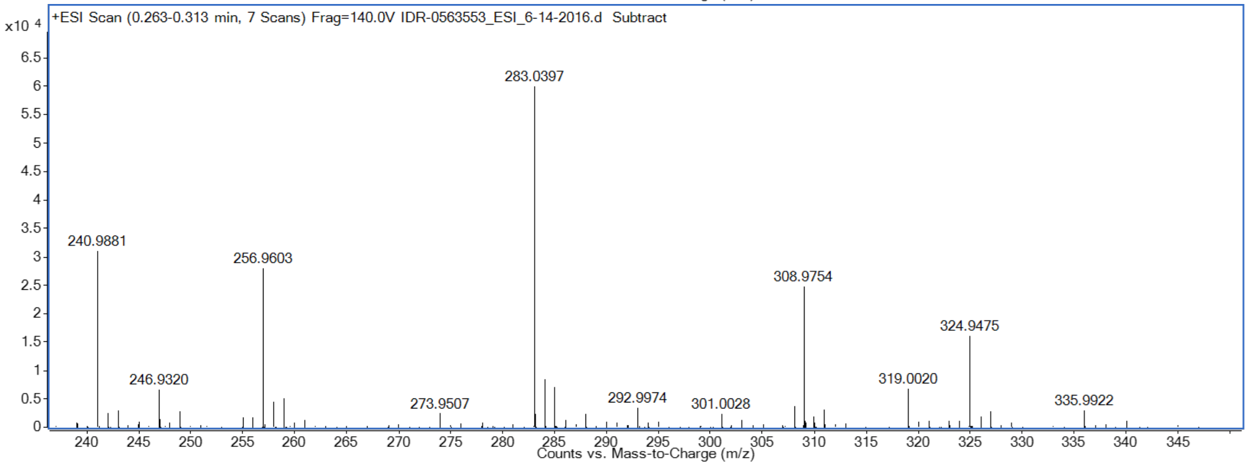


#### (26) IDR-0532360/TPN-0002025

$[M+H]^+$ calcd for $C_{16}H_{15}N_3S_2$: 314.0780

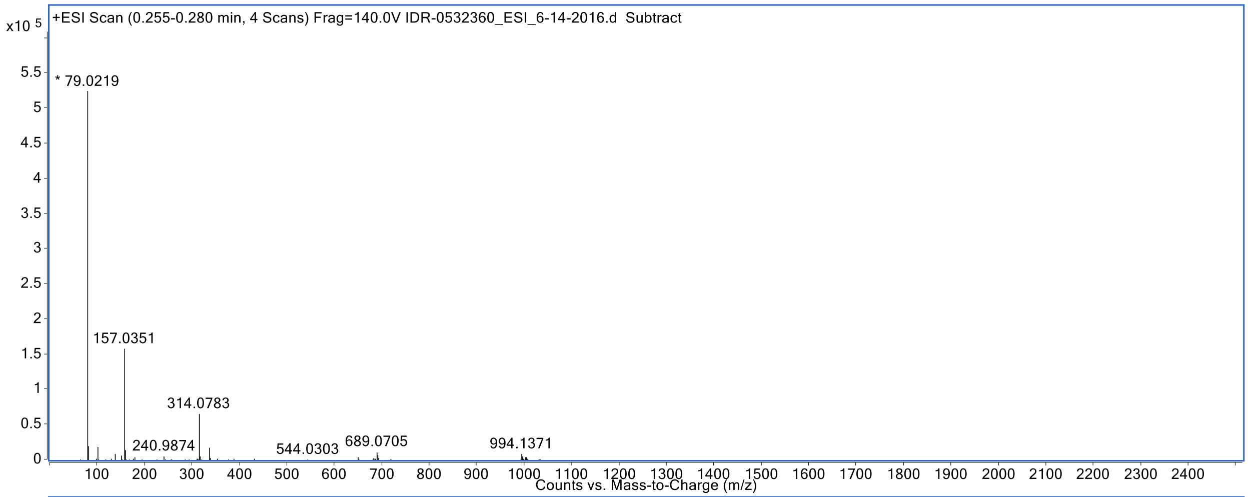


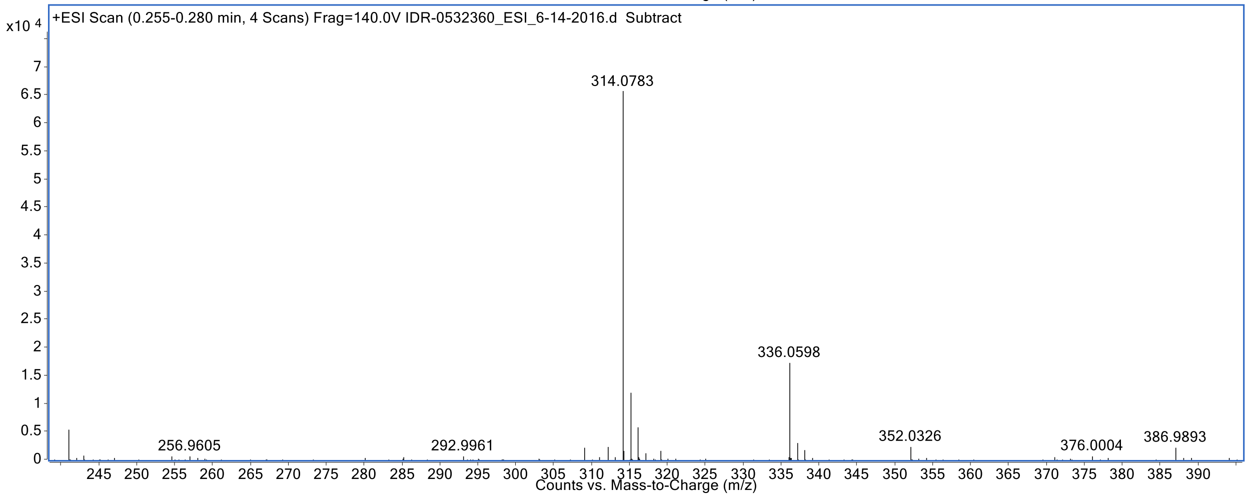


#### (28) IDR-0532392/TPN-0002024

$[M+H]^+$ calcd for $C_{12}H_{15}N_3S_2$: 266.0780

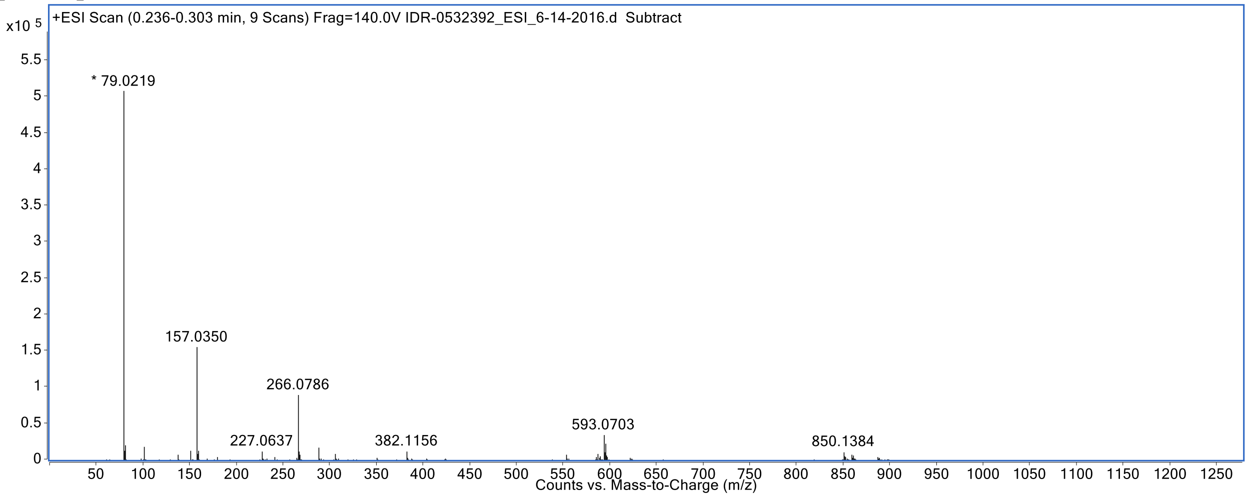

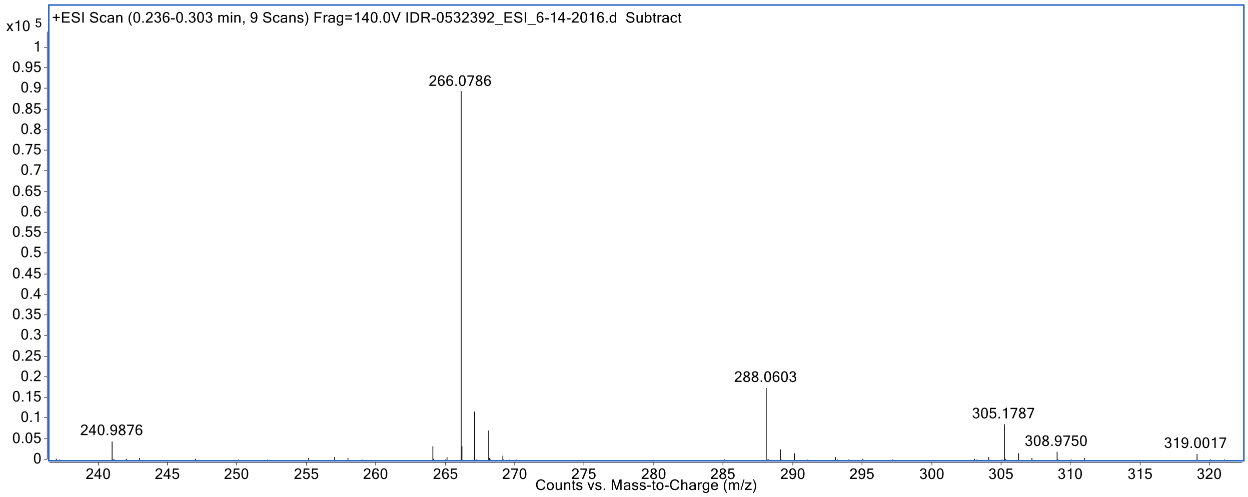


#### (29) IDR-0578333/TPN-0002002

[M+H]+ calcd for $C_9H_{15}N_3OS$, 214.1020, found 214.1007

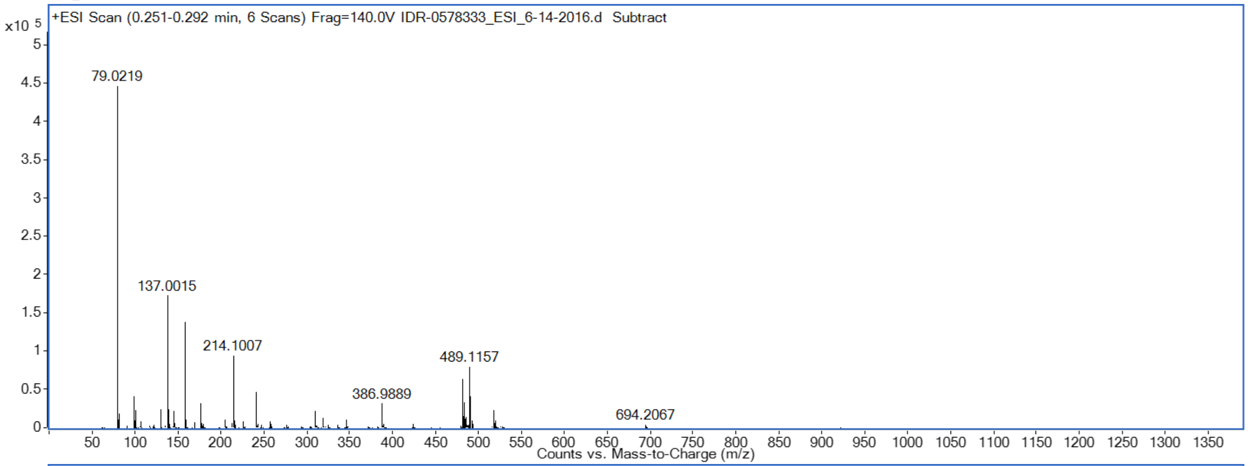


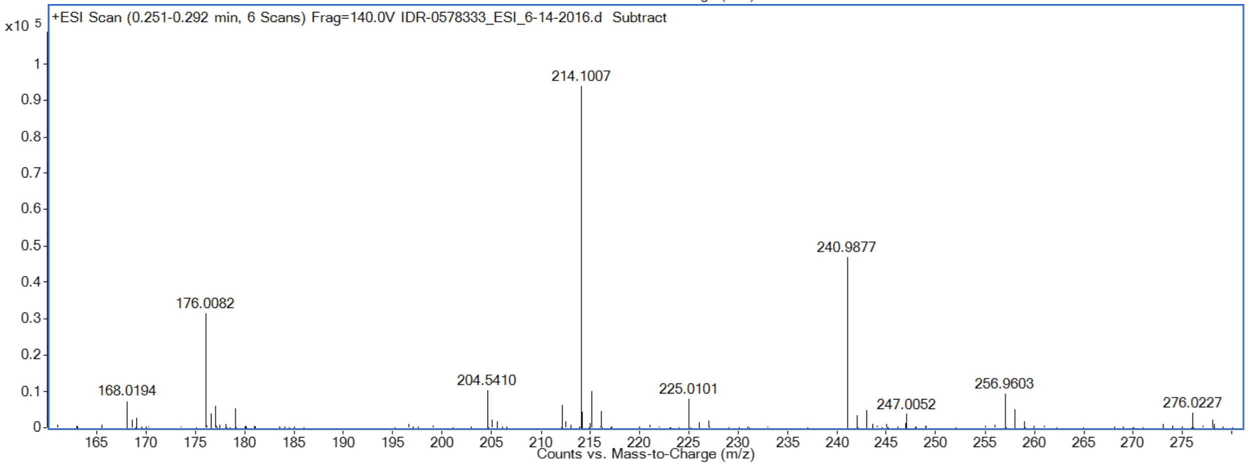


#### (30) IDR-0571381/TPN-0002010

[M+H]+ calcd for $C_{12}H_{15}N_3S_2$, 266.0790, found 266.0786

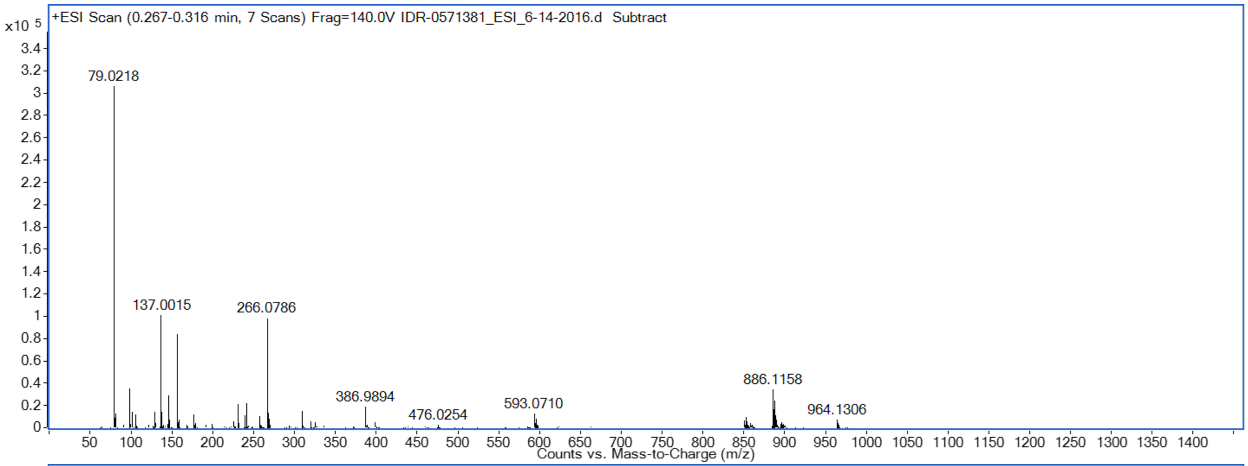


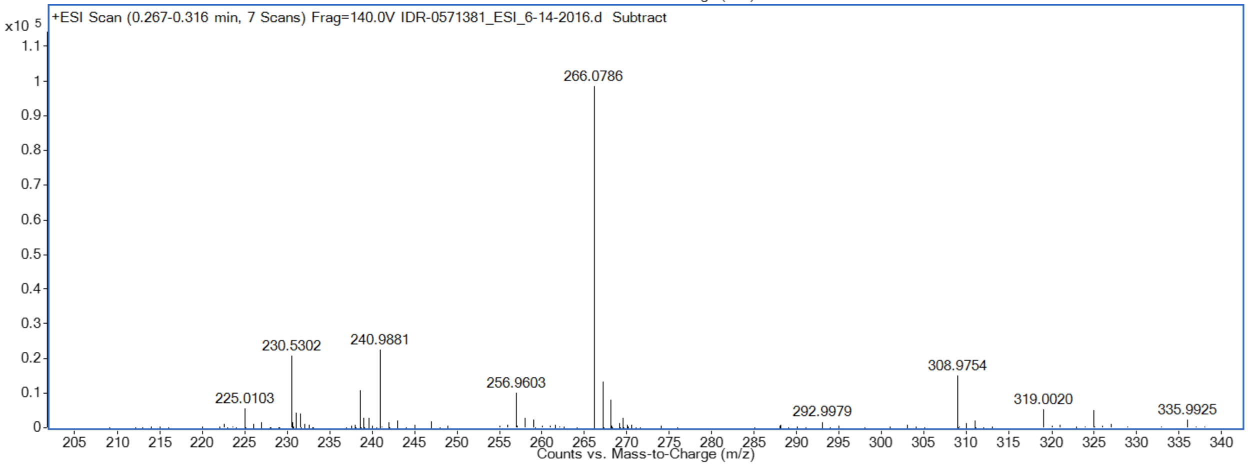


#### (31) IDR-0532393/TPN-0002023

$[M+H]^+$ calcd for $C_{11}H_{12}ClN_3S_2$: 286.0234

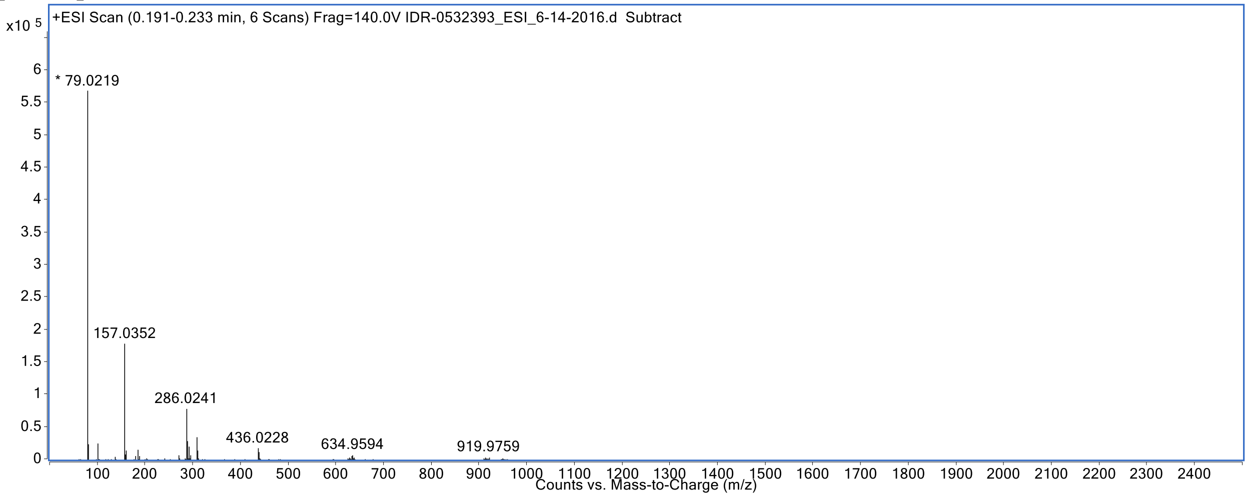

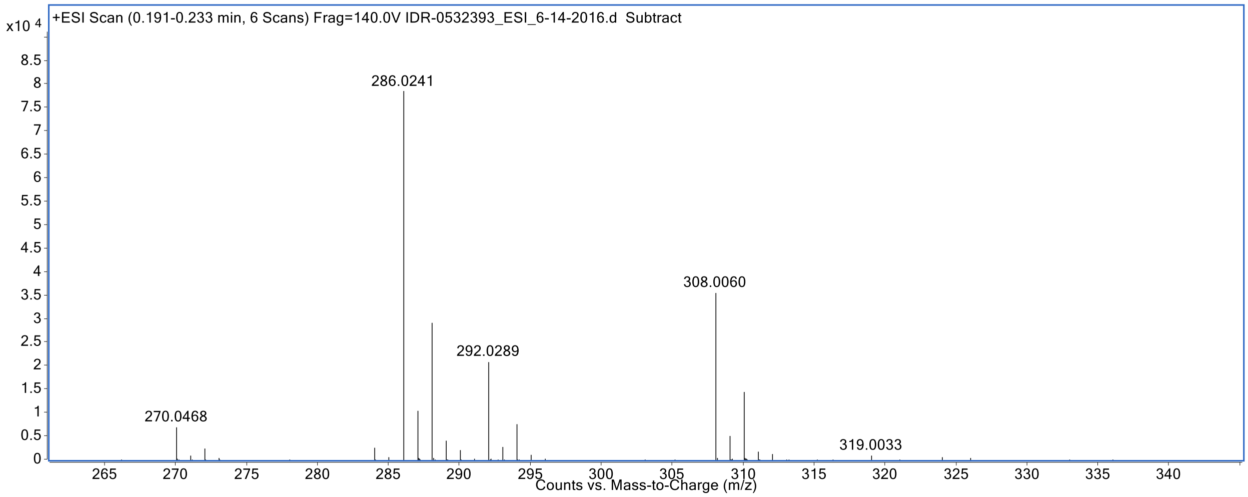


**(32) IDR-0571501/TPN-0002008**

$[M+H]^+$ calcd for $C_{12}H_{14}ClN_3OS$, 284.0630, found 284.0622

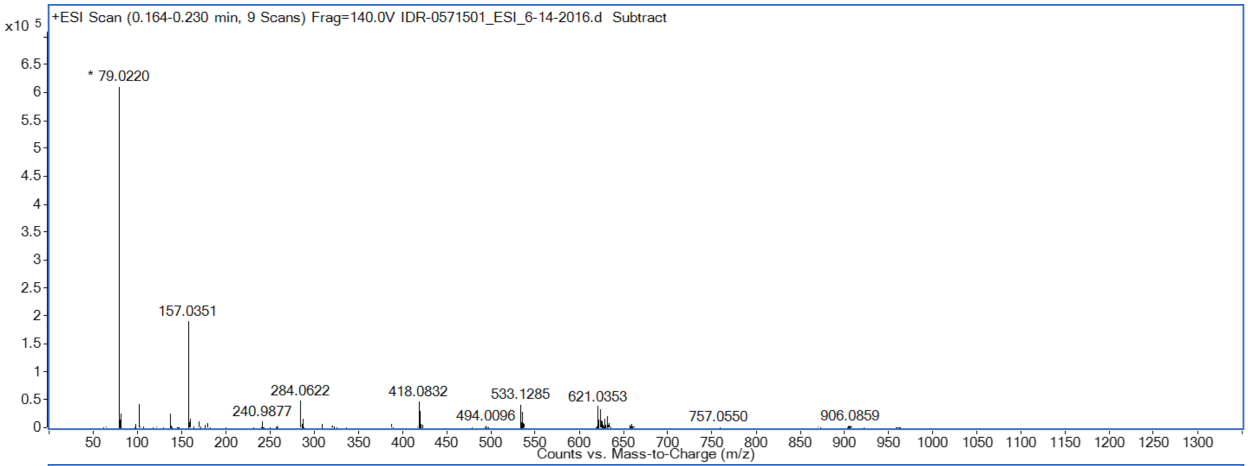


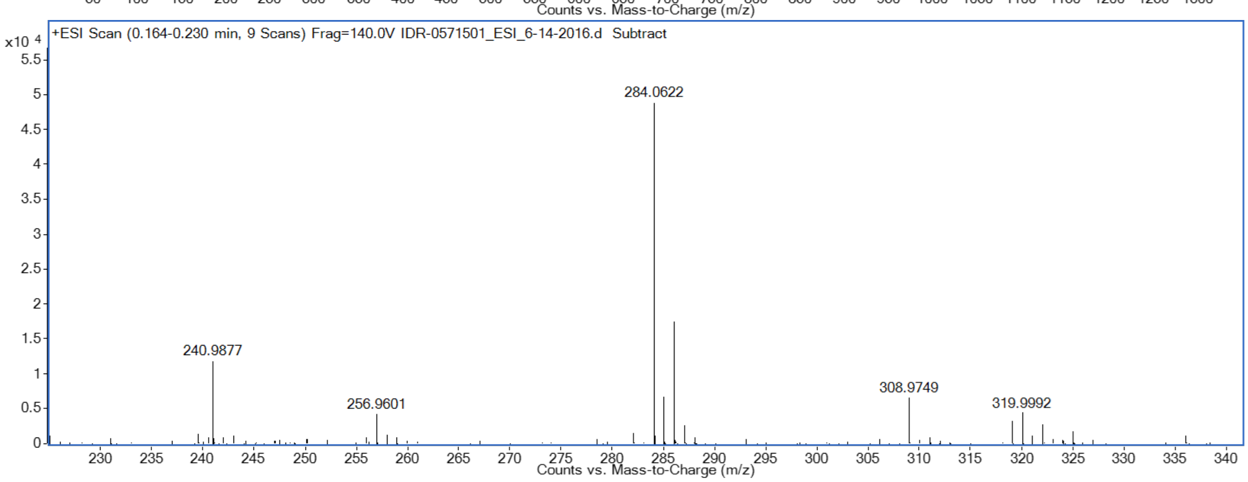


**(33) IDR-0571502/TPN-0002007**

$[M+H]^+$ calcd for $C_{12}H_{14}ClN_3OS$, 284.0630, found 284.0621

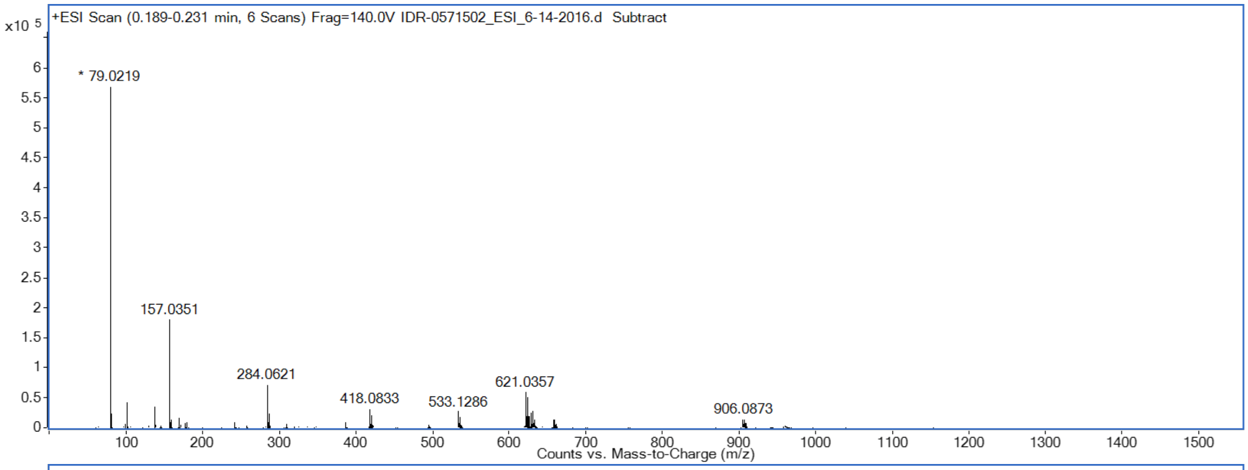


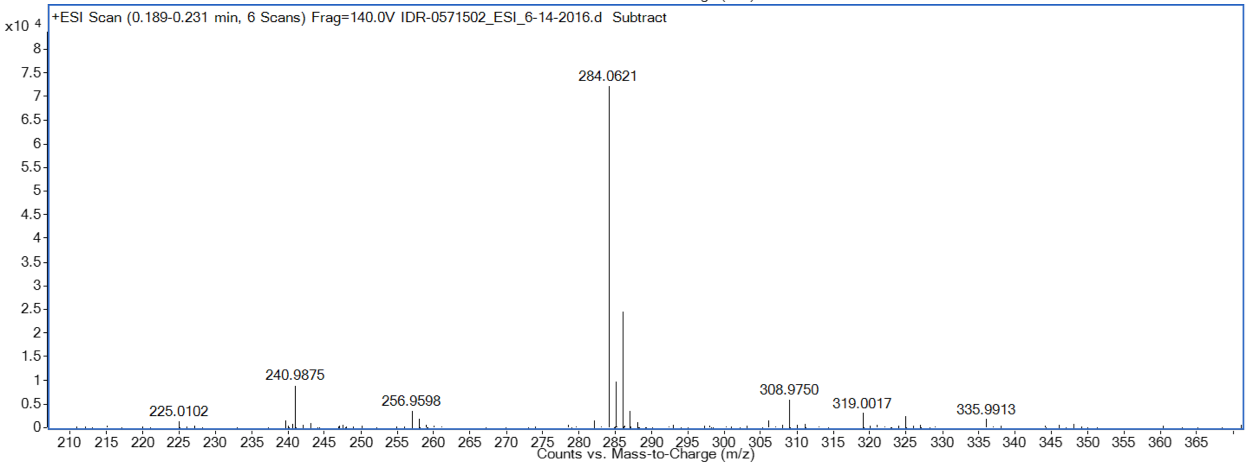


#### (34) IDR-0571503/TPN-0002006

$[M+H]^+$ calcd for $C_{12}H_{14}ClN_3OS$, 284.0630, found 284.0621

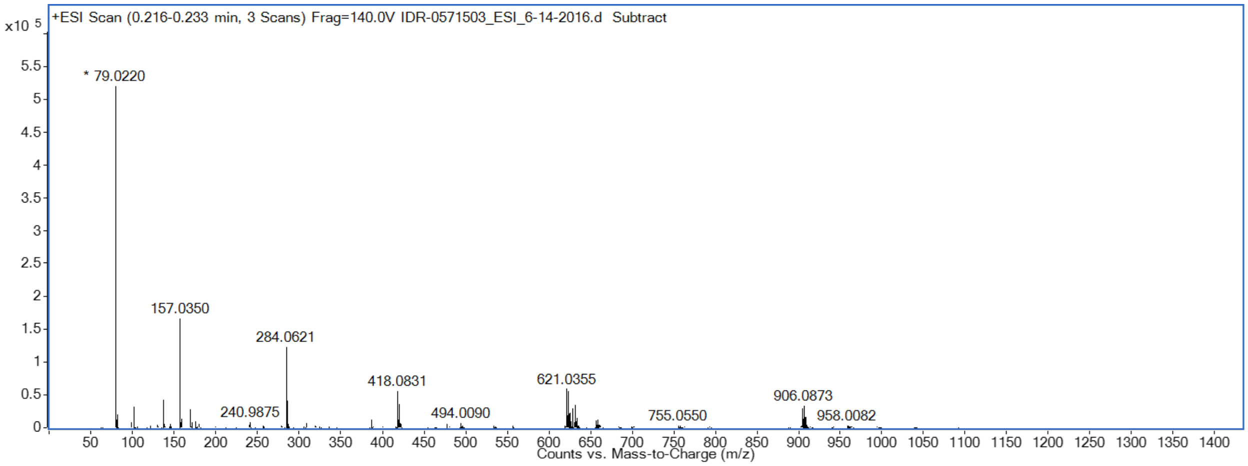

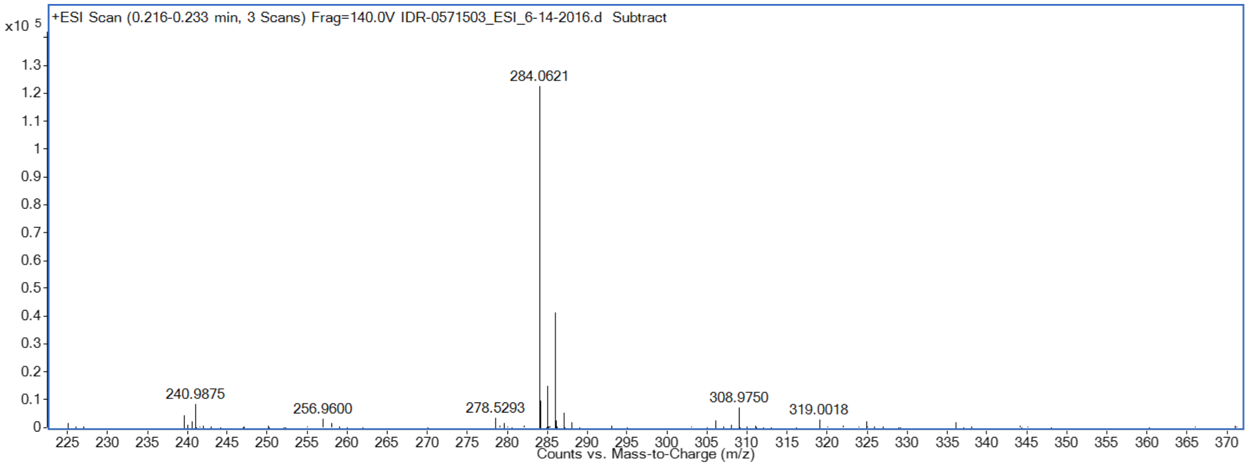


#### (35) IDR-0576402/TPN-0002003

$[M+H]^+$ calcd for $C_{13}H_{17}N_3O_2S$, 280.1120, found 280.1118

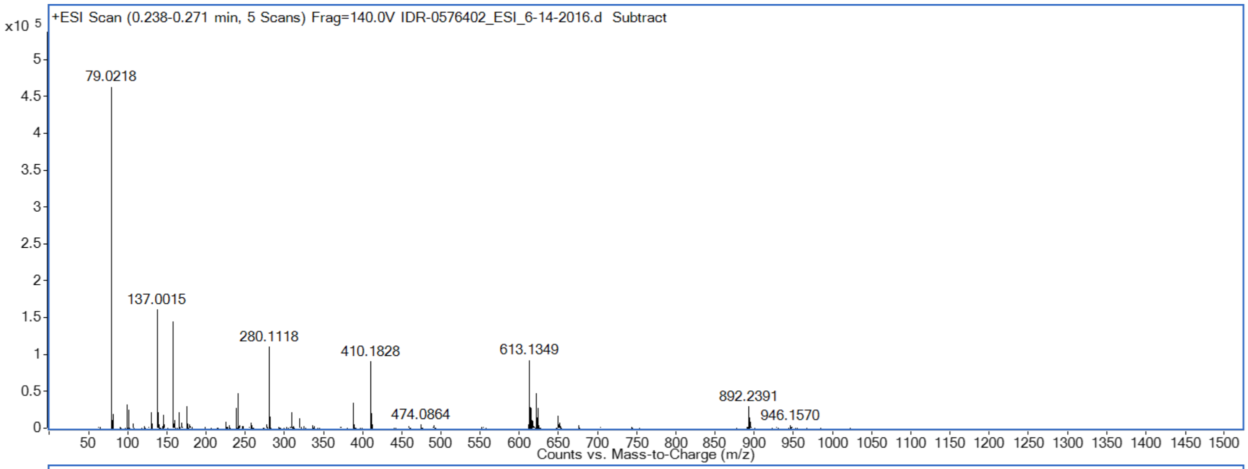


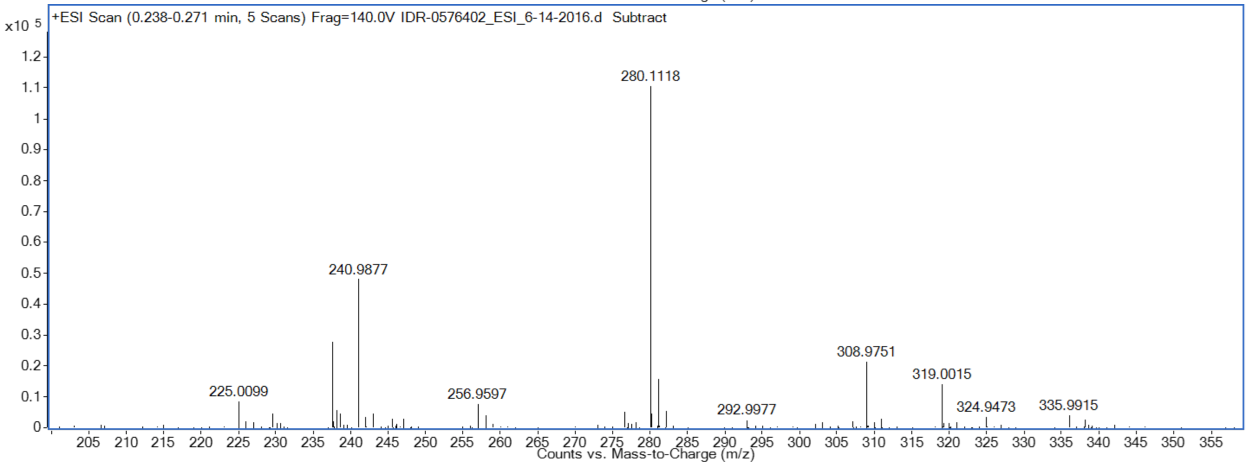
